# Context-dependent Interactors Regulate TDP-43 Dysfunction in ALS/FTLD

**DOI:** 10.1101/2025.04.07.646890

**Authors:** Longxin Xie, Yuehua Zhu, Bryan T. Hurtle, Matthew Wright, John L. Robinson, Jocelyn C. Mauna, Emily E. Brown, Marilyn Ngo, Cristian A. Bergmann, Jiazhen Xu, Jessica Merjane, Amanda M. Gleixner, Gabriela Grigorean, Feilin Liu, Wilfried Rossoll, Edward B. Lee, Evangelos Kiskinis, Maria Chikina, Christopher J Donnelly

**Affiliations:** Department of Neurobiology, University of Pittsburgh, Pittsburgh, PA, USA; LiveLikeLou Center for ALS Research, University of Pittsburgh, Pittsburgh, PA, USA; Department of Computational and Systems Biology, University of Pittsburgh, Pittsburgh, PA, USA; School of Medicine, Tsinghua University, Beijing, China; The Ken & Ruth Davee Department of Neurology, Feinberg School of Medicine, Northwestern University,, Chicago, IL, USA; Center for Neurodegenerative Disease Research, Department of Pathology and Laboratory Medicine, Institute on Aging, Perelman School of Medicine, University of Pennsylvania, Philadelphia, PA, USA; Integrated Systems Biology PhD Program, University of Pittsburgh, Pittsburgh, PA, USA; Human Genetics PhD Program, University of Pittsburgh, Pittsburgh, PA; Interdisciplinary Biomedical PhD Program, Molecular Pathology, University of Pittsburgh, Pittsburgh, PA, USA; Department of Proteomics, University of California, Davis, CA, USA; Department of Neuroscience, Mayo Clinic, Jacksonville, FL, USA

## Abstract

TDP-43 mislocalization, aggregation, and loss of splicing function are neuropathological hallmarks in over 97% of Amyotrophic Lateral Sclerosis (ALS), 45% of Frontotemporal Lobar Degeneration (FTLD), and 60% of Alzheimer’s Disease, which has been reclassified as LATE-NC. However, the mechanisms underlying TDP-43 dysfunction remain elusive. Here, we utilize APEX2-driven proximity labeling and mass spectrometry to characterize the context-dependent TDP-43 interactome in conditions of cytoplasmic mislocalization, impaired RNA-binding contributing to aggregation, and oxidative stress. We describe context-dependent interactors, including disrupted interactions with splicing-related proteins and altered biomolecular condensate (BMC) associations. By integrating ALS and FTLD snRNA-seq data, we uncover disease-relevant molecular alterations and validate our dataset through a functional screen that identifies key TDP- 43 regulators. We demonstrate that disrupting nuclear speckle integrity, particularly through the downregulation of the splicing factor SRRM2, promotes TDP-43 mislocalization and loss of function. Additionally, we identify NUFIP2 as an interactor associated with mislocalization that sequesters TDP-43 into cytoplasmic aggregates and co-localizes with TDP-43 pathology in patient tissue. We also highlight HNRNPC as a potent TDP-43 splicing regulator, where precise modulation of TDP-43 or HNRNPC can rescue cryptic exon splicing. These findings provide mechanistic insights and potential therapeutic targets for TDP-43 dysfunction.

## Introduction

Amyotrophic lateral sclerosis (ALS) and frontotemporal lobar degeneration (FTLD) are progressive, age-related neurodegenerative diseases manifesting in the degeneration of motor neurons in the brain and spinal cord or neurons in the frontal and temporal lobes, respectively ^1,2^. While certain ALS and FTLD cases are directly linked to genetic mutations, such as the C9orf72 repeat expansion (reviewed by ^3^), the majority arise sporadically, including ∼85-90% of ALS ^4^ and ∼60% of FTLD cases ^5^. In addition to their genetic and clinical overlap, ALS and FTLD share a common neuropathological feature: TDP-43 mislocalization and aggregation, present in ∼97% of ALS cases and ∼ 45% of FTLD cases. ^6,7^. TDP-43 is an RNA-binding protein that plays a critical role in RNA metabolism, including splicing, transport, and stability^8,9^; however, the mechanisms driving its pathological mislocalization, oligomerization, and splicing dysfunction remain unclear.

Several lines of research in iPSC-derived human and mouse models show that TDP-43 dysfunction has dual mechanisms of toxicity in ALS/FTLD, demonstrating both gain-of-function (GOF) toxicity and loss-of-function (LOF) events ^10^. Its cytoplasmic inclusions and cleavage products can induce toxicity^11–16^, yet partial depletion of TDP-43 also drives neurodegeneration, exemplified by the emergence of cryptic exons in essential transcripts (CFTR, STMN2, UNC13A, and HDGFL2) ^17–23^. Thus, TDP-43 pathobiology poses a significant therapeutic challenge: alleviating TDP-43 GOF must not compromise its critical nuclear roles ^24,25^. Different strategies targeting TDP-43 clearance^26–28^ or restoring individual splicing events^29,30^ show therapeutic promise but are complicated by the need to preserve TDP-43’s essential nuclear functions.

Mounting evidence suggests that TDP-43 pathology is characterized by dynamic shifts in its RNA or protein–protein interactome ^31^. TDP-43 can localize to various biomolecular condensates (BMCs), including stress granules, P-bodies, and nuclear pore complexes, each with the potential to facilitate normal function or accelerate disease progression depending on context^32–35^ (reviewed by ^36,37^). Several studies indicate that modulating TDP-43’s interacting partners, including short RNAs, HDAC6, HSP70, and HSPB1, may either promote or inhibit its liquid-liquid phase separation (LLPS), aggregation potential, cellular dynamics, and subsequent neuronal function toxicity^15,38–41^. Given the complexity of TDP-43 dysfunction in ALS/FTLD, defining the context- dependent TDP-43 interactome and the consequence on its physiological function is critical for elucidating disease mechanisms and identifying potential therapeutic targets^42^. APEX2 peroxidase-mediated proximity labeling offers a powerful tool to map the local proteome surrounding TDP-43, providing a global perspective on its interactome under diverse cellular conditions^38,39,43,44^. Though prior studies have explored TDP-43 pathology in genetic models (e.g. drosophila, mice and iPSCs)^45–48^ and identified its common interacting partners^34,38,39,42,49–51^, a systematic examination of TDP-43’s interactomes, alongside functional changes such as localization, RNA-binding status, and environmental stressors, has not been conducted.

To address this gap, we profiled the interactome of TDP-43 under various pathological conditions using APEX2-proximity labeling followed by mass spectrometry^52,53^. We focused on three drivers of TDP-43 dysfunction: cytoplasmic mislocalization, impaired RNA-binding, and oxidative stress, all of which induce TDP-43 LOF ^54^. We explored context-dependent lost or gained interactors, as well as altered TDP-43’s interactions with critical cellular BMCs under these conditions. We then systematically identified specific interactors that exacerbate or mitigate TDP-43 LOF, including key RNA-binding proteins. We further integrate our findings with single-nucleus RNA-sequencing (snRNA-seq) data from ALS/FTLD patient neurons^55^, providing direct evidence that context-dependent TDP-43 interactors identified in our cellular system mirror dysregulated pathways in human disease.

This study expands on a unifying principle in which TDP-43 pathology arises from context-specific “interactome switches” that reshape its functionality. By identifying key regulators that maintain normal TDP-43 status (e.g. SRRM2 and nuclear speckles), drive its mislocalization (e.g. NUFIP2) or buffer its splicing LOF (e.g. HNRNPC), we highlight novel mechanistic pathways and potential therapeutic targets for ALS/FTLD and related disorders. This approach underscores the importance of charting the local proteome of TDP-43 across diverse disease-relevant conditions to understand the dynamic nature of upstream mechanisms driving TDP-43 dysfunction and, ultimately, to develop precision strategies^54,56^ for restoring its normal nuclear functions without inducing gain-of-function toxicity. Beyond novel discoveries, our study provides a comprehensive and context-specific TDP-43 interactome dataset, serving as a valuable resource for the field to explore mechanisms of TDP-43 dysfunction and identify potential therapeutic targets.

## Results

### Cellular mislocalization, RNA-binding deficiencies, and extracellular cell stressors induce TDP-43 loss of function

To identify interactors that modulate TDP-43 function, we utilized the CFTR-UNC13A TDP-43 Sensor (CUTS)^54^, a biosensor that monitors TDP-43 splicing activity through GFP expression upon the aberrant inclusion of a cryptic exon (Figure 1A). We examined various triggers of TDP- 43 loss of splicing function (LOF), including knockdown ^17,57^, localization mutants, cell stressors, and RNA-binding perturbations (Extended Figure 1A, Supplementary Table 2). CUTS-HEK293 cell line treated with siRNA-mediated TDP-43 depletion significantly increased GFP fluorescence compared to control siRNA, confirming the CUTS biosensor’s sensitivity to TDP-43 LOF (Extended Figure 1B). Overexpression of mislocalization and aggregation-prone TDP-43 variants in CUTS-HEK293 cells, including cytoplasmic and RNA-binding-deficient mutants (ΔNLS, 5FL, 5FL/ΔNLS) also elevated CUTS-GFP fluorescence, suggesting impaired splicing activity of endogenous TDP-43 (Figure 1B, Extended Figure 1C)^54^. Similarly, transfection of these mutant variants (ΔNLS, 5FL, 5FL/ΔNLS) did not fully restore TDP-43 function) in TARDBP-/- HeLa cells^58^, as assessed by comparing the GFP signal intensity to TDP-43 rescue (Figure 1C, Extended Figure 1D^15,59^. This demonstrates varying degrees of splicing impairment among these dysfunctional variants. CUTS-GFP fluorescence increased in a dose-dependent manner upon treatment with cellular stressors known to disrupt TDP-43 localization, including oxidative stress (NaAsO2) ^32,60,61^, hyperosmotic pressure (sorbitol)^62^, translation inhibition (puromycin)^63^, and proteasome inhibition (MG132)^64^, highlighting their pathological impact on TDP-43 splicing function (Figure 1D, Extended Figure 1E). However, we did not detect increased LOF by membrane potential disruption (KCl)^65,66^ or ER stress (Tunicamycin)^67^ in the CUTS-HEK93 system (Figure 1D). NaAsO2 (AS) triggered the highest level of LOF among all the stressors, which is consistent with recent studies ^68^ likely due to the rapid formation of splicing-deficient AS-induced nuclear TDP-43 condensates. Additionally, consistent with recent findings, we observed significant TDP-43 LOF in the CUTS system by multivalent UG-rich RNA oligos ([UG]17; Extended Figure 1F-1G)^69^.

**Figure 1:**
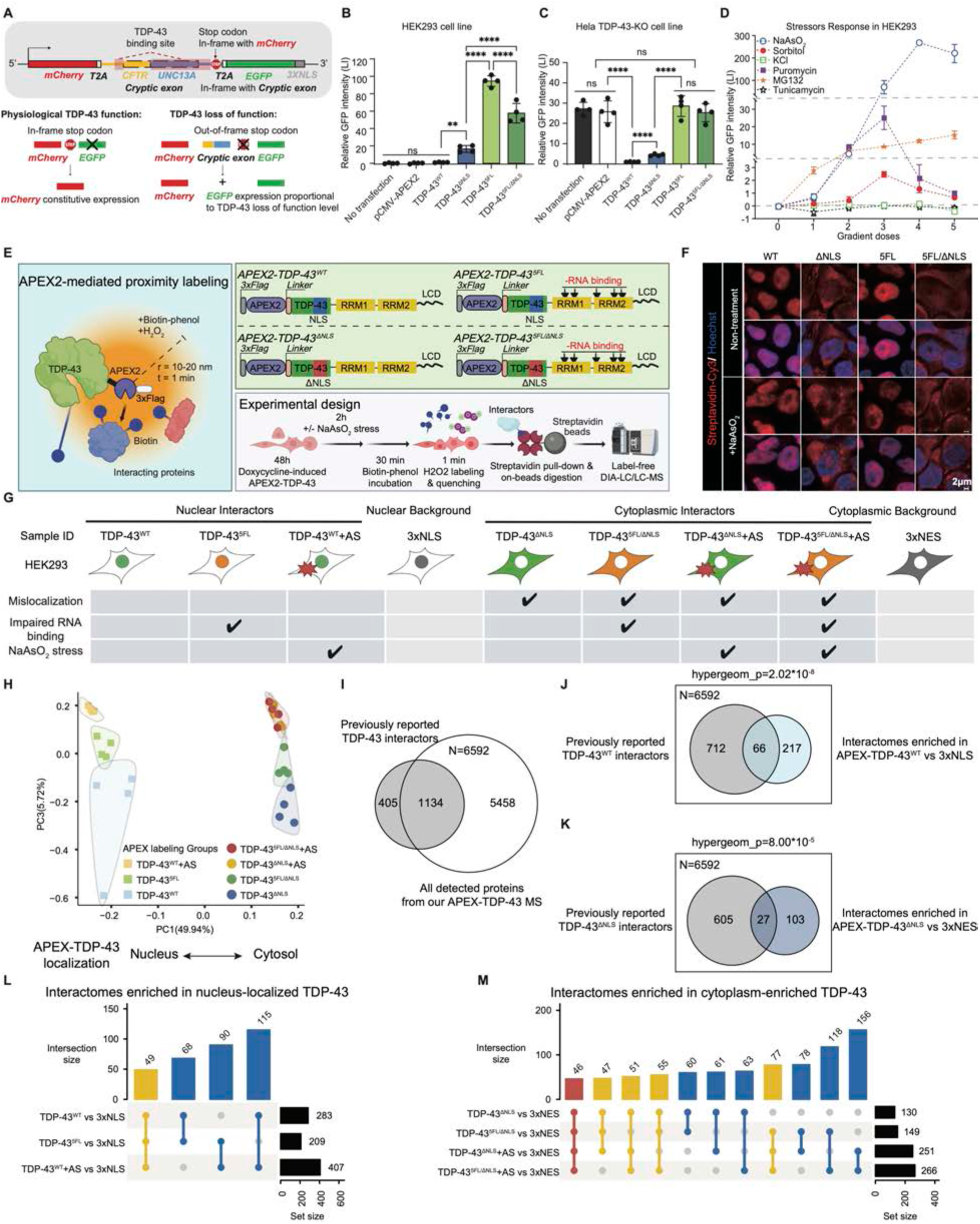
APEX2 proximity labeling and DIA mass spectrometry enable quantitative comparison of context-dependent TDP-43’s interactors. **(A)** Schematic diagram of the CUTS (CFTR-UNC13A TDP-43 Sensor) biosensor. CUTS biosensor was designed to detect TDP-43 loss of function in splicing by expressing nucleus GFP signal (EGFP-3XNLS) proportional to the level of TDP-43 LOF in real-time. **(B)** Relative GFP fluorescence intensity quantification from live confocal imaging in stable HEK293 cells expressing doxycycline-inducible (72 h of 1 mg/mL doxycycline) CUTS with the transfection of pCMV-APEX2 backbone-only, TDP-43^WT^, TDP-43^ΔNLS^, TDP-43^5FL^, TDP- 43^5FL/ΔNLS^ with pCMV-APEX2 backbone or non-transfected (72 h, N=3 biological replicates). **(C)** Relative GFP fluorescence intensity quantification from live confocal imaging in Hela TDP- 43-KO cells with the co-transfection of doxycycline-inducible (72h of 1 mg/mL doxycycline) CUTS plasmid and pCMV-APEX2 backbone-only, TDP-43^WT^, TDP-43^ΔNLS^, TDP-43^5FL^, TDP- 43^5FL/ΔNLS^ with pCMV-APEX2 backbone or no plasmid (72 h, N=3 biological replicates). For (B) and (C), statistical significance was determined by one-way ANOVA and Tukey’s multiple comparison test (* = P < 0.05; ** = P < 0.01; *** = P < 0.001; **** = P < 0.0001). Data are the mean ± s.d. **(D)** Relative GFP fluorescence intensity quantification from live confocal imaging in stable HEK293 cells expressing doxycycline-inducible (72 h of 1 mg/mL doxycycline) CUTS with the treatment of NaAsO2 (2 h, gradient doses: 0, 67.5, 125, 250, 500, 1000 μM), Sorbitol (4 h, gradient doses: 0, 100, 200, 400, 800, 1600 mM), KCl (4 h, gradient doses: 0, 12.5, 25, 50, 100, 200 mM), Puromycin (24 h, gradient doses: 0, 1.25, 2.5, 5, 10, 20 μg/mL), MG132 (24 h, gradient doses: 0, 5, 10, 20, 40, 80 μM), and Tunicamycin (24 h, gradient doses: 0, 5, 10, 20, 40, 80 μM) (N=2 biological replicates). Data are the mean ± s.d. **(E)** Schematic diagram of APEX2-TDP-43 system. HEK293 stable cell line with doxycycline- inducible TDP-43^WT^, TDP-43^ΔNLS^, TDP-43^5FL^, or TDP-43^5FL/ΔNLS^ were generated. BP and H2O2 were added sequentially to enable proximity labeling. **(F)** Representative immunofluorescence staining images of the biotin signal labeled by APEX2- TDP-43 variants in stable HEK293 cell lines (48 h of 1 mg/mL doxycycline) with or without NaAsO2 stress (250 μM, 2 h). Blue = Hoechst; Red = Streptavidin-Cy3. Scale bar = 2 µm. **(G)** Schematic diagram of the experimental design of the proximity labeling for data-independent acquisition (DIA) quantitative mass spectrometry with the combination settings of TDP-43 mislocalization, impaired RNA binding, and NaAsO2 stress. APEX2-3XNLS and APEX2-3XNES were included as compartment background controls. N=4 independent biological replicates for each one of the settings. **(H)** Principal component analysis (PCA) of the raw interactome lists from the seven experiment groups of APEX2-TDP-43 proximity labeling. PC1 (49.94% variance) and PC3 (5.72% variance) are shown as they distinguish different groups. **(I)** Venn diagram of the proteins detected and quantified by all the groups of APEX2-TDP-43 in this study with the combination of previously reported TDP-43 interactors. **(J)** Venn diagram of the proteins enriched in APEX2-TDP-43^WT^ group with previously reported TDP-43^WT^ (mainly in the nucleus) interactors. **(K)** Venn diagram of the proteins enriched in APEX2-TDP-43^ΔNLS^ group with previously reported TDP-43^ΔNLS^ (mainly in the cytoplasm) interactors. **(L)** Overlapping diagram of the enriched protein interactors among TDP-43^WT^, TDP-43^5FL^, and TDP-43^WT^+AS groups. **(M)** Overlapping diagram of the enriched protein interactors among TDP-43^ΔNLS^, TDP-43^5FL/ΔNLS^, TDP-43^ΔNLS^+AS, and TDP-43^5FL/ΔNLS^+AS groups. For (J) to (M), protein interactors from each group are enriched with log2FC > 0.3 and P < 0.05, comparing with their respective compartment controls (APEX2-3XNLS for the nucleus, APEX2- 3XNES for the cytoplasm).

### APEX2 Proximity Proteomics to Characterize the Context-Specific TDP-43 Interactome

To determine the protein interactome contributing to context-dependent TDP-43 LOF identified in (Figure 1B-D, Extended Figure 1), we employed APEX2-mediated proximity labeling. This system enables spatiotemporal biotinylation of protein interactors within ∼10–20 nm over a 1- minute window in live cells (Figure 1E)^43,70^. Both transiently expressed cDNA constructs or inducible and stably expressing HEK293 cell lines were generated using the *Piggybac* system^71^ with a Tet3G doxycycline-inducible cassette ^72,73^ expressing APEX2-TDP-43 variants (WT, ΔNLS, 5FL, and 5FL/ΔNLS) were generated to perform proximity labelling of TDP-43 interacting proteins (Figure 1E-1F, Extended Figure 2A-2C). While transient overexpression of ΔNLS, 5FL, and 5FL/ΔNLS variants formed nuclear or cytoplasmic aggregates as previously reported (Extended Figure 2B) ^15,31,38,39^, the doxycycline-inducible stable cell lines did not exhibit noticeable morphological changes (Figure 1F). Such differences might be caused by different protein concentrations, as these phase-separation structures were mostly found following overexpression. Consistent with this, APEX2-TDP-43 levels after transient transfection were higher than endogenous TDP-43 (Extended Figure 2D), whereas expression levels in the tunable doxycycline-inducible stable cell lines remained comparable to endogenous TDP-43 levels (Extended Figure 2E). Therefore, we prioritized the doxycycline-inducible stable cell lines as they better mimic physiological conditions since TDP-43 is not overexpressed in ALS/FTLD ^74,75^. This approach also allows for the identification of early TDP-43 protein interactors that may trigger insoluble condensate formation.

**Figure 2:**
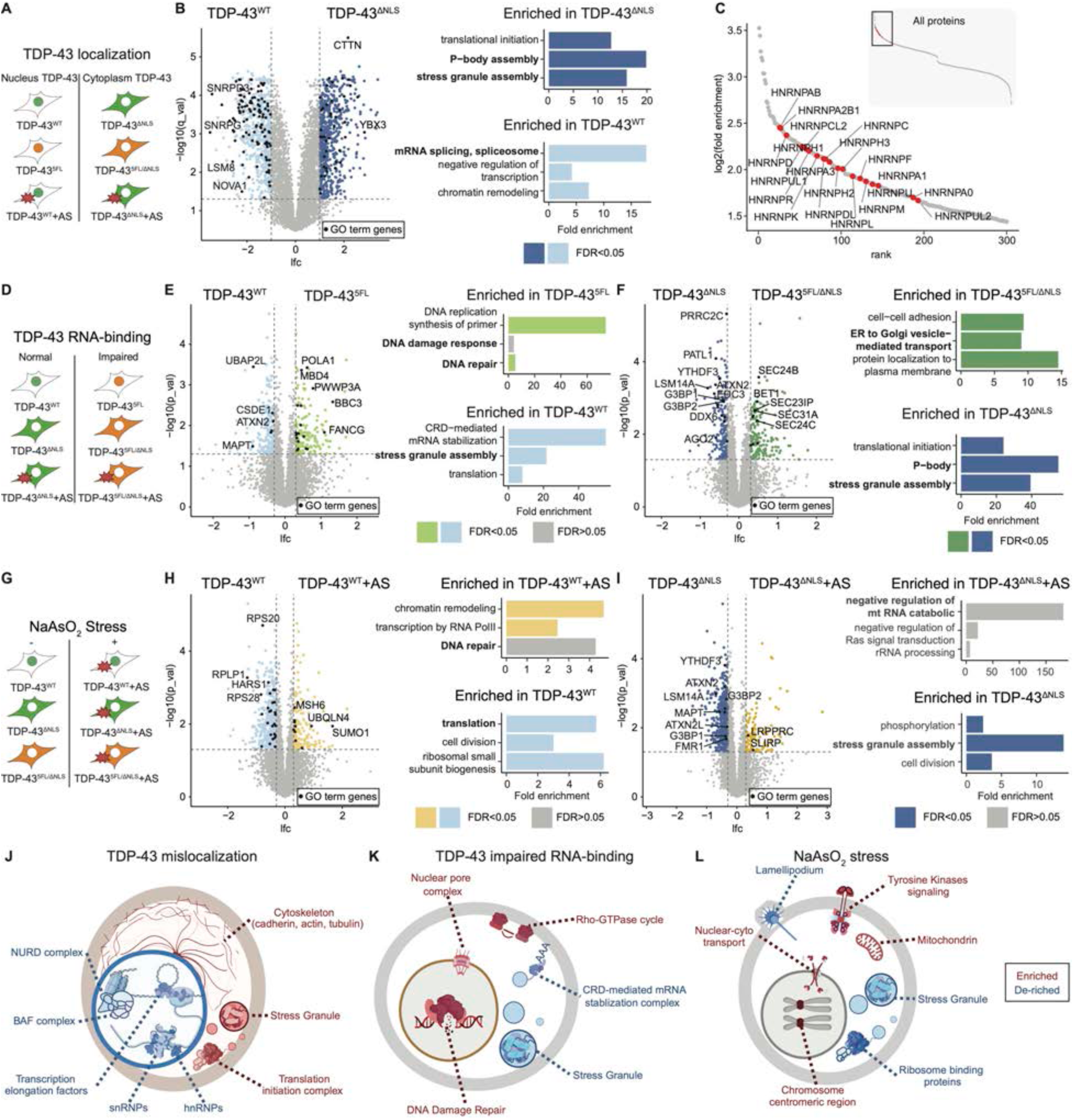
Mislocalization, impaired RNA binding, and NaAsO2 lead to distinct TDP-43 interactomes. (A) Schematic diagram of the pathological comparison design for mislocalization, with ΔNLS as the single variable. Direct comparisons include TDP-43^ΔNLS^ vs TDP-43^WT^, TDP-43^5FL/ΔNLS^ vs TDP-43^5FL^, and TDP-43^ΔNLS^+AS vs TDP-43^WT^+AS. (B) Representative volcano plot (left) of statistical significance against the log2 fold change (TDP- 43^ΔNLS^ vs TDP-43^WT^) of protein interactors labelled by APEX2-TDP-43 variants and the top-3 (based on p values) GO term analysis of the proteins enriched on each side. The q values were calculated using two-sided Student’s t-test. Proteins with q value < 0.05 and log2FC < -1 are regarded as enriched in TDP-43^WT^ (top-left quadrants); with q value < 0.05 and log2FC > 1 are regarded as enriched in TDP-43^ΔNLS^ (top-right quadrants). (C) Ranking plot of the log2 fold change of the proteins in TDP-43^ΔNLS^ vs TDP-43^WT^ with enlargement on the top-left quadrants and labeling of hnRNPs. (D) Schematic diagram of the pathological comparison design for impaired RNA-binding, with 5FL as the single variable. Direct comparisons include TDP-43^5FL^ vs TDP-43^WT^, TDP-43^5FL/ΔNLS^ vs TDP-43^ΔNLS^, and TDP-43^5FL/ΔNLS^+AS vs TDP-43^ΔNLS^+AS. (E) Representative volcano plot (left) of statistical significance against the log2 fold change (TDP- 43^5FL^ vs TDP-43^WT^) of protein interactors labelled by APEX2-TDP-43 variants and the top-3 (based on p values) GO term analysis of the proteins enriched on each side. The unadjusted p values were calculated using two-sided Student’s t-test. Proteins with p value < 0.05 and log2FC < -0.3 are regarded as enriched in TDP-43^WT^ (top-left quadrants); with p value < 0.05 and log2FC > 0.3 are regarded as enriched in TDP-43^5FL^ (top-right quadrants). (F) Representative volcano plot (left) of statistical significance against the log2 fold change (TDP- 43^5FL/ΔNLS^ vs TDP-43^ΔNLS^) of protein interactors labelled by APEX2-TDP-43 variants and the top- 3 (based on p values) GO term analysis of the proteins enriched on each side. The unadjusted p values were calculated using two-sided Student’s t-test. Proteins with p value < 0.05 and log2FC < -0.3 are regarded as enriched in TDP-43^ΔNLS^ (top-left quadrants); with p value < 0.05 and log2FC > 0.3 are regarded as enriched in TDP-43^5FL/ΔNLS^ (top-right quadrants). (G) Schematic diagram of the pathological comparison design for NaAsO2 stress, with +AS as the single variable. Direct comparisons include TDP-43^WT^+AS vs TDP-43^WT^, TDP-43^ΔNLS^+AS vs TDP-43^ΔNLS^, and TDP-43^5FL/ΔNLS^+AS vs TDP-43^5FL/ΔNLS^. (H) Representative volcano plot (left) of statistical significance against the log2 fold change (TDP- 43^WT^+AS vs TDP-43^WT^) of protein interactors labelled by APEX2-TDP-43 variants and the top-3 (based on p values) GO term analysis of the proteins enriched on each side. The unadjusted p values were calculated using two-sided Student’s t-test. Proteins with p value < 0.05 and log2FC < -0.3 are regarded as enriched in TDP-43^WT^ (top-left quadrants); with p value < 0.05 and log2FC > 0.3 are regarded as enriched in TDP-43^WT^+AS (top-right quadrants). **(I)** Representative volcano plot (left) of statistical significance against the log2 fold change (TDP- 43^ΔNLS^+AS vs TDP-43^ΔNLS^) of protein interactors labelled by APEX2-TDP-43 variants and the top-3 (based on p values) GO term analysis of the proteins enriched on each side. The unadjusted p values were calculated using two-sided Student’s t-test. Proteins with p value < 0.05 and log2FC < -0.3 are regarded as enriched in TDP-43^ΔNLS^ (top-left quadrants); with p value < 0.05 and log2FC > 0.3 are regarded as enriched in TDP-43^ΔNLS^+AS (top-right quadrants). **(J)** Schematic diagram of significantly lost (blue) and gained (red) interacting protein clusters by STRING database under mislocalization. Proteins with at least 2 times significantly enriched on each side of the direct comparisons in (A) are included. **(K)** Schematic diagram of significantly lost (blue) and gained (red) interacting protein clusters by STRING database under impaired RNA-binding. Proteins with at least 2 times significantly enriched on each side of the direct comparisons in (D) are included. **(L)** Schematic diagram of significantly lost (blue) and gained (red) interacting protein clusters by STRING database under NaAsO2 stress. Proteins with at least 2 times significantly enriched on each side of the direct comparisons in (G) are included.

To systematically profile TDP-43 interactome changes under different pathological conditions, we combined APEX2-mediated proximity labeling with streptavidin pull-down of biotinylated proteins (Extended Figure 2F–2G) and label-free data-independent acquisition mass spectrometry (DIA-MS). DIA-MS enables accurate peptide quantification and deep proteome coverage ^52,53^. We selected three key pathological factors that induce TDP-43 LOF—mislocalization (ΔNLS), impaired RNA binding (5FL), and NaAsO₂ stress (+AS)—along with their double and triple combinations (Figure 1G). To account for compartment-specific effects, we included doxycycline-inducible stable HEK293 cell lines expressing APEX2-3XNLS and APEX2-3XNES as nuclear and cytoplasmic controls, respectively (Figure 1G).

A total of 6592 proteins were quantified across four biological replicates for all conditions. Principal component analysis (PCA) revealed distinct clustering of interactomes based on TDP- 43’s subcellular localization, RNA-binding ability, and NaAsO₂ induced stress (Figure 1H), confirming a successful capture of context-dependent interactions. To assess dataset reliability, we compared our results with eight previously published TDP-43 interactome lists ^34,38,39,42,49–51^ using various methods and cell lines (Figure 1I-1K, Table 1), where five of them resembled TDP-43^WT^ interactomes, two of them resembled TDP-43^ΔNLS^ interactomes, and one with combined datasets. Among 1539 reported interactors, 1134 (73.68%) were also identified by our APEX2-TDP-43 system (Figure 1I), showing strong reproducibility.

We next identified the TDP-43-associated proteins under each condition by comparing them to their compartment controls (3xNLS for the nucleus, 3xNES for the cytoplasm) (Extended Figure 3A-3G, Table 2). TDP-43^WT^ and TDP-43^ΔNLS^ exhibited significant overlap with reported interactomes (Figure 1J-1K), given the relatively strict filter setting (Extended Figure 3A- 3B). Notably, while all conditions shared core interactions, each exhibited distinct interactors and overlapping features (Figure 1L–M), suggesting TDP-43 resides in diverse cellular environments. Gene ontology (GO) analysis using DAVID GO ^76,77^ further highlighted different functional classifications of TDP-43 interactors across various pathological conditions (Extended Figure 3H, Table 3). While all seven groups showed significant enrichment for mRNA processing and splicing interactors, TDP-43^ΔNLS^ preferentially interacted with proteins related to P-body formation, mRNA decapping, and stress granule (SG) assembly, while TDP-43^5FL/ΔNLS^ and TDP- 43^5FL/ΔNLS^+AS interacted with proteins involved in DNA damage repair and cell cycle regulation (Extended Figure 3H).

**Figure 3:**
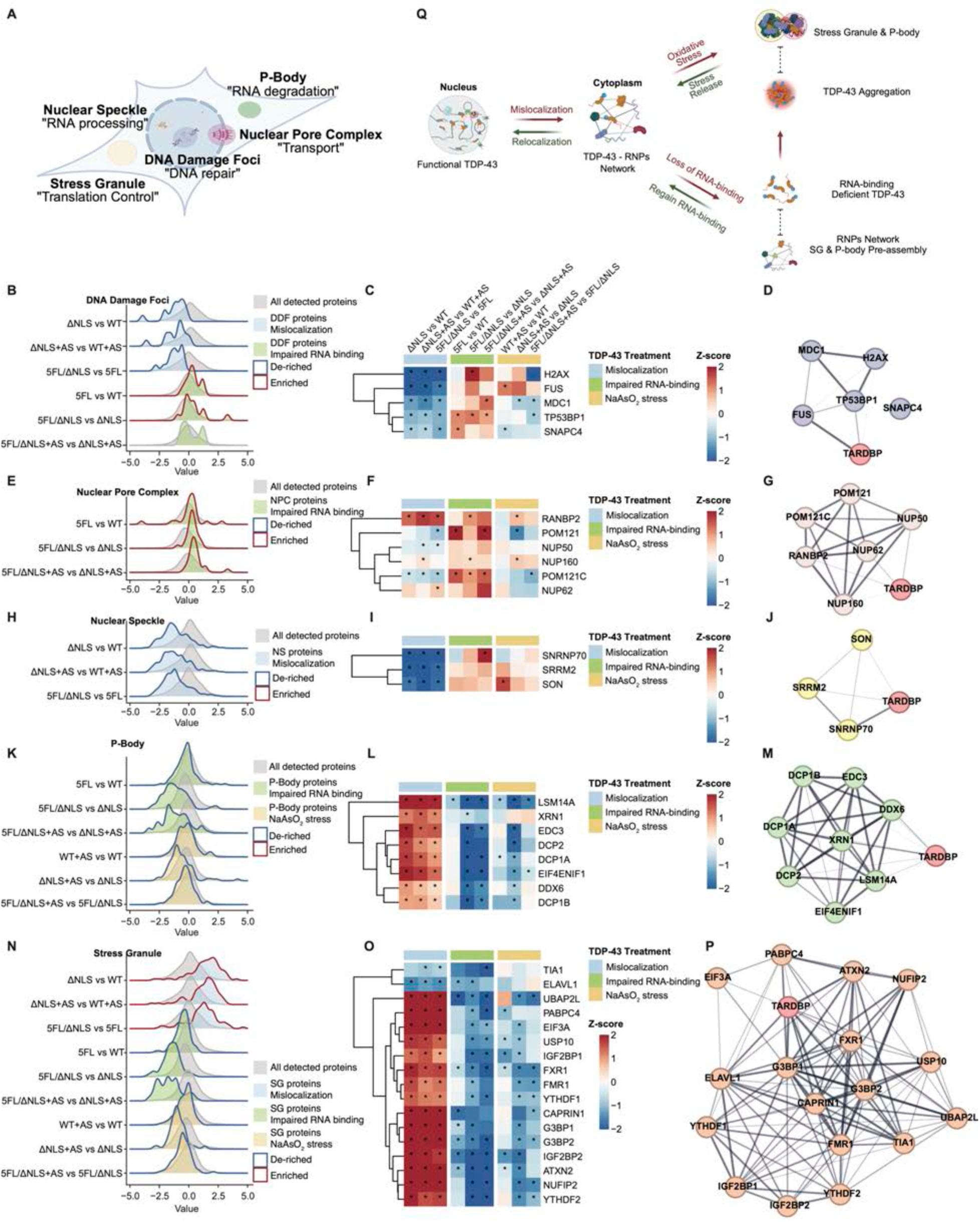
TDP-43 physiologically interacts with RNP condensates in the cytoplasm. **(A)** Schematic diagram of the subcellular locations as well as the main functions of the representative BMCs (stress granule, P-body, DNA damage foci, nuclear pore complex, and nuclear speckle). **(B)** Distribution density plots of DNA damage foci proteins and all proteins in the comparisons of TDP-43^ΔNLS^ vs TDP-43^WT^, TDP-43^5FL/ΔNLS^ vs TDP-43^5FL^, TDP-43^ΔNLS^+AS vs TDP-43^WT^+AS, TDP-43^5FL^ vs TDP-43^WT^, TDP-43^5FL/ΔNLS^ vs TDP-43^ΔNLS^, and TDP-43^5FL/ΔNLS^+AS vs TDP- 43^ΔNLS^+AS. **(C)** Heatmap of critical protein components of DNA damage foci from the nine pathological comparisons. **(D)** Protein network graph by STRING database of TDP-43 and critical protein components of DNA damage foci. **(E)** Distribution density plots of nuclear pore complex proteins and all proteins in the comparisons of TDP-43^5FL^ vs TDP-43^WT^, TDP-43^5FL/ΔNLS^ vs TDP-43ΔNLS, and TDP-43^5FL/ΔNLS^+AS vs TDP- 43^ΔNLS^+AS. **(F)** Heatmap of critical protein components of nuclear pore complex from the nine pathological comparisons. **(G)** Protein network graph by STRING database of TDP-43 and critical protein components of nuclear pore complex. **(H)** Distribution density plots of nuclear speckle proteins and all proteins in the comparison of TDP-43^ΔNLS^ vs TDP-43^WT^, TDP-43^5FL/ΔNLS^ vs TDP-43^5FL^, and TDP-43^ΔNLS^+AS vs TDP-43^WT^+AS. **(I)** Heatmap of critical protein components of nuclear speckle from the nine pathological comparisons. **(J)** Protein network graph by STRING database of TDP-43 and critical protein components of nuclear speckle. **(K)** Distribution density plots of P-body proteins and all proteins in the comparisons of TDP-43^5FL^ vs TDP-43^WT^, TDP-43^5FL/ΔNLS^ vs TDP-43^ΔNLS^, TDP-43^5FL/ΔNLS^+AS vs TDP-43^ΔNLS^+AS, TDP- 43^WT^+AS vs TDP-43^WT^, TDP-43^ΔNLS^+AS vs TDP-43^ΔNLS^, and TDP-43^5FL/ΔNLS^+AS vs TDP-435FL/ΔNLS. **(L)** Heatmap of critical protein components of P-body from the nine pathological comparisons. **(M)** Protein network graph by STRING database of TDP-43 and critical protein components of P- body. **(N)** Distribution density plots of stress granule proteins and all proteins in the comparisons of TDP- 43^ΔNLS^ vs TDP-43^WT^, TDP-43^5FL/ΔNLS^ vs TDP-43^5FL^, TDP-43^ΔNLS^+AS vs TDP-43^WT^+AS, TDP-43^5FL^ vs TDP-43^WT^, TDP-43^5FL/ΔNLS^ vs TDP-43^ΔNLS^, TDP-43^5FL/ΔNLS^+AS vs TDP-43^ΔNLS^+AS, TDP-43^WT^+AS vs TDP-43^WT^, TDP-43^ΔNLS^+AS vs TDP-43^ΔNLS^, and TDP-43^5FL/ΔNLS^+AS vs TDP- 435FL/ΔNLS. **(O)** Heatmap of critical protein components of stress granule from the nine pathological comparisons. **(P)** Protein network graph by STRING database of TDP-43 and critical protein components of stress granule. **(Q)** Schematic diagram of the proposed mechanism of TDP-43’s interaction with cytoplasm RNPs (stress granule and P-body) under different conditions. For (C), (F), (I), (L), and (O), cell color represents normalized mean z-values with “*” labels p value < 0.05. The p values were calculated using two-sided Student’s t-test.

### Differential Interactomes Due to TDP-43 Mislocalization, Impaired RNA binding, and NaAsO2-induced Stress

We performed nine comparisons to determine the impact of TDP-43 mislocalization (Figure 2A- 2C), impaired RNA binding (Figure 2D-2F), and NaAsO2-induced stress (Figure 2G-2I) on its interactome. Using APEX2-TDP-43 variants, we identified specific interactors gained or lost under each condition, highlighting key functional shifts in TDP-43 protein networks (Figure 2, Tables 4-6). For TDP-43 mislocalization with ΔNLS as the single variable (Figure 2A, Table 6), interactome analysis revealed a significant loss of interactions with mRNA splicing factors as judged by GO term enrichment (Figure 2B, Table 6). Network analysis identified gene clusters that were lost in TDP-43’s interactome under mislocalization. These included: hnRNPs and snRNPs; transcription elongation factors including POLR2A and CDK9 ^78^; NURD complex & Histone deacetylase including RBBP4, MTA1, MBD3, CHD3, and HDAC3 ^79,80^; and BAF complex (SWI/SNF proteins)^81,82^ (Extended Figure 4A, Figure 2J, Table 6). The significant reduction of associated hnRNPs was evident in the ranking plot of lost interactors (Figure 2C), supporting TDP-43’s critical function in the regulation of splicing in the nucleus together with different hnRNPs. In the cytoplasm, mislocalized TDP-43 gained interactions with translational initiation factors, SG and P-body components (Figure 2B), as well as cytoskeleton proteins (Extended Figure 4B, Figure 2J, Table 6). These results underscore the critical role of TDP-43’s localization in maintaining its association with RNA processing machinery in both the nucleus and cytoplasm, while also linking mislocalization to pathological aggregation-related proteins.

**Figure 4:**
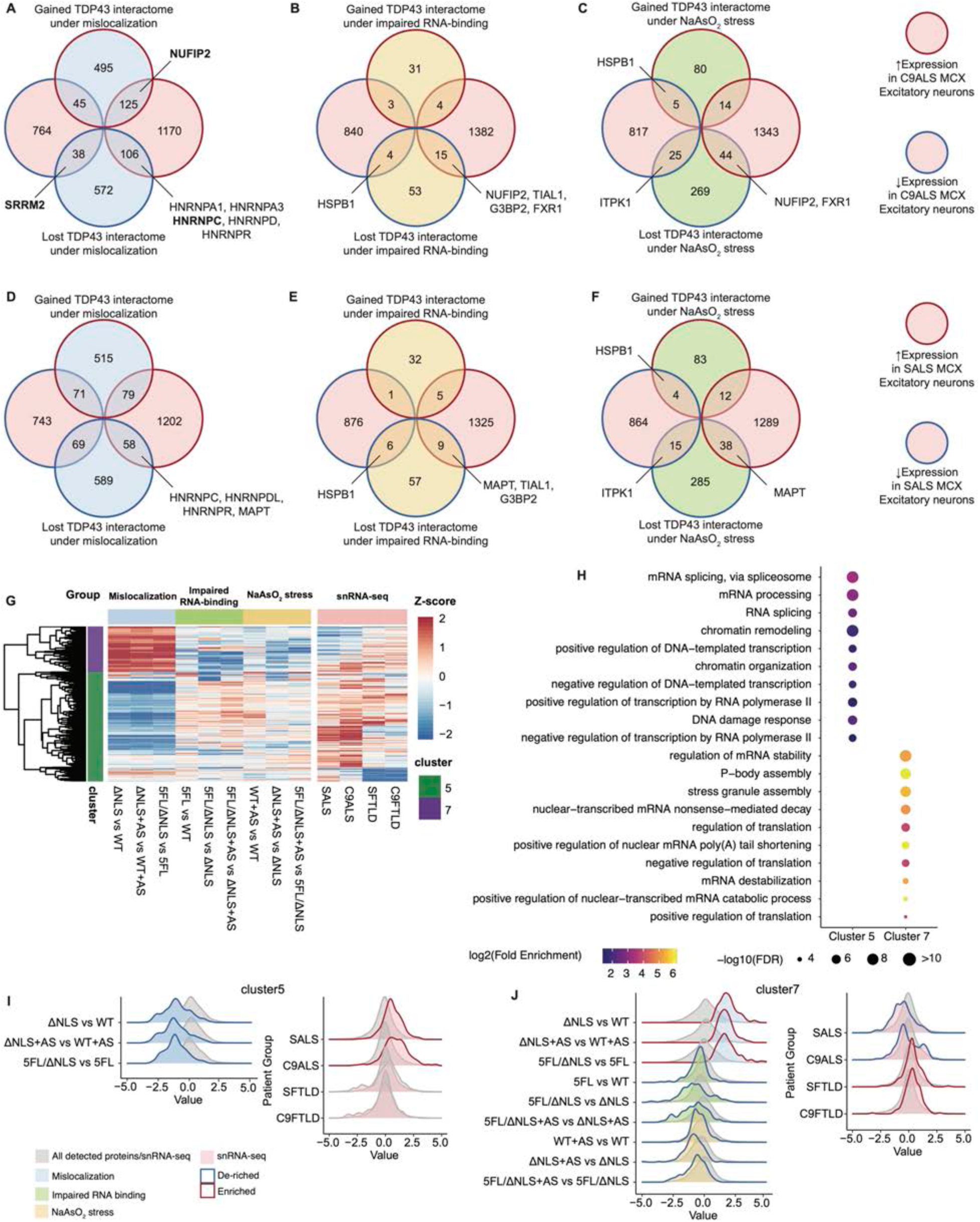
Context-dependent TDP-43 interactors are altered in ALS/FTLD patients. (A) Venn diagram between lost or gained TDP-43 interacting protein-coding genes under mislocalization by APEX2 proximity labeling (vertical axis) and the up or down-regulated expression of the genes in the excitatory neurons of the motor cortex (MCX) from C9-ALS patients by snRNA-seq (horizontal axis). (B) Venn diagram between lost or gained TDP-43 interacting protein-coding genes under impaired RNA-binding by APEX2 proximity labeling (vertical axis) and the up or down-regulated expression of the genes in the excitatory neurons of the motor cortex (MCX) from C9-ALS patients by snRNA-seq (horizontal axis). (C) Venn diagram between lost or gained TDP-43 interacting protein-coding genes under NaAsO2 stress by APEX2 proximity labeling (vertical axis) and the up or down-regulated expression of the genes in the excitatory neurons of the motor cortex (MCX) from sALS patients by snRNA-seq (horizontal axis). (D) Venn diagram between lost or gained TDP-43 interacting protein-coding genes under mislocalization by APEX2 proximity labeling (vertical axis) and the up or down-regulated expression of the genes in the excitatory neurons of the motor cortex (MCX) from sALS patients by snRNA-seq (horizontal axis). (E) Venn diagram between lost or gained TDP-43 interacting protein-coding genes under impaired RNA-binding by APEX2 proximity labeling (vertical axis) and the up or down-regulated expression of the genes in the excitatory neurons of the motor cortex (MCX) from sALS patients by snRNA-seq (horizontal axis). (F) Venn diagram between lost or gained TDP-43 interacting protein-coding genes under NaAsO2 stress by APEX2 proximity labeling (vertical axis) and the up or down-regulated expression of the genes in the excitatory neurons of the motor cortex (MCX) from C9-ALS patients by snRNA-seq (horizontal axis). (G) Representative heatmap of the gene clusters acquired from the combination of APEX2 proximity labeling with the nine pathological comparisons and snRNA-seq of the excitatory neurons in the motor cortex of ALS or FTLD patients. Clustering is performed by hierarchical clustering method. Cell color represents normalized mean z-values. Cluster 5 and 7 from a total of 10 clusters are demonstrated. (H) Gene ontology (GO) term analysis of genes enriched in Cluster 5 and Cluster 7 from (D). FDR was calculated based on adaptive linear step-up adjusted p-values. (I) Distribution density plots of the proteins from the genes in Cluster 5. Left plot shows all detected proteins from APEX2 proximity labeling in the comparisons of TDP-43^ΔNLS^ vs TDP- 43^WT^, TDP-43^5FL/ΔNLS^ vs TDP-43^5FL^, and TDP-43^ΔNLS^+AS vs TDP-43^WT^+AS. Right plot shows all detected genes from snRNA-seq in the comparisons of sALS, C9-ALS, sFTLD, and C9-FTLD patients with controls in the motor cortex excitatory neurons. (J) Distribution density plots of the proteins from the genes in Cluster 7. Left plot shows all detected proteins from APEX2 proximity labeling in the comparisons of TDP-43^ΔNLS^ vs TDP- 43^WT^, TDP-43^5FL/ΔNLS^ vs TDP-43^5FL^, TDP-43^ΔNLS^+AS vs TDP-43^WT^+AS, TDP-43^5FL^ vs TDP-43^WT^, TDP-43^5FL/ΔNLS^ vs TDP-43^ΔNLS^, TDP-43^5FL/ΔNLS^+AS vs TDP-43^ΔNLS^+AS, TDP-43^WT^+AS vs TDP-43^WT^, TDP-43^ΔNLS^+AS vs TDP-43^ΔNLS^, and TDP-43^5FL/ΔNLS^+AS vs TDP-43^5FL/ΔNLS^. Right plot shows all detected genes from snRNA-seq in the comparisons of sALS, C9-ALS, sFTLD, and C9-FTLD patients with controls in the motor cortex excitatory neurons.

Impaired RNA binding, modeled by expressing the TDP-43^5FL^ and TDP-43^5FL/ΔNLS^ variants (Figure 2D, Table 6), led to loss of interactions with the cytoplasmic RNA processing machinery, particularly with stress granules (SGs) and P-bodies – interactions that are enriched under mislocalization (Figure 2E-2F, Extended Figure 4C, Table 6). This loss was accompanied by a gain in interactions with proteins associated with DNA damage repair pathways, Rho-GTPase signaling (including DSG2 and PKP4) ^83^, and the nuclear pore complex (Figure 2E, Extended Figure 4D, Table 6). These findings suggest that the RNA-binding ability of TDP-43 is critical for its engagement with SG and P-body proteins, and its disruption may redirect TDP-43 toward DNA repair and transport processes (Figure 2K, Table 6).

NaAsO2-induced oxidative stress introduced a unique shift in the TDP-43 interactome (Figure 2G, Table 6). Although TDP-43 is a well-established component of SGs ^32^ and P-bodies^33^, its interactions with these structures were unexpectedly lost under NaAsO2 stress, which induces the formation of SGs and P-bodies ^84^. Additionally, associations of TDP-43 with ribosomal proteins (especially from the 43S complex), chromosome centromeric region (including CCNB1, HJURP, and NUF2) ^85^, and lamellipodium-associated proteins (including ABI1, BAIAP2, and CTTN) were also lost (Figure 2H-I, Extended Figure 4E, Table 6). Conversely, TDP-43 gained associations with tyrosine kinase signaling proteins (including Ras signaling), mitochondrial proteins (including TOMM34, PC, PCCB, and ACACA)^86–91^, and nucleocytoplasmic transport factors (IPO4, IPO5, and XPO5)^92^ ^34,93–95^ (Figure 2I, Extended Figure 4F, Table 6). Interestingly, under NaAsO2 stress, TDP-43^WT^ exhibited bidirectional alterations in its interaction with mRNA splicing factors (Figure 2H, Table 6). Specifically, interactions with EFTUD2, RBM25, SON, FUS, DHX15, CRIPT, PPIG, HNRNPUL1, and endogenous TDP-43 (suggesting self-aggregation or oligomerization) were gained, while interactions with SRSF3, SRSF10, and SRRM1 were lost. These observations highlight how environmental and extracellular stressors (e.g. NaAsO2) can modulate TDP-43 interactomes and functions, implicating additional pathways in stress-induced pathobiology.

In addition to identifying the interactome, we aimed to determine whether our approach could capture known interactors and reveal their pathological relationship with TDP-43. Of the 1134 reported proteins (Extended Figure 5A), we modeled each pathological factor independently, adjusting using the other two factors as confounders (see Methods), and examined the changes in TDP-43 interactions under mislocalization (Extended Figure 5B), impaired RNA binding (Extended Figure 5C), and NaAsO2 stress (Extended Figure 5D). We further investigated nine individual proteins previously reported as modifiers of TDP-43 pathobiology, including HSPB1^39,96^, ATXN2^97–102^, DBR1^103^, POM121^34,104–106^, HSPA5^107^, UPF1^108–110^, NUP214^34^, XPO5^34^, and TP53BP1^111^, and mapped their interactions with TDP-43 under different conditions (Extended Figure 5E-5F). Notably, HSPB1, a stress response protein, showed increased interaction with TDP-43 under NaAsO2 stress in all three comparisons (Extended Figure 5E-F), consistent with recent findings ^39^. We show significantly decreased HSPB1 in 5FL variants, supporting that TDP-43 RNA recognition motifs (RRMs) are necessary for HSPB1-TDP-43’s interaction, suggesting the potential involvement of RNA (Extended Figure 5E-5F). Furthermore, we performed correlation analysis by calculating the Pearson correlation coefficient (PCC) matrix for the known modifiers and TDP-43 interactors across all nine pathological comparisons (Table 7), which may identify TDP-43 regulators with similar effects. Together, these results support the robustness of our system in identifying conserved and novel mechanisms in TDP-43 dysfunction.

**Figure 5:**
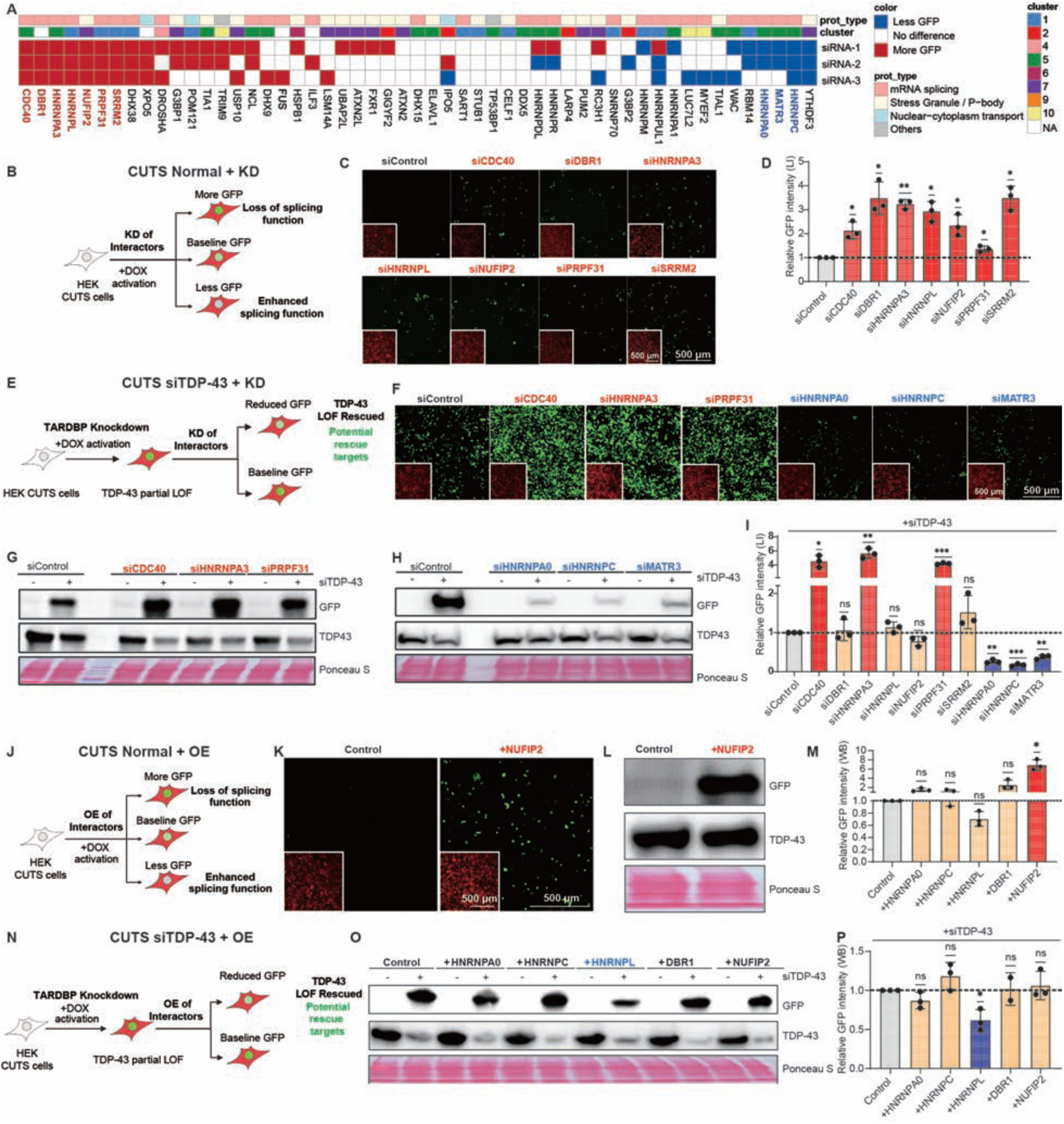
TDP-43 interactors regulate its splicing function bidirectionally. **(A)** Heatmap of the ranked preliminary CUTS siRNA screening result (N = 3 biological replicates) of 53 selected proteins mostly from Cluster 1, 5, and 7 of three categories (mRNA splicing, stress granule or P-body, and nuclear-cytoplasm transport). Cell color shows the summary of the results from each replicate with whether increasing GFP signal (causing LOF, +1) or lowering GFP signal (rescuing LOF, -1) comparing to control siRNA (siControl). **(B)** Schematic diagram of the experimental design of normal CUTS with siRNA knockdown of TDP-43’s interactors. **(C)** Representative live confocal images of the stable HEK293 cells expressing doxycycline-inducible (72 h of 1 mg/mL doxycycline) CUTS with reverse transfected siRNA control (siControl) or the top hits that cause LOF in the preliminary screening in (A) (20 nM, 96 h). **(D)** Relative GFP fluorescence intensity quantification normalized to siControl from live confocal imaging in (C) (N=3 biological replicates). **(E)** Schematic diagram of the experimental design of CUTS under partial TDP-43 knockdown (siTDP-43) with siRNA knockdown of TDP-43’s interactors. **(F)** Representative live confocal images of the stable HEK293 cells expressing doxycycline-inducible (72 h of 1 mg/mL doxycycline) CUTS with reverse co-transfected siTDP-43 (5 nM, 96 h) with siControl or the top hits that cause or rescue LOF in the preliminary screening in (A) (20 nM, 96 h). **(G)** Western blot analysis of the stable HEK293 cells expressing doxycycline-inducible (72 h of 1 mg/mL doxycycline) CUTS with reverse co-transfected siTDP-43 (5 nM, 96 h) with siControl or siCDC40, siHNRNPA3, and siPRPF31 (20 nM, 96 h) that exacerbate the LOF in (F). **(H)** Western blot analysis of the stable HEK293 cells expressing doxycycline-inducible (72 h of 1 mg/mL doxycycline) CUTS with reverse co-transfected siTDP-43 (5 nM, 96 h) with siControl or siHNRNPA0, siHNRNPC, siMATR3 (20 nM, 96 h) that rescue the LOF in (F). **(I)** Relative GFP fluorescence intensity quantification normalized to siControl (+siTDP-43) from live confocal imaging in (F) (N=3 biological replicates). **(J)** Schematic diagram of the experimental design of normal CUTS with cDNA overexpression of TDP-43’s interactors. **(K)** Representative live confocal images of the stable HEK293 cells expressing doxycycline-inducible (72 h of 1 mg/mL doxycycline) CUTS with control (mock) or transfection of CMV-driven NUFIP2 cDNA in a plasmid (500 ng, 72 h). For (C), (F), and (K), Green = GFP signal; Red = mCherry signal. Scale bar = 100 µm. **(L)** Western blot analysis of the cells in (K). **(M)** Relative GFP protein intensity quantification normalized to control from western blot analysis of the stable HEK293 cells expressing doxycycline-inducible (72 h of 1 mg/mL doxycycline) CUTS with control (mock) or transfection of five available cDNA plasmids of the hits driven by CMV promoter (500 ng, 72 h). (N=3 biological replicates). **(N)** Schematic diagram of the experimental design of CUTS under partial siTDP-43 with cDNA overexpression of TDP-43’s interactors. **(O)** Western blot analysis of the stable HEK293 cells expressing doxycycline-inducible (72 h of 1 mg/mL doxycycline) CUTS with reverse transfected siTDP-43 (5 nM, 96 h) and control (mock) or transfection of five available cDNA plasmids of the hits driven by CMV promoter (500 ng, 72 h). **(P)** Relative GFP protein intensity quantification normalized to control from western blot analysis in (O) (N=3 biological replicates). For (D), (I), (M), and (P), statistical significance was determined by one-sample two-sided t-test (* = P < 0.05; ** = P < 0.01; *** = P < 0.001; **** = P < 0.0001). Data are the mean ± sd.

### Systematic Mapping of TDP-43 Interactions with Biomolecular Condensate

Emerging evidence underscores the pathological significance of BMCs, membraneless subcellular compartments created through phase separation, in neurodegenerative diseases^112–114^. As a hallmark in ALS/FTLD and a frequent co-pathology in AD, TDP-43 is a component of various BMCs under physiological and pathological conditions^36,37,115^. To systematically characterize these interactions, we performed the global mapping of TDP-43 interactions with BMCs under mislocalization, impaired RNA binding, and NaAsO₂-induced stress conditions. Unlike functional enrichment, where only significantly altered proteins are selected, this analysis considers all known components of specific BMCs to acquire an overall landscape, owing to our high-throughput quantitative coverage. These results reveal insights into the context-dependent altered interactions between TDP-43 and 19 previously annotated BMCs, including DNA damage foci, nuclear pore complex (NPC), nuclear speckle, SGs, and P-body (Figure 3A, Extended Figure 6A, Table 8).

**Figure 6:**
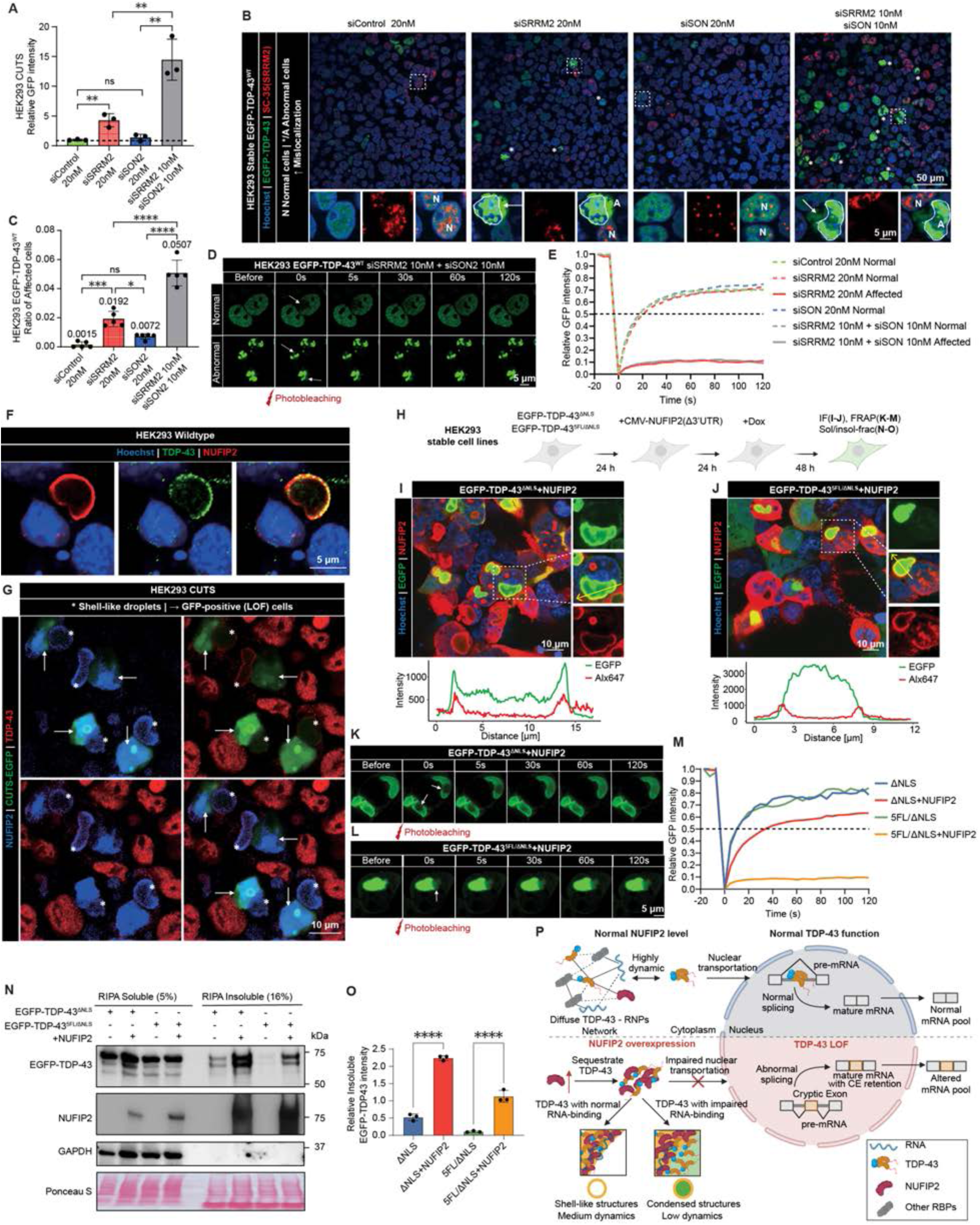
SRRM2 loss or NUFIP2 overexpression induces TDP-43 mislocalization, aggregation, and loss of function. **(A)** Relative GFP fluorescence intensity quantification from live confocal imaging in stable HEK293 cells expressing doxycycline-inducible (72 h of 1 mg/mL doxycycline) CUTS with reverse transfected siRNA control (siControl) (20 nM, 96 h), SRRM2 (siSRRM2) (20 nM, 96 h), SON (siSON) (20 nM, 96 h), or siSRRM2 + siSON (10 nM each, 96 h). Statistical significance was determined by a two-sided Student’s t-test (* = P < 0.05; ** = P < 0.01; *** = P < 0.001; **** = P < 0.0001). Data are the mean ± s.d. **(B)** Representative immunofluorescence staining images of stable HEK293 cells expressing EGFP- TDP-43^WT^ with reverse transfected siControl, siSRRM2, siSON (20 nM, 96 h), or siSRRM2 + siSON (10 nM each, 96 h). “Normal” cells are labeled as “N”, “abnormal” cells are labeled as “A” or “*”. The nuclei of “abnormal” cells are outlined. T Blue = Hoechst; Green = EGFP-TDP-43; Red = SC-35 (SRRM2). Scale bar = 5 µm. **(C)** Ratio of the abnormal cells (with nuclear inclusions and cytoplasmic mislocalization of EGFP- TDP-43^WT^) from all the cells (not including dead cells with disrupted nucleus) in (B) (N=5 replicates with each >230 cells). **(D)** Representative images of FRAP analysis of “normal” or “abnormal” (nuclear inclusions and cytoplasmic mislocalization) EGFP- TDP-43^WT^ under siSRRM2 + siSON (10 nM each, 96 h). **(E)** FRAP graph of the “normal” or “abnormal” EGFP- TDP-43^WT^ from siControl, siSRRM2, siSON (20 nM, 96 h), or siSRRM2 + siSON (10 nM each, 96 h) (N>=8 for each condition). **(F)** Representative immunofluorescence staining images of wildtype HEK293 cells with NUFIP2 overexpression. Blue = Hoechst; Green = TDP-43; Red = NUFIP2. Scale bar = 2 µm. **(G)** Representative immunofluorescence staining images of the stable HEK293 cells expressing doxycycline-inducible (48 h of 1 mg/mL doxycycline) CUTS with or without NUFIP2 overexpression via the transfection of CMV-NUFIP2(Δ3’UTR). Blue = NUFIP2; Green = EGFP from CUTS; Red = TDP-43. Scale bar = 5 µm. **(H)** Schematic diagram of the experiment design to test the impact of NUFIP2 overexpression on cytoplasmic TDP-43 with or without impaired RNA-binding. IF, immunofluorescence staining; FRAP, fluorescence recovery after photobleaching; Sol/insol-frac, RIPA soluble/insoluble fractionation. **(I)** Representative immunofluorescence staining images of the “shell-like” structures in stable HEK293 cells expressing EGFP- TDP-43^ΔNLS^ under NUFIP2 overexpression. Magnified views (right) of the region with a white square in the main image (left) represent the example of a “shell- like” structure. An intensity plot is shown for EGFP- TDP-43^ΔNLS^ (green) and NUFIP2 (red) following the direction of the arrow. **(J)** Representative immunofluorescence staining images of the “condensed” structures in stable HEK293 cells expressing EGFP- TDP-435FL/ΔNLS under NUFIP2 overexpression. Magnified views (right) of the region with a white square in the main image (left) represent the example of a “condensed” structure. An intensity plot is shown for EGFP- TDP-435FL/ΔNLS (green) and NUFIP2 (red) following the arrow. **(K)** Representative images of FRAP analysis of EGFP- TDP-43^ΔNLS^ “shell-like” structures by NUFIP2 overexpression. **(L)** Representative images of FRAP analysis of EGFP- TDP-43^5FL/ΔNLS^ “condensed” structures by NUFIP2 overexpression. For (K) and (L), different time points before or after photobleaching are shown with arrows indicating the bleached regions. For (I) to (L), Blue = Hoechst; Green = EGFP-TDP-43; Red = NUFIP2. **(M)** FRAP graph of the diffused EGFP- TDP-43^ΔNLS^ (green, N=6) and EGFP- TDP-43^5FL/ΔNLS^ (blue, N=5) signals together with EGFP- TDP-43^ΔNLS^ structures (red, N=6) and EGFP- TDP- 43^5FL/ΔNLS^ aggregates (purple, N=6) by NUFIP2 overexpression. **(N)** Representative western blot analysis of EGFP- TDP-43^ΔNLS^ and EGFP- TDP-43^5FL/ΔNLS^ with or without NUFIP2 overexpression in RIPA-solution and insoluble fractions from HEK293 cells. **(O)** Quantification of the relative insoluble EGFP-TDP-43 intensities to ponceau from the western blots in (R) (N=3 biological replicates). Statistical significance was determined by one-way ANOVA and Tukey’s multiple comparison test (* = P < 0.05; ** = P < 0.01; *** = P < 0.001; **** = P < 0.0001). Data are the mean ± s.d. **(P)** Working model of the impacts on TDP-43 by NUFIP2 overexpression. When overexpressed, NUFIP2 sequestrates TDP-43 in the cytoplasm to form “shell-like” structures. This phase transition impairs TDP-43’s nuclear localization thus leads to its LOF in splicing, creating CE retention. TDP-43 could be further transformed into “condensed” aggregates with significantly decreased dynamics when it loses RNA-binding.

TDP-43 displayed opposing interaction patterns with DNA damage foci under different pathological conditions (Figure 3B)^51^. When mislocalized, TDP-43 greatly lost its interaction with most DNA damage foci proteins, indicating proper nuclear localization is essential for its involvement in DNA damage repair (Figure 3B-C)^116^. However, with impaired RNA binding, TDP-43 gained interactions with DNA damage-associated proteins (Figure 3B-3C), especially TP53BP1, consistent with its previously reported RNA-independent interaction with TDP-43^111^. This suggests that TDP-43 is recruited to genomic repair components when RNA binding is disrupted (Figure 3D), though the impact on repair pathways is unknown.

Similarly, TDP-43 significantly gained interactions with NPC components under impaired RNA binding conditions (Figure 3E-3F). This supports several lines of emerging evidence of NPC’s involvement with TDP-43 pathobiology (Figure 3G)^34,35^. Among them, interestingly, we detected a strikingly elevated interaction between POM121 (and its homolog, POM121C) and TDP-43, which is impaired in ALS/FTLD^104,105^. This adds to previous findings that RNA-deficient TDP-43 mislocalizes to the cytoplasm via passive diffuse through the NPC^117^, suggesting a possible crosstalk between POM121 and RNA-deficient TDP-43.

Nuclear speckles^118^ also showed altered context-dependent interactions with TDP-43. Mislocalized TDP-43 significantly lost interactions with nuclear speckle proteins (Figure 3H-3I), including both SRRM2 and SON, the scaffold components of nuclear speckles^119^. Given the recent discovery of nuclear speckle’s participation in regulating RNA splicing and its disruption evidenced in ALS/FTLD ^120–122^, this finding suggests that proper nuclear localization is critical for TDP-43’s association with nuclear speckle and its splicing-related functions (Figure 3J).

In addition to nuclear BMCs, TDP-43 exhibited altered dynamic interactions with cytoplasmic SGs and P-bodies across the pathological conditions tested. This is consistent with our finding in functional enrichment (Figure 2J-2L). Under mislocalization conditions, TDP-43 gained robust interactions with SG and P-body proteins, consistent with their roles in cytoplasmic stress responses (Figure 3K-3P). However, these interactions were markedly diminished under impaired RNA binding, suggesting that TDP-43 RNA-binding capacity is critical for maintaining these associations (Figure 3K-3P). Notably, under NaAsO2 stress, TDP-43 lost interactions with SG and P-body proteins, contradicting prior expectations that stress universally promotes these associations (Figure 3K-3P). This pattern is consistent with most SG/P-body proteins, including several of their core components: DCP1A, DDX6, EDC3^123^, G3BP1/2^124^, UBAP2L^125^ and USP10^126^ (Figure 3L, 3O).

Cytoplasmic TDP-43 is known to aggregate under stress but is not necessarily driven by SG condensation^127^, which was also revealed by the increased labeling of endogenous TDP-43 under NaAsO2 stress (Extended Figure 2F-2G). Our findings strongly suggest that under oxidative stress, cytoplasmic TDP-43 is not localized to SGs but self-aggregates, which is consistent with several recent studies ^31,61,128^. This differs from previously established paradigms^84,129,130^ and suggests a working model in which TDP-43 interactions with SGs and P-bodies are facultative and tightly regulated. Excessive or deficient interactions may disrupt cellular homeostasis, underscoring the fine-tuned balance required for TDP-43 functionality (Figure 3Q).

TDP-43 exhibited further context-dependent shifts among other important BMCs (Extended Figure 6A). To further evaluate the impact of the pathological factors on different BMCs, we ranked their associations with BMCs in all nine pathological comparisons (Extended Figure 6B- 6J). This provided detailed insights into TDP-43 dynamic interacting patterns among BMCs.

### TDP-43 Interactors Converge with ALS/FTLD Patient-derived snRNA-seq Signatures

To determine whether the context-dependent TDP-43 interactors identified in our HEK293 cell models might uncover disease-relevant interactors, we performed an integrated analysis with a recently published snRNA-seq dataset^55^ of ALS/FTLD patients and extracted the data from excitatory neurons of the motor cortex (MCX). In support of our results, among all 6592 proteins detected in our mass spectrum dataset from HEK293 cells, 5443 (82.57%) of these also have their coding genes quantified in the snRNA-seq data from patient-derived neurons (Extended Figure 7A, Table 9). Correspondingly, differentially expressed (DE) genes from either C9-ALS (2248 genes) or sALS (2222 genes) patients are found in our proteomic data (56.27% and 52.34%) (Extended Figure 7B). This high-overlapping ratio highlights the potential of HEK293 cells for high-throughput preliminary screenings before neuronal proteomics analysis, likely due to their unique background, which shares certain molecular characteristics with neurons^131,132^. We also found 18 genome-wide association study (GWAS) annotated genes^133–135^ present in both snRNA-seq and our mass spectrum datasets (Extended Figure 7C).

**Figure 7:**
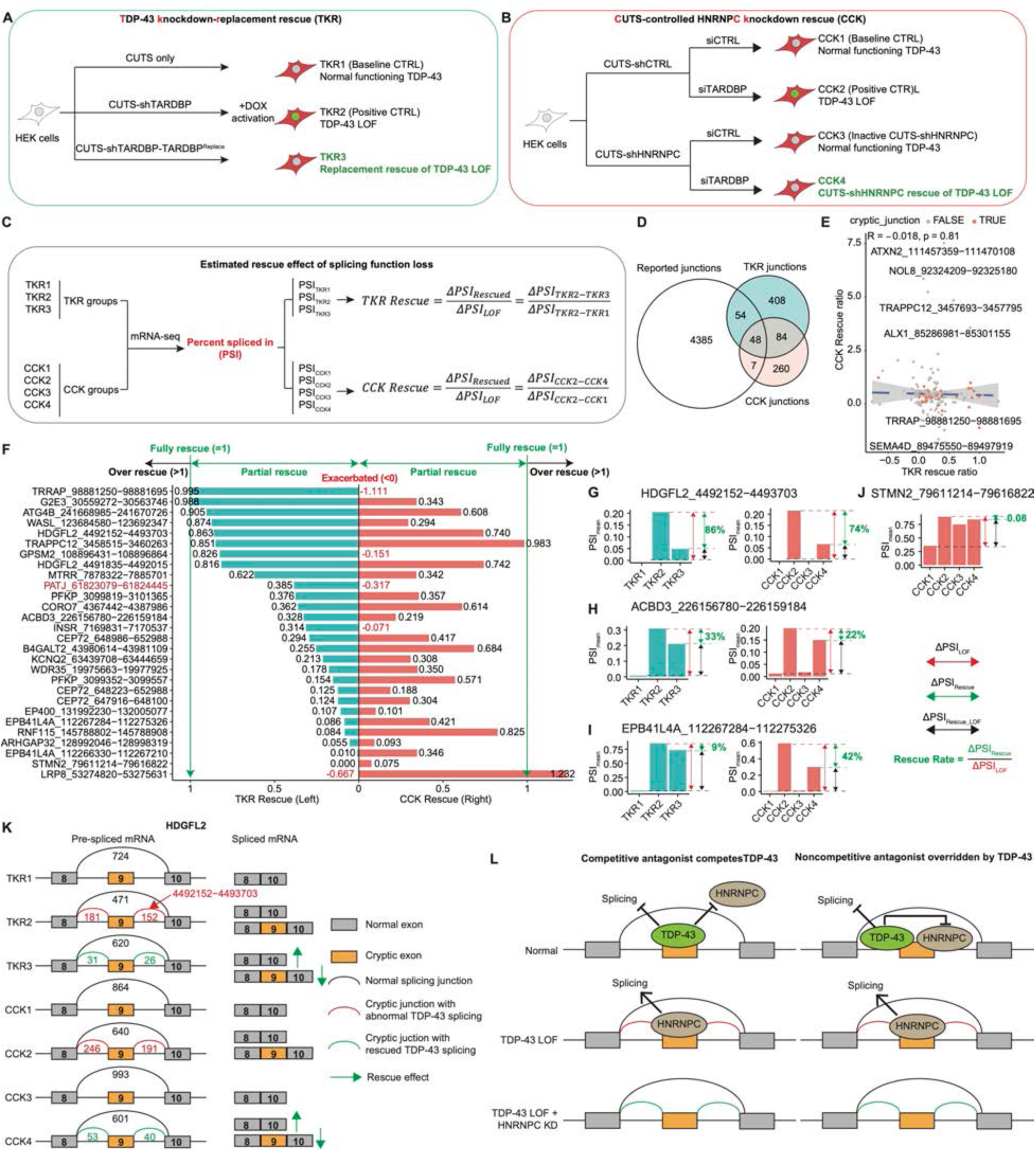
TDP-43 knockdown-replacement (TKR) and CUTS-controlled HNRNPC knockdown (CCK) partially rescue cryptic exon splicing. **(A)** Schematic diagram of TDP-43 knockdown-replacement (TKR) experimental design. Three HEK293 stable cell lines built by *Piggybac* were leveraged. **TKR1**: expressing only CUTS, as the baseline control with normal TDP-43 function; **TKR2**: expressing CUTS together with shRNA targeting endogenous TDP-43, as the positive control with TDP-43 loss of function (LOF); **TKR3**: expressing shRNA targeting endogenous TDP-43 and CUTS-controlled expressing of codon-optimized EGFP-TDP-43, as the TKR therapy group. All the three constructs were activated by doxycycline-induced expression for 4 days with N=3 biological replicates. **(B)** Schematic diagram of CUTS-controlled HNRNPC knockdown (CCK) therapy’s experiment design. Two HEK293 stable cell lines with four different siRNA treatment groups were used. **CCK1**: CUTS-shControl cell line with siControl, as the baseline control with normal TDP-43 function; **CCK2**: CUTS-shControl cell line with siTDP-43, as the positive control with TDP-43 LOF; **CCK3**: CUTS-shHNRNPC cell line with siControl, as the inactive control for CUTS- shHNRNPC with normal TDP-43 function; **CCK4**: CUTS-shHNRNPC cell line with siTDP-43, as the CCK therapy group. All the groups were activated by siRNA treatment for 7 days with N=3 biological replicates. **(C)** Schematic diagram of the mRNA sequencing samples design and the rescue ratio for both TKR and CCK groups. The rescue ratio for each group was calculated from the changing in the percent spliced-in (ΔPSI, calculated by MAJIQ) of specific splicing junctions, showing as the ratio between rescued ΔPSI (positive control vs therapy group) and LOF ΔPSI (positive control vs baseline control). **(D)** Venn diagram of the significantly changed (|ΔPSI|>0.1, probability_of_changing>0.95, calculated by MAJIQ) splicing junctions among the positive controls from TKR, CCK experiments and a previous study in iPSC-derived model. Splicing junctions with overlapping of at least 2 groups with the same direction of ΔPSIs were selected as the evaluation indexes for measuring TDP-43 LOF. **(E)** Co-relationship analysis of the rescue ratio between TKR and CCK therapies over the selected junctions from F. Cryptic junctions were labeled as they indicated the cryptic exon (CE) retention caused by TDP-43 LOF. **(F)** Bar plot of the rescue ratio from TKR (left) and CCK (right) of 28 cryptic junctions in (G). The junctions were ranked by TKR rescue ratio, from high to low. **(G)** Bar plot analysis of the PSI values of the cryptic junction in HDGFL2 (492152−4493703) in TKR groups (left) and CCK groups (right). **(H)** Bar plot analysis of the PSI values of the cryptic junction in ACBD3 (226156780−226159184) in TKR groups (left) and CCK groups (right). **(I)** Bar plot analysis of the PSI values of the cryptic junction in EPB41L4A (112267284−112275326) in TKR groups (left) and CCK groups (right). **(J)** Bar plot analysis of the PSI values of the cryptic junction in STMN2 (79611214−79616822) in CCK groups. For G-J, the grouped (bars) and individual (points) PSI values samples were calculated separately by MAJIQ. **(K)** Representative splicing plots of the cryptic exon (Exon 9) in HDGFL2 in TKR and CCK groups by Voila. **(L)** Two possible working models of HNRNPC regulating TDP-43’s splicing CE targets. HNRNPC can be a competitive antagonist of TDP-43 to its binding site on pre-mRNA (left): Under normal TDP-43 function, HNRNPC is blocked from TDP-43’s binding sites the splicing of CE is repressed; Without TDP-43’s binding (LOF), HNRNPC binds to these sites and promotes splicing, resulting in CE retention; If HNRNPC is further knocked-down, the CE retention is partially rescued. HNRNPC can also be a noncompetitive antagonist (right) with binding to the CE sites next to TDP-43 and promotes splicing. However, under normal condition HNRNPC is repressed by TDP-43; Without TDP-43, this inhibition is released and HNRNPC promotes CE retention.

We next cross-referenced lost or gained TDP-43 interactors from our APEX2 proximity labeling experiments under TDP-43 mislocalization, impaired RNA binding, or NaAsO₂-induced stress conditions with genes that were significantly up- or down-regulated in C9-ALS and sALS patient neurons (Figure 4A-4F). Notably, we found HSPB1 to be one of the few proteins having gained interaction with TDP-43 under stress while it was significantly down-regulated in both C9-ALS and sALS patient neurons (Figure 4C, 4F), which is supported by a recent study ^39^. We also identified some hnRNPs (especially HNRNPC and HNRNPR), SRRM2 (a scaffold protein of nuclear speckle), and NUFIP2 (a core protein of SG) that showed lost or gained interaction with TDP-43 under mislocalization while significantly altered in patients (Figure 4A, 4D).

To further dissect these relationships, we performed hierarchical clustering on the combined dataset of TDP-43 proximity-labeled proteins (from nine distinct pathological comparisons) and the differentially expressed genes in ALS/FTLD patient neurons (Extended Figure 7D, Table 9- 10). This analysis yielded 10 primary gene clusters, of which two showed particularly intriguing features: Cluster 5 and Cluster 7 (Figure 4G, Table 9-10). Gene Ontology (GO) enrichment of each cluster corroborated their distinct functional profiles (Figure 4H, Table 11).

Cluster 5 was enriched with mRNA splicing factors (Figure 4H), which exhibited globally lost interaction with TDP-43 under mislocalization yet were up-regulated in C9-ALS and sALS patient neurons (but not in FTLD) (Figure 4I). This pattern may reflect feedback or compensatory up- regulation of splicing machinery in response to diminished TDP-43 functional engagement. In contrast, Cluster 7 encompassed many proteins implicated in stress granules (SGs), P-bodies, and nonsense-mediated decay (NMD) (Figure 4H). These factors displayed a globally gained interaction with mislocalized TDP-43 but lost interactions under impaired RNA binding and NaAsO₂ stress, consistent with our previous analysis (Figure 4J). Notably, Cluster 7 genes showed a distinctive transcriptional response in patients—being generally down-regulated in C9-ALS and sALS, yet paradoxically up-regulated in C9-sFTLD (Figure 4J). Such divergent expression patterns might reflect differential protective or pathogenic roles of RNA granule dynamics in TDP- 43 pathology across ALS versus FTLD-TDP subtypes.

To expand our observations, we examined the broader concordance between all detected proteins in this study and differentially expressed genes in ALS or FTLD patient neurons (Extended Figure 7D, Table 11), revealing that many of these context-dependent TDP-43 interactors exhibit robust, shared signatures of dysregulation in ALS/FTLD, underscoring their potential roles in disease-related mechanisms (Extended Figure 7E-7G). Furthermore, we examined the behavior of known ALS/FTLD GWAS-associated genes^133–135^ across our TDP-43 proximity-labeled and snRNA-seq datasets for a “GTP” (Genomics-Transcriptomics-Proteomics) triple-omics analysis. Six out of 18 GWAS-reported ALS/FTLD genes (RPSA, KIF5A, TNIP1, TBK1, SPATA2, and ATRN) exhibited significant alterations in TDP-43 interactions across all three pathological factors and showed significant changes in at least one patient-derived snRNA-seq dataset (C9 or sporadic ALS/FTLD) (Extended Figure 7C, Table 12).

Together, these multiomics analyses demonstrate that TDP-43 pathological states in our APEX2 proximity labeling approach capture key molecular alterations also present in ALS/FTLD patient neurons. By pinpointing gene clusters with distinctive TDP-43 binding and transcriptomic patterns—such as the splicing-factor–enriched Cluster 5 and the cyto-RNA-granule–associated Cluster 7—our results suggest potential novel mechanistic avenues involving both upstream triggers (TDP-43 mislocalization or RNA-binding defects) and downstream compensatory or pathogenic responses (aberrant RNA splicing, granule assembly, or decay processes). This integrated view highlights how context-dependent TDP-43–protein interactions and disease-specific gene expression changes may converge to drive the pathogenesis of ALS/FTLD and related TDP-43-associated disorders.

### A Functional Splicing Screen Reveals Bidirectional Modulation of TDP-43 Function by the Disease-Relevant Interactome

Building on the multiomics integration previously described, we next sought to determine whether specific context-dependent TDP-43-interacting proteins could influence the TDP-43 splicing function. From the 10 identified clusters (Extended Figure 7D), we selected 53 proteins enriched in mRNA splicing factors (primarily from Clusters 1 and 5), SG/P-body-related components (primarily from Cluster 7), additional key nuclear-cytoplasmic transport factors, and reported TDP-43 modifiers. Their cluster assignments, broad functional classes, and dysregulation patterns in APEX2 proximity labeling and patient snRNA-seq data are shown in the heatmap in Extended Figure 8A and Table 13.

**Figure 8:**
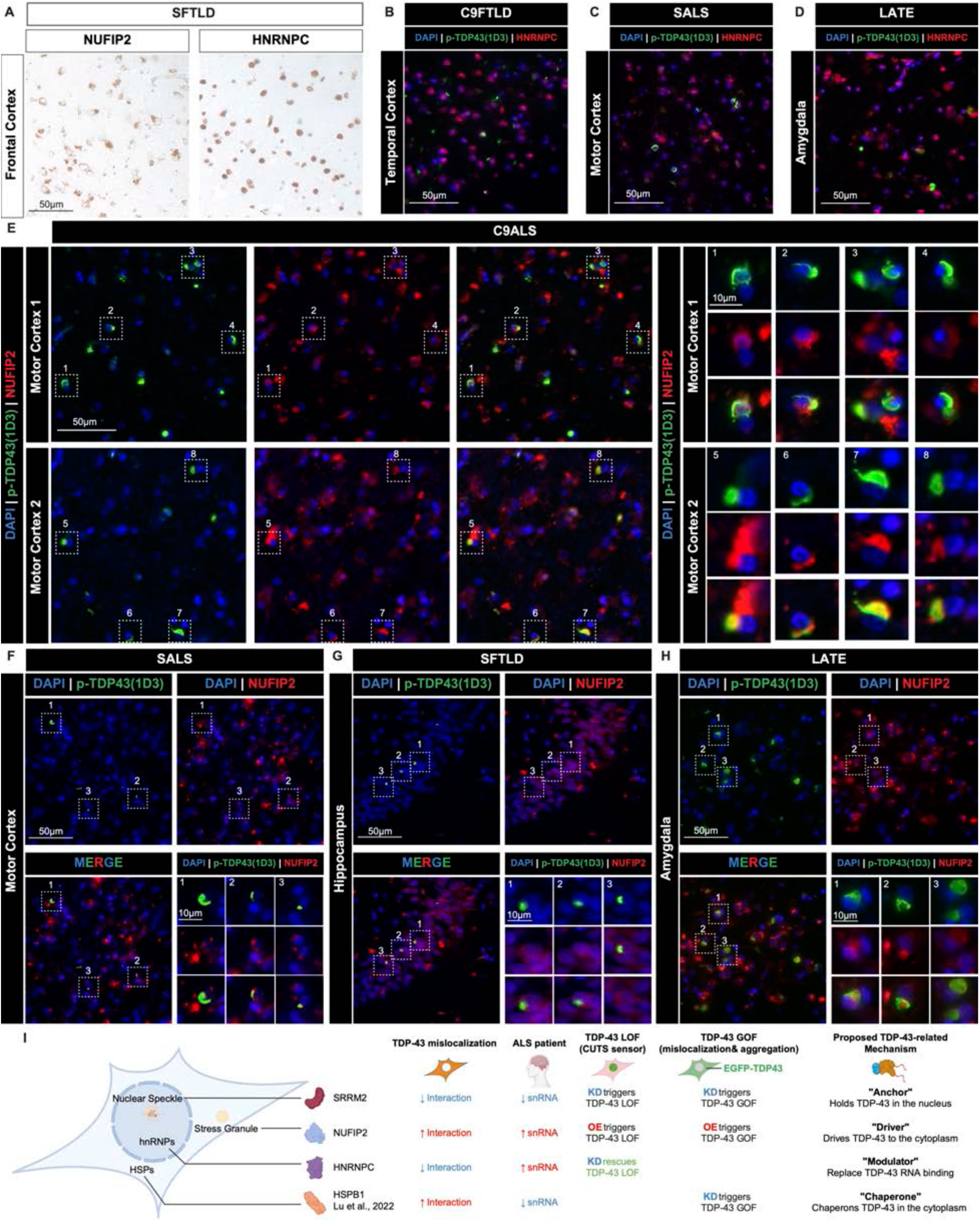
HNRNPC and NUFIP2 immunostaining in patient tissue with TDP-43 pathology. **(A)** Immunohistochemistry (IHC) staining of NUFIP2 (left) and HNRNPC (right) in the frontal cortex of a sporadic FTLD (sFTPD) patient. **(B)** Immunofluorescence (IF) staining of phosphorylated TDP-43 (p-TDP-43 using 1D3 antibody) and HNRNPC in the temporal cortex of a C9-FTLD patient. **(C)** Immunofluorescence (IF) staining of phosphorylated TDP-43 (p-TDP-43 using 1D3 antibody) and HNRNPC in the motor cortex of a sporadic ALS (sALS) patient. **(D)** Immunofluorescence (IF) staining of phosphorylated TDP-43 (p-TDP-43 using 1D3 antibody) and HNRNPC in the amygdala of a limbic-predominant age-related TDP-43 encephalopathy (LATE) patient. For (B) to (D), Blue = DAPI; Green = p-TDP-43; Red = HNRNPC; Scale bar = 50 µm. **(E)** Immunofluorescence (IF) staining of phosphorylated TDP-43 (p-TDP-43 using 1D3 antibody) and NUFIP2 in two slides of the motor cortex from a C9-ALS patient. Magnified views (right) of the region with a white square in the main image (left) represent the colocalization of p-TDP-43 and NUFIP2 in individual neurons. **(F)** Immunofluorescence (IF) staining of phosphorylated TDP-43 (p-TDP-43 using 1D3 antibody) and NUFIP2 in the motor cortex of an sALS patient. Magnified views (bottom right) of the region with a white square in the main image represent the colocalization of p-TDP-43 and NUFIP2 in individual neurons. **(G)** Immunofluorescence (IF) staining of phosphorylated TDP-43 (p-TDP-43 using 1D3 antibody) and NUFIP2 in the hippocampus of an sFTLD patient. Magnified views (bottom right) of the region with a white square in the main image represent the colocalization of p-TDP-43 and NUFIP2 in individual neurons. **(H)** Immunofluorescence (IF) staining of phosphorylated TDP-43 (p-TDP-43 using 1D3 antibody) and NUFIP2 in the amygdala of a LATE patient. Magnified views (bottom right) of the region with a white square in the main image represent the colocalization of p-TDP-43 and NUFIP2 in individual neurons. **(I)** An overall summary and mechanistic model of an interacting landscape with different forces regulating TDP-43’s functions. (i) **SRRM2** (and nuclear speckles) has a decreased interaction with TDP-43 under TDP-43 mislocalization and is also down-regulated in ALS patients. It acts as a “**Anchor**” of TDP-43 in the nucleus as the knockdown of SRRM2 triggers TDP-43 LOF (splicing defects) and GOF (aggregation and mislocalization). (ii) **NUFIP2** has an increased interaction with TDP-43 under mislocalization and is up-regulated in patients. Therefore, it plays the role of a “**Driver**” that, under overexpression, causes TDP-43 GOF and LOF. (iii) **HNRNPC** has lost interaction with TDP-43 under mislocalization but is up-regulated in ALS patients. It is proposed to be a functional “**Modulator**” of TDP-43’s splicing and thus, the knockdown of HNRNPC could partially rescue TDP-43 LOF. (iv) **HSPB1** has increased interaction with TDP-43 in the cytoplasm while down-regulated in patients. It was reported to be a “**Chaperone**” of TDP-43 and buffer TDP- 43’s aggregation ^39^. The knockdown of HSPB1 would lead to TDP-43 GOF (aggregation and mislocalization).

We employed our CUTS biosensor to functionally interrogate how each of these proteins impacts TDP-43’s splicing role^54^ and performed a preliminary semi-quantitative screen by knocking down each of the 53 selected proteins using siRNA. Following siRNA-mediated knockdown, CUTS- HEK293 cells enable the measurement of GFP intensity as a proportional indicator of TDP-43 splicing LOF, relative to a control siRNA (siControl) (schematic in Figure 5B). The resulting heatmap (Figure 5A) provides a ranked summary of the number of replicates in which each knockdown either increased (+1) or decreased (−1) in detectable GFP fluorescence compared to the siControl. This classification enabled rapid identification of putative enhancers (those that increased GFP levels) and potential suppressors of TDP-43 LOF (those that reduced GFP levels).

We validated the top enhancers (Figure 5A, left) by quantitative live-cell imaging of CUTS- HEK293 cells under single knockdown conditions (“CUTS Normal + KD”; schematic in Figure 5B). The knockdown of seven proteins—CDC40, DBR1, HNRNPA3, HNRNPL^136^, NUFIP2, PRPF31, and SRRM2—caused modest but significant GFP upregulation, indicating that loss of these factors can simulate TDP-43 LOF (Figure 5C, 5D).

To amplify and clarify their effects, we then tested a subset of hits, including the seven LOF triggers and top rescuers (Figure 5A, right) in a partial TDP-43 LOF background by co-silencing TDP-43 together with each candidate (“CUTS siTDP-43 + KD”; schematic in Figure 5E). While CDC40, HNRNPA3, and PRPF31 knockdown exacerbated the GFP increase with TDP-43 siRNA, indicating a further impaired TDP-43’s splicing function, HNRNPA0, HNRNPC, and MATR3 knockdown partially rescued it, as shown by live-cell confocal imaging and immunoblotting (Figure 5F–5I). Notably, HNRNPC, a hnRNP from Cluster 5 with significant up-regulation in ALS patients (Extended Figure 8A), displayed the most robust GFP reduction (∼90%), underscoring its potential role as a buffering or compensatory factor against TDP-43 LOF.

In parallel, we assessed whether the overexpression (OE) of selected interactors could also modulate TDP-43 splicing function in the standard CUTS background (“CUTS Normal + OE”; schematic in Figure 5J). Surprisingly, among five candidates tested (DBR1, HNRNPA0, HNRNPC, HNRNPL, and NUFIP2), NUFIP2 OE increased GFP signals, indicative of a LOF-like phenotype (Figure 5K-5M). Conversely, under partial TDP-43 knockdown (“CUTS siTDP-43 + OE”; schematic in Figure 5N), overexpressed HNRNPL mildly attenuated GFP signal (Figure 5O, 5P). This observation aligns with a recent report^136^ that HNRNPL can partially rescue cryptic exon inclusion in UNC13A under TDP-43 LOF, since the CUTS biosensor partly contains the UNC13A-CE.

Altogether, our CUTS-based screening assays revealed both positive and negative regulators of TDP-43 splicing function among the context-dependent interactors identified by our proximity labeling, functional analysis, BMC mapping, and patient-derived snRNA-seq integration. The ability of specific knockdowns or overexpression events to drive or counteract TDP-43 LOF strongly supports our central hypothesis that distinct TDP-43–binding partners, modulated by pathological cues, critically impact TDP-43’s role in mRNA splicing. Such context-dependent interactions may thus serve as potential mechanistic (Figure 6) or therapeutic (Figure 7) targets in ALS/FTLD and related disorders associated with TDP-43.

### SRRM2 and Nuclear Speckles Modulate Nuclear TDP-43 Retention and Splicing Function

Building on our previous multiomics and CUTS screening analyses, we next investigated the mechanism by which SRRM2, a top hit that induces robust TDP-43 loss of function (Figure 5C- 5D), contributes to TDP-43 pathology. SRRM2 is a key scaffold protein of nuclear speckles alongside SON ^119^, and BMC mapping indicated TDP-43 interactions with nuclear speckle components are severely diminished under TDP-43 mislocalization (Extended Figure 6A, Figure 3H-3J). Compared to SRRM2, SON KD alone did not trigger a significant LOF in the CUTS biosensor, but co-knockdown of SRRM2 and SON, with the same total siRNA concentration, significantly exacerbated the GFP increase beyond that observed with SRRM2 KD alone, indicating a synergistic effect (Extended Figure 8B, Figure 6A).

To visualize TDP-43 localization changes, we generated a stable HEK293 cell line via *Piggybac* with doxycycline-inducible expression of EGFP-TDP-43^WT^ and treated it with either siRNA-mediated KD of SRRM2, SON, or a combination (Figure 6B). In control and SON KD cells, EGFP-TDP-43^WT^ primarily remained diffuse within the nucleus. However, SRRM2 KD or double KD (SRRM2 + SON) induced a distinct population of “abnormal” cells (∼2% and ∼5% of all the live cells, respectively) marked by nuclear TDP-43 foci and cytoplasmic mislocalization (Figure 6B-6C). Immunostaining for nuclear speckles with SC-35 (SRRM2) revealed that nearly all cells harboring nuclear TDP-43 inclusions similarly exhibited disrupted nuclear speckles (>96%), underscoring the link between nuclear speckle integrity and proper TDP-43 localization (Extended Figure 8C).

Fluorescence recovery after photobleaching (FRAP) was performed to assess whether TDP-43 nuclear inclusions in the abnormal cells exhibited altered biophysical properties. Compared with diffuse nuclear TDP-43, TDP-43 inclusions within SRRM2 KD or double KD cells showed substantially reduced dynamics (Figure 6D-6E, Supplemental Video 1-2), suggesting a more solid TDP-43 structure that is consistent with pathological aggregation.

Together, these findings suggest that SRRM2-containing nuclear speckles function as a crucial mechanism for retaining nuclear TDP-43. Dysregulated SRRM2, possibly due to reduced expression in patient-derived C9-ALS neurons (Figure 4A, Extended Figure 8A), and disrupts nuclear speckles, can lead to TDP-43 mislocalization and aggregation, and ultimately resulting in TDP-43 loss of function (LOF). This mechanism highlights nuclear speckles as a vital regulatory structure that safeguards TDP-43 splicing function by maintaining its proper nuclear distribution and mobility.

### NUFIP2 Promotes TDP-43 Mislocalization and Loss of Function

NUFIP2 overexpression was identified to regulate TDP-43 splicing function (Figure 5K-5M). NUFIP2 is a member of FMR1-related proteins (others include FMR1, FXR1, and FXR2) but has not been reported to be linked to TDP-43 (Extended Figure 8D). We showed that among these four FMR1-related proteins, only NUFIP2 OE was sufficient to induce a robust increase in GFP signal, signifying TDP-43 LOF (Extended Figure 8E, 8I). This underscores the unique role of NUFIP2 in triggering TDP-43 dysfunction, aligning with its identification as a top promoter of LOF in our systematic screening.

Notably, NUFIP2-driven TDP-43 LOF is dose-dependent (Extended Figure 8F-8I). Stable expression of NUFIP2 under the EF1A promoter increased NUFIP2 protein levels marginally above endogenous amounts, whereas transient transfection of NUFIP2 cDNA under a CMV promoter produced a higher expression. Removing the 3′UTR (Δ3′UTR) from the cDNA construct further elevated NUFIP2 beyond the full-length cDNA control (Extended Figure 8F-8G). Consistent with expression levels, only CMV-NUFIP2(cDNA) and CMV-NUFIP2(Δ3′UTR) triggered increased GFP fluorescence in the CUTS reporter, with the Δ3′UTR construct causing a markedly more substantial LOF effect (Extended Figure 8H-8I). These findings highlight a direct correlation between NUFIP2 concentration and its capacity to impair TDP-43 function.

The proximity labeling data highlights a significantly increased interaction between cytoplasmic TDP-43 and NUFIP2 (Extended Figure 8A). We next examined whether this LOF phenotype was accompanied by TDP-43 mislocalization. Interestingly, cells overexpressing NUFIP2 displayed a cytoplasmic “shell-like” (ring-shaped on the cross-section) distribution for NUFIP2, within which endogenous TDP-43 localizes (Figure 6F, Extended Figure 8J, Supplemental Video 3). The appearance of “shell-like” NUFIP2 condensates and TDP-43’s mislocalization in individual cells corresponded with CUTS-GFP activation, underscoring a mechanistic link between the cytoplasmic sequestration of TDP-43 and its nuclear LOF (Figure 6G). These “shell-like” structures broaden our understanding of TDP-43 aggregation patterns and suggest that NUFIP2 actively promotes TDP-43 to accumulate in the cytoplasmic compartments and may facilitate aberrant phase transitions.

The interactome data demonstrated decreased interactions between NUFIP2 and RNA-deficient TDP-43 (Extended Figure 8A). To determine how NUFIP2-driven condensates influence TDP-43 under different RNA-binding statuses, we employed cell lines expressing cytoplasmic EGFP- TDP-43 with or without intact RNA-binding motifs (TDP-43^ΔNLS^ vs. TDP-43^5FL/ΔNLS^) (Figure 6H). In cells lacking NUFIP2 overexpression, EGFP-TDP-43^ΔNLS^ and EGFP-TDP-43^5FL/ΔNLS^ appeared predominantly diffuse or occasionally formed small puncta (Extended Figure 8K). Following NUFIP2 overexpression, EGFP-TDP-43^ΔNLS^ formed the “shell-like” structures (Figure 6I, Extended Figure 8K), whereas the EGFP-TDP-43^5FL/ΔNLS^ aggregated into “condensed” inclusions (Figure 6J, Extended Figure 8K). FRAP analysis confirmed that the NUFIP2-induced “shell-like” TDP-43 structures had intermediate dynamics, whereas the “condensed” structures by RNA- deficient TDP-43 displayed minimal signal recovery, indicative of low molecular mobility (Figure 6K-6M Extended Figure 8L, Supplemental Video 4-5).

To quantify the stability of these condensates, we examined RIPA-soluble and insoluble fractions of EGFP-TDP-43. NUFIP2 overexpression significantly raised the insoluble fraction of both TDP- 43^ΔNLS^ (∼4.3-fold increase) and TDP-43^5FL/ΔNLS^ (∼10.7-fold increase) (Figure 6N–6O). The “condensed” structures formed by the 5FL/ΔNLS mutant showed particularly lower FRAP recovery and higher insoluble transitions, mirroring their more solid-state molecular architecture. Thus, although NUFIP2 induces broad cytoplasmic mislocalization of TDP-43 and the exact phase properties and biophysical outcomes depend strongly on TDP-43’s RNA-binding status.

Collectively, our results support a working model in which NUFIP2, when elevated, sequesters TDP-43 in cytoplasmic “shell-like” compartments, thereby hindering TDP-43 nuclear localization and splicing function. If TDP-43 further loses the capacity to bind RNA, it transitions into “condensed” and low-dynamics structures (Figure 6P). Notably, NUFIP2 was significantly up- regulated in C9-ALS and sFTLD patient tissues (Figure 4A, Extended Figure 8A). These findings highlight NUFIP2’s potency in reshaping TDP-43 phase separation and emphasize its underlying connection with TDP-43 pathology in disease.

### Cis- and Trans-Rescue Strategies for TDP-43 Loss of Function: Targeted TDP-43 Replacement and HNRNPC Knockdown

In the light of our previous identification of functional TDP-43 interactors, we next sought to determine whether targeting TDP-43 itself or its regulatory partners could restore TDP-43’s physiological splicing activity. We tested two distinct strategies to rescue TDP-43 splicing function (Extended Figure 9A). The first involved a cis-targeting approach, in which endogenous TDP-43 is selectively knocked down while simultaneously replaced by a codon-optimized TDP- 43 expressed under carefully controlled conditions (“TDP-43 knockdown-replacement,” TKR). This design addresses the detrimental effects of TDP-43 overexpression by controlling exogenous TDP-43 levels to not exceed the wild-type levels, thus avoiding excessive TDP-43 accumulation, as recently demonstrated^54,56^. The second, a trans-regulatory approach, leverages our functional screening findings by modulating TDP-43’s protein partners activated by CUTS (only under TDP- 43 LOF). We prioritized CUTS-controlled HNRNPC knockdown (CCK) for proof-of-concept based on its robust capacity to rescue TDP-43 LOF events in our CUTS assays (Figure 5).

For TKR, we generated stable HEK293 cell lines harboring an shRNA targeting TARDBP mRNA and a doxycycline-inducible, codon-optimized EGFP-TDP-43 cassette (Extended Figure 9B). The inducible expression cassette was further tuned with our CUTS biosensor to mirror the autoregulatory properties of wild-type TDP-43 to avoid pathological overexpression (Extended Figure 9C–9D). We confirmed high-level silencing (∼85%) of endogenous TDP-43 and partial but significant recovery (up to 70% total TDP-43 in the best case) from the inducible EGFP-TDP-43 (Extended Figure 9E), achieving our goal of mimicking physiological TDP-43 levels.

In parallel, for the CCK strategy, we engineered stable HEK293 cells with an EGFP-NLS– shHNRNPC cassette, whose expression is activated by CUTS only upon TDP-43 LOF (Extended Figure 9F). Under siRNA-induced TDP-43 depletion, robust HNRNPC knockdown was achieved, while baseline HNRNPC expression remained largely unaffected (Extended Figure 9G–9H). Notably, CCK cells also partially restored the splicing profiles of several endogenous TDP-43 targets, including CE insertions in ATG4B, DNAJC5^137^, and HDGFL2^21^ (Extended Figure 9I). This data provided initial evidence that the trans-regulation of certain TDP-43 interactors—here, lowering HNRNPC—can buffer or compensate for TDP-43 deficiency.

To more comprehensively evaluate these two rescue approaches, we performed mRNA sequencing on the TKR and CCK stable lines, each with appropriate baseline and positive (TDP-43 LOF) controls. The TKR system was modified to have doxycycline-inducible expression of shTARDBP (Figure 7A–7B, Extended Figure 10A-10B). We defined a “rescue ratio” for each splicing junction—calculated as the fraction of LOF-induced splicing dysregulation in the positive controls that were reversed by either TKR or CCK (Figure 7C). We then curated a panel of 198 TDP-43- responsive splicing junctions based on intersections with previously published TDP-43 knockdown data in iPSC-derived neurons^20^ (Figure 7D, Extended Figure 10F-10G, Table 14). Across these events, TKR and CCK exhibited disparate rescue profiles, reflected by minimal correlation (R = −0.018, P = 0.81), suggesting distinct mechanisms (Figure 7E, Table 15). Notably, CCK rescued 154 (77.78%) events, showing a broad-acting effect against TDP-43 LOF.

We further focused on the 28 CEs from these events as a hallmark downstream readout of TDP- 43 LOF pathology (Figure 7F, Table 15 cryptic_exon=TRUE). Ranking the CE splicing rescue by TKR (left panels) revealed diverse levels of responsiveness to exogenous TDP-43, wherein events such as HDGFL2-CE were nearly fully corrected (>80% rescue), but others (e.g. EPB41L4A-CE^22^) were far less responsive. Intriguingly, CCK (right panels) also restored most of these CEs—24 out of 28 (85.71%)—to some degree, with some matching or even surpassing TKR’s efficacy. For instance, HDGFL2-CE was strongly rescued (Figure 7G, 7K), ACBD3-CE was mildly rescued (Figure 7H, Extended Figure 10C) by both TKR and CCK, whereas EPB41L4A-CE was poorly rescued by TKR but substantially corrected by CCK (Figure 7I, Extended Figure 10D). Although TKR generally could not rescue the neuron-specific STMN2-CE^18,29,138,139^ in our HEK293 models (likely due to low STMN2 expression), a partial rescue was detected with CCK, aligning with the knockdown of HNRNPC as a potentially broad-acting strategy (Figure 7J, Extended Figure 10E). Additionally, we found that STMN2 mRNA was significantly down-regulated under TDP-43 knockdown in CCK2 but was partially rescued (P = 0.05) in CCK4, which is consistent with the splicing outcome (Extended Figure 10B). Interestingly, we also detected a dramatic increase of NPTX2, a recently reported TDP-43-regulated gene highly overexpressed in both LOF cellular models and patients^140^. However, this up-regulation was not rescued by HNRNPC knockdown (Extended Figure 10B), potentially due to a different regulatory mechanism (inhibition by TDP- 43 binding to the 3’UTR) rather than splicing.

Taken together, these results reveal that (i) TKR partially restores TDP-43 function by directly replenishing TDP-43 under autoregulated control, thereby highlighting event-specific sensitivities to TDP-43 dosage, and (ii) CCK mitigates TDP-43 LOF in splicing via a mechanistically distinct route. Based on these results as well as previous research showing HNRNPC and TDP-43 have many shared binding sites^141^, we further propose that HNRNPC may act as a competitive or noncompetitive antagonist to TDP-43 at certain splicing sites (Figure 7L). Reducing HNRNPC can alleviate cryptic exon retention and other LOF-driven splicing aberrations—even when TDP- 43 remains partially depleted—thus offering a “trans-regulatory” axis of intervention.

### NUFIP2 and HNRNPC show Distinct TDP-43 Colocalization Patterns in Patient Tissue

SRRM2 pathology and nuclear speckles disruption were recently reported in C9-ALS patients, with post-mortem staining having a >60% co-occurrence with TDP-43 mislocalization in the soma of neurons ^122^. To evaluate the disease relevance of our findings on NUFIP2 and HNRNPC, we examined their expression and subcellular localization in a small cohort of post-mortem tissues from patients with TDP-43 proteinopathy (Supplementary Table 6). Both proteins were broadly distributed in the frontal cortex of sporadic FTLD-TDP cases (Figure 8A), consistent with our prior cell-based and multiomics data showing their ubiquitous presence.

Strikingly, HNRNPC remained confined to the nucleus even in neurons exhibiting cytoplasmic phosphorylated TDP-43 (p-TDP-43) inclusions (Figure 8B–D). This pattern was observed across distinct disease contexts, including the temporal cortex from a C9-FTLD patient (Figure 8B), the motor cortex from an sALS patient (Figure 8C), and the amygdala from a LATE patient (Figure 8D). These findings confirm our APEX2 interactome observations—namely that HNRNPC loses its binding to mislocalized TDP-43 in the cytoplasm but is not redistributed. Moreover, they support the notion that HNRNPC could be leveraged as a “trans-regulatory” target to ameliorate TDP-43 LOF phenotypes without altering the total TDP-43 protein levels.

In contrast, NUFIP2 colocalized with some cytoplasmic p-TDP-43 inclusions in C9-ALS motor cortex (Figure 8E), suggesting a NUFIP2–TDP-43 association in disease. This colocalization was not consistently observed in other TDP-43 proteinopathies tested, including sALS motor cortex (Figure 8F), sFTLD hippocampus (Figure 8G), or LATE amygdala (Figure 8H). The preferential accumulation of NUFIP2 in C9-ALS aligns with snRNA-seq data showing elevated NUFIP2 mRNA levels most prominently in C9-ALS patient neurons.

## Discussion

TDP-43 pathology is a hallmark of ALS/FTLD, playing a central role in neurodegeneration through its cytoplasmic mislocalization (GOF) and nuclear loss of physiological functions (LOF)^10^. Understanding the TDP-43 interactome during these events is essential for deciphering the molecular mechanisms driving pathogenesis and identifying potential therapeutic targets. By employing APEX2-based proximity proteomics in HEK293 cells, our study provides a comprehensive landscape of the TDP-43 interactome across physiological and pathological contexts, revealing key molecular programs associated with TDP-43 mislocalization, impaired RNA-binding, and oxidative stress. We mapped the dynamic changes in TDP-43 interactions and found strong concordance with various recent findings from cellular models, iPSCs, and postmortem analyses of ALS/FTLD patients^34,39,104,107,109,111,122,128,136^.

A major aspect of our findings is the context-dependent remodeling of TDP-43 interactions with various biomolecular condensates (BMCs), or membraneless organelles (MLOs), which are often formed through liquid-liquid phase separation (LLPS) and are essential for cellular organization homeostasis^37^. A recent study leveraging microscopic images of human neurons analyzed 24 organelles, including paraspeckles, nuclear speckles, stress granules (SGs), and P-bodies, and revealed their pathobiological relevance to TDP-43 pathology^142^, which is also identified in our study. Both studies reinforce that aberrant phase separation and organelle dysfunction represent fundamental mechanisms underlying TDP-43-associated neurodegeneration. Notably, consistent with recent works^31,61,128^, we also found a reduction in TDP-43:SG interactions under oxidative stress and aggregate-prone conditions (RNA deficient TDP-43).

Beyond mapping interactome shifts, we functionally investigated the consequences of TDP-43 LOF by selectively perturbing key interactomes to assess their contribution to TDP-43 cellular roles via the CUTS splicing sensor^54^,. This approach led to the identification of proteins that play essential roles in TDP-43 LOF phenotypes, shedding light on potential therapeutic targets for TDP- 43 proteinopathies. SRRM2, a key nuclear speckle scaffold, is essential for RNA processing^119–121,143^. Recent studies have linked SRRM2 dysfunction to neurodegenerative diseases, showing that in C9-ALS, SRRM2 is sequestered into poly-GR cytoplasmic inclusions, leading to nuclear speckle disruption and widespread RNA splicing defects^122^ . Our findings further demonstrate that SRRM2 or SRRM2/SON downregulation induces TDP-43 inclusions, mislocalization, and LOF, underscoring the interplay between TDP-43 and nuclear speckles. Notably, SRRM2 is downregulated in C9-ALS patients’ excitatory neurons in the motor cortex (MCX) and loses its interaction with TDP-43 under mislocalization, suggesting that SRRM2 acts as a critical nuclear “anchor” for functional TDP-43 (Figure 8G). This supports the idea that SRRM2 disruption is upstream of TDP-43 pathology, aligning with evidence that poly-GR triggers TDP-43 dysfunction in C9-ALS^144^. Beyond ALS/FTLD, nuclear speckle dysfunction may contribute to other neurodegenerative diseases. Tau pathology was previously reported to be a driver of nuclear speckle component mislocalization^145^, suggesting that SRRM2 dysregulation could also contribute to Alzheimer’s disease (AD) and related disorders. This broader impact of nuclear speckle dysfunction raises the intriguing possibility of shared pathogenic mechanisms, especially TDP-43 dysfunction, across neurodegenerative diseases^146–149^.

Our study identifies NUFIP2 overexpression as a potent trigger of TDP-43 mislocalization in cell models, with colocalization between NUFIP2 and TDP-43 in C9-ALS patient MCX tissues exhibiting TDP-43 mislocalization. These findings suggest that NUFIP2 may sequester TDP-43, driving its cytoplasmic mislocalization and subsequent LOF. Notably, NUFIP2 RNA is elevated in patient tissue, and like other stress granule components, it shows increased TDP-43 interaction when TDP-43 is mislocalized, suggesting that NUFIP2 may act as a potential “driver” of TDP-43 pathology (Figure 8G). Intriguingly, NUFIP2 knockdown in the CUTS biosensor system also triggers TDP-43 LOF (Figure 5C-5D), indicating the possibility of a dual effect on TDP-43 function, which raises the need for future investigation. In addition, NUFIP2’s role in post-transcriptional regulation of mRNA through the NMD pathway adds another layer to its involvement in cellular stress responses. Previous findings suggest that NUFIP2 is a cofactor for Roquin-mediated mRNA decay^150^, linking its role in RNA quality control to TDP-43 pathology. This pathway has recently been highlighted for its role in ALS/FTLD^41,151–153^. Further exploration of NUFIP2 neuropathology across disease subtypes, and its interaction with components of the NMD machinery could reveal insights into TDP-43 mislocalization and dysfunction, which may advance our understanding of the ALS/FTLD pathogenesis.

HNRNPC is upregulated in sALS and C9-ALS patients’ MCX excitatory neurons alongside other hnRNPs and loses its interaction with TDP-43 when TDP-43 is mislocalized, coinciding with impaired RNA binding of TDP-43. We thus propose HNRNPC as a potential “modulator” of TDP- 43 splicing function (Figure 8G). Using the CUTS sensor assay, we demonstrate that HNRNPC knockdown consistently rescues TDP-43 LOF phenotypes. Furthermore, RNAseq and splicing event analyses indicate the rescue of TDP-43 cryptic exon inclusion upon HNRNPC KD. Notably, these rescued cryptic exon events were distinct from those observed in TDP-43 replacement experiments, suggesting that HNRNPC mediates TDP-43 LOF rescue through different but essential mechanisms. Previous studies on the hnRNP family have linked their various functions in neurodegenerative diseases, with the knockdown of several hnRNP orthologs, including HNRNPC, alleviating TDP-43 (TBPH) toxicity in *Drosophila* models^154^, which aligns with our findings. However, to understand the specific impact of HNRNPC on splicing regulation, future studies should map HNRNPC and TDP-43 RNA binding sites under normal conditions or knocking down both proteins, as well as characterize HNRNPC-driven splicing events independent of TDP-43. This could provide insights into how alternative RNA processing pathways emerge in the context of TDP-43 pathology.

Recent studies identified HSPB1 as a molecular chaperone of TDP-43, with its downregulation leading to TDP-43 inclusions and mislocalization—suggesting a broader role for heat shock proteins in maintaining TDP-43 homeostasis^39^. This matches with our multi-omics analysis that HSPB1 is downregulated in ALS/FTLD patient samples while gaining interaction with mislocalized TDP-43 under oxidative stress (Figure 8G). Thus, by integrating the key examples of SRRM2, NUFIP2, HNRNPC, and HSPB1, our study uncovers a complex interacting landscape governing TDP-43’s function. With robust evidence from proteomic shifts, patient-derived datasets, and functional screening, we present a comprehensive framework for understanding how diverse cellular machineries converge on TDP-43 pathology (Figure 8G). This regulatory landscape deepens our insight into ALS/FTLD pathogenesis and opens new avenues for therapeutic intervention, targeting these distinct pathways to restore TDP-43 homeostasis.

Moreover, emerging evidence indicates that certain TDP-43 protein interactors, such as G3BP1 in SGs, are likewise regulated by TDP-43 at the mRNA level^155^. This feedback mechanism, reminiscent of TDP-43’s own autoregulation^156^, underline TDP-43’s multifaceted regulatory network spanning both RNA and protein interactions. In our datasets, we identified a subset of genes whose splicing is regulated by TDP-43 and that have the potential to interact with TDP-43 at the protein level in a context-dependent manner, including BBC3 (encoding the PUMA protein)^157^, PFKP^158^, LENG8^159^, and ACBD3^160,161^, all serving critical cellular functions related to neurodegenerative disorders (Extended Figure 10H). Further investigations to dissect these intricate interactions will likely yield deeper insights into TDP–43–mediated pathobiology.

The development for ALS/FTLD therapies remains challenging due to the lack of clear genetic markers and late-stage clinical presentation^162–164^, highlighting the need for targeted therapeutic strategies. This study identifies key modulators of TDP-43 pathology and demonstrates two LOF rescue strategies: (i) finely tuning TDP-43 interactors, such as CUTS-controlled HNRNPC knockdown (CCK), and (ii) knocking down endogenous mutant TDP-43 while introducing a biosensor-controlled exogenous TDP-43 (TKR). Both approaches restored TDP-43-specific splicing events through distinct mechanisms. Our datasets, together with our findings, provide a valuable resource as well as a framework for identifying TDP-43 modulators, and lay the foundation for translational applications.

## Methods

### Plasmid generation

The plasmid information is listed in Supplementary Table 1. All the vectors, unless specified from other resources, were generated by NEBuilder HiFi DNA Assembly Master Mix (NEB, E2621L) following the manufacturer’s protocol. All the plasmids were verified using whole-plasmid sequencing via Oxford Nanopore provided by Plasmidsaurus.

### Cell culture and plasmid transfection

HEK293 cells (female, purchased from ATCC), Hela TDP-43 knock-out (KO) cells (a kind gift from Dr. Shawn M Ferguson^58^ were maintained in DMEM (Thermo Fisher Scientific) supplemented with 10% HyClone Bovine Growth Serum (GE Healthcare Life Sciences) and 1x GlutaMAX (Thermo Fisher Scientific) at 37°C and 5% CO2, with a humidified atmosphere. Cells were seeded onto coverslips or plates coated with collagen (50 mg/mL, GIBCO) and allowed to incubate overnight prior to transfections using Lipofectamine 3000 (Invitrogen, L3000015) with a proper amount of DNA performed according to manufacturer’s instructions.

### *Piggybac* stable cell line generation

HEK293 cells were seeded onto 6-well plates 24h before transfection and were grown to about 70% confluence. 2.5ug of plasmid with *Piggybac* backbone was co-transfected with 0.5ug Super *PiggyBac* Transposase (PB200PA-1) plasmid respectively to each well using Lipofectamine 3000 (Invitrogen, L3000015) according to manufacturer’s instructions. One well without transposase was also conducted as the negative control. The media was changed 48h after transfection into selection media with puromycin (Sigma, P8833) to a final concentration of 5 μg/mL, or hygromycin B (Gibco, 10687010) to a final concentration of 500 μg/mL. The selection media was then refreshed every 2 days. The control cells were all died in about 5 days. The surviving cell lines were continuously cultured after selection and passaged for 2–3 generations with a reduced concentration of puromycin (2.5 μg/mL) or hygromycin B (250 μg/mL) to obtain stable cell lines.

The induced expression of desired constructs was verified via Immunofluorescence staining or Western blotting.

### APEX2-medicated proximity labeling

Stable HEK293 cell lines expressing 3xFlag-APEX2-TDP-43 (WT, ΔNLS, 5FL, 5FL+ΔNLS), 3xFlag-APEX2-3XNES, and 3xFlag-APEX2-3XNLS were split into 24-well plates containing pre-coated coverslips (for Immunofluorescence staining), 6-well plates (for Western blotting), or 10-cm dishes (for Mass Spectrum) 24h before induction. Doxycycline (Sigma, D9891) was added into the media to a final concentration of 1 μg/mL to induce the expression of APEX2 constructs. For transient transfected experiments, wildtype HEK293 cells were transfected with pCMV- 3xFlag-APEX2-TDP-43 (WT, ΔNLS, 5FL, 5FL+ΔNLS) plasmids using Lipofectamine 3000. After 48h of expression (for transfection, 24h), the plates were replaced with fresh media containing 500 μM of biotin phenol (ApexBio, 41994-02-9) and were incubated for 30 min. The labeling was initiated by adding H2O2 (Sigma, 216763) to reach a final concentration of 1 mM for precisely 1 min. The reaction was quenched by immediately adding freshly prepared 2X quenching buffer (10 mM Trolox (Sigma, 238813), 20 mM sodium ascorbate (Sigma, A4034), and 20 mM sodium azide (Fisher Scientific, BP922I) in pre-chilled DPBS (Thermo Scientific, 14190235)) of the same volume as the media, and rinsing two more times with 1X quenching solution (50% 2X quenching buffer diluted with pre-chilled DPBS). Negative controls without biotin phenol (BP), H2O2, or doxycycline (Dox) were also included in the validation tests. The cells were further proceeded according to different experimental purposes.

### Cell stress treatment

Stressors’ information is listed in Supplementary Table 2. All the stressors were mixed with the media to desired final concentration and added to cells via media changing. For recovery, the cells were washed one time with fresh media and then changed into recovery media.

### RNA Oligonucleotides

RNA oligonucleotides ([AC]17 and [UG]17) were ordered from Horizon Discovery, HPLC purified, fully 2′OMe modified.

### SDS-PAGE and Western blotting

Cells were lysed on-plate with pre-chilled RIPA buffer (Boston Bioproducts, BP-115X) with 1% protease inhibitor (PI) cocktail (Sigma, P8340) on ice for 10 min and then sonicated at 20% power by a sonicator (Brason) for 3 times, with each time 15 s in an ice bath and 2 min of cooling between each sonication. Protein concentrations were measured using the Pierce BCA Protein Assay Kit (Thermo Scientific, 23227). The samples were then denatured by adding 4X Laemmli Sample Buffer (Biorad, 1610747) and incubating at 95 °C for 10 min before Sodium Dodecyl Sulfate PolyAcrylamide Gel Electrophoresis (SDS-PAGE). The denatured samples were separated by SDS-PAGE using 4%–20% Mini-PROTEAN TGX Precast Gels (Biorad, 4561096) and transferred to 0.45 μm Nitrocellulose Membranes (Biorad, 1620146) using Mini Gel Tank and Blot Module Set (Invitrogen, NW2000). Total protein levels were measured by Ponceau S dye (ThermoFisher A40000279). Following water and TBS washes, membranes were blocked in 5% non-fat milk (Thermo Scientific, 50-488-786) in TBS-T (0.1% Tween 20 (Fisher Scientific, BP337) in TBS) and incubated with primary antibodies in the blocking buffer at 4 °C overnight with mild shaking. The primary antibodies and their dilution ratios were as follows: mouse-anti-FLAG (Sigma, F1804), mouse-anti-GAPDH (Cell Signaling, 2118), mouse-anti-GFP (Santa Cruz, sc- 9996), mouse-anti-TDP-43 (Proteintech, 60019-2-Ig), rabbit-anti-TDP-43 (Proteintech, 10782-2- AP), rabbit-anti-mCherry (Cell Signaling, 43590), rabbit-anti-DBR1 (Proteintech, 16019-1-AP), rabbit anti-NUFIP2 (Proteintech, 17752-1-AP), rabbit-anti-HNRNPC (Proteintech, 11760-1-AP), rabbit-anti-HNRNPL (Proteintech, 18354-1-AP). After incubation, membranes were washed thrice with TBS-T and incubated with Donkey-anti-mouse (JacksonImmunoResearch, 715035151) or Donkey-anti-rabbit (JacksonImmunoResearch, 711035152) secondary antibodies conjugated with HRP, or Streptavidin-HRP (EMD Millipore, OR03L) at room temperature for 1h. After washing again with TBS-T for three times, the membranes were visualized using Western Lightning ECL Pro (Revvity, NEL1201001EA) or Supersignal West Femto Maximum Sensitivity Chemiluminescent Substrate (Thermo Scientific, 34095) in an Amersham ImageQuant 800 GxP biomolecular imager system (Amersham, 29653452). Western blotting results were quantified using Fiji ImageJ (V2.3.1) software.

### RIPA soluble/insoluble (sol/insol) fractionation

HEK293 cells were processed for fractionation directly from culture. Briefly, cell pellets scraped with pre-chilled PBS were transferred into 1.5 mL microcentrifuge tubes. The samples were centrifuged at 500× g for 5 min at 4 °C to pellet the cells. The supernatant was then carefully removed, and each pellet was resuspended in pre-chilled RIPA buffer supplemented with PI. The suspensions were incubated on ice for 10 min to facilitate complete lysis. Following lysis, the samples were centrifuged at 17,000 × g for 10 min at 4 °C. The supernatants, which constitute the RIPA-soluble fraction, were collected and stored at −20 °C until further analysis. The remaining insoluble material was further processed by washing. Each pellet was resuspended in the same RIPA buffer (with PI) and centrifuged at 17,000 × g for 5 min at 4 °C. This wash was repeated for a total of two washes, with the supernatant discarded after each centrifugation. Thereafter, the insoluble pellets were resuspended in a modified RIPA buffer containing 4 M urea, PI, and Benzonase (Sigma, E1014-25KU). The resuspended pellets were then agitated at 1350 rpm for 10 min on a thermomixer (Eppendorf) at 25 °C to promote extraction of the remaining proteins. A single 10-second sonication pulse at 20% power was applied to fully dissolve the pellets and acquire the final insoluble fraction. Both soluble and insoluble fractions were subsequently analyzed by SDS-PAGE and western blotting, as described above.

### Immunofluorescence staining

Cells on coverslips were fixed with 4% PFA (Electron Microscopy Sciences, 15714-S) atroom temperature for 20 min. The cells were then washed with PBS (Fisher Scientific, 20012050) and blocked with 5% Normal Donkey Serum (Jackson ImmunoResearch, 017-000-121) in PBS- T (0.1% Tween 20 in PBS). The cells were incubated with primary antibodies in the blocking buffer at 4 °C overnight. The primary antibodies and their dilution ratios were as follows: mouse-anti-FLAG (Sigma, F1804), mouse-anti-G3BP1 (Santa Cruz, sc-365338), mouse-anti-TDP-43 (Proteintech, 60019-2-Ig), mouse-anti-SC35(SRRM2) (Abcam, ab11826), rabbit-anti-TDP-43 (Proteintech, 10782-2-AP), rabbit anti-ATXN2 (Proteintech, 21776-1-AP), rabbit anti-NUFIP2 (Proteintech, 17752-1-AP). After incubation, cells were washed thrice with PBS-T and incubated with secondary antibodies, Streptavidin, or Hoechst at room temperature for 1h as follows: DyLight 405 AffiniPure Donkey Anti-Mouse IgG (H+L) (Jackson ImmunoResearch, 715-475- 151), Alexa Fluor 488 Donkey anti-mouse IgG (H+L) (Jackson ImmunoResearch, 715-545-150), Alexa Fluor 488 Donkey anti-rabbit IgG (H+L) (Jackson ImmunoResearch, 711-545-152), Alexa Fluor 594 Donkey anti-mouse IgG (H+L) (Jackson ImmunoResearch, 715-585-151), Alexa Fluor 594 Donkey anti-rabbit IgG (H+L) (Jackson ImmunoResearch, 711-585-152), Alexa Fluor 647 Donkey anti-mouse IgG (H+L) (Jackson ImmunoResearch, 715-605-150), Alexa Fluor 647 Donkey anti-rabbit IgG (H+L) (Jackson ImmunoResearch, 711-605-152), Cy3-Streptavidin (Jackson ImmunoResearch, 016-160-084), Cy5-Streptavidin (Jackson ImmunoResearch, 016- 170-084), and Hoechst 33258 solution (Sigma, 94403). Cells were washed two times in PBS-T and one time in PBS. The coverslips were mounted onto slides with Vibrance Antifade Mounting Medium (Vector Laboratories, H-1700) for 24 h away from light. Then they were proceeded to be visualized by confocal microscopy.

### Confocal microscopy

All confocal experiments were performed on a Nikon A1 laser-scanning confocal microscope system. For regular confocal microscopy, CFI Plan Apo Lambda 60X Oil immersion objective (Nikon) was used. For live-cell imaging, a Tokai HIT stage-top incubator preheated (37°C and 5% CO2) for 10 min prior to imaging, and CFI Plan Apo Lambda 10X or 60X (Oil) objectives (Nikon) were used. Nikon Elements imaging software was used to control the microscope and perform visualization. The images presented were representatives of at least two independent experiments with three or more biological replicates per experiment. The data presented are representatives of at least two independent experiments utilizing three or more replicates per experiment.

### Fluorescence recovery after photobleaching (FRAP)

FRAP imaging was performed on HEK293 cell lines cultured in DMEM containing 10% FBS on 24-well glass-bottom plates. Where indicated, cells expressing EGFP-TDP-43 variants were induced for 48 hours by adding doxycycline (1 μg/mL) to the medium. Using a Nikon A1 laser-scanning confocal microscope with a 60× oil-immersion objective, granules or inclusions were located, and circular regions of interest (ROIs) with a diameter of roughly 2 μm were photobleached for 500 ms at 50% laser power (488 nm line). For each experiment, 2–5 baseline images were examined before bleaching, and fluorescence recovery was monitored for up to 3 minutes at 5-second intervals. Nikon NIS-Elements software was used to quantify recovery curves, normalizing the fluorescence intensity in the bleached region to its pre-bleach level and setting the intensity immediately after bleaching to zero. At least 6–10 bleached structures per condition were analyzed to calculate averages and standard deviations. The data presented are representatives of at least 5 independent images, from at least two biological replicates.

### Human post-mortem tissues

All patient post-mortem tissue were provided by the Center for Neurodegenerative Disease Research (CNDR) at the University of Pennsylvania and information is listed in Supplementary Table 5.

### IHC and IF staining in post-mortem tissue

Primary antibodies included NUFIP2 (Proteintech 17751-1-AP), HNRNPC (Proteintech 11760-1- AP), and 1D3 (pTDP-43, a gift of Elisabeth Kremmer and Manuela Neumann). Sections were subject to microwave antigen retrieval and stained with primary antibody overnight. For immunohistochemistry, NUFIP2 was diluted 1:250, HNRNPC was diluted 1:2000, and 1D3 was diluted 1:400; the next day, biotinylated secondary antibodies were added, followed by an ABC kit (Vectorlabs) and visualized using immPACT DAB (Vectorlabs). For immunofluorescence, NUFIP2 was diluted 1:200, HNRNPC was diluted 1:1000, and 1D3 was diluted 1:200; the next day, fluorescently labeled secondary antibodies (Invitrogen Alexa Fluor 488G and Alexa Fluor 594R) were added, followed by DAPI Fluoromount-G (SouthernBiotech). Images were captured on a Nikon Eclipse Ni microscope.

### siRNA transfection

The siRNA (siGenome Smatpool siRNA, Dharmaco) information for CUTS screening and following experiments is listed in Supplementary Table 3. siGENOME non-Targeting siRNA for control (Dharmaco, D-001206-13-05) was used as control (siControl). The indicated amount (usually to the final concentration of 20 nM) in each experiment was transfected into HEK 293 cells using Lipofectamine RNAiMAX reagent (Invitrogen, 13778150) according to the manufacturer’s protocol. For TDP-43 knockdown, the following siRNA was used: ON-TARGETplus SMARTpool siRNA against TARDBP (Dharmaco, L-012394-00-0005).

### cDNA and ORF plasmids

The cDNA and ORF-expressing plasmids information for CUTS screening and following experiments is listed in Supplementary Table 4.

### Streptavidin pull-down of biotinylated proteins

For the enrichment of biotinylated proteins, 15% of the cell lysates from one 10-cm dish were diluted five times with RIPA buffer and incubated with 50 μL NanoLINK Streptavidin Magnetic Beads (TriLink Biotechnologies, M-1002) overnight at 4℃ on a rotator. Beads were collected against a magnetic stand, and the supernatant was set aside for future analysis (termed flow- through). A total of eight washes were performed prior to on-beads digestion: (1) RIPA; (2) 50% RIPA, 50% UWB; (3) UWB (20 mM Tris-HCl pH7.6 (Rockland, MB-003), 150 mM NaCl (Invitrogen, AM9759), 2M Urea (Fisher Scientific, M-13269)); (4) 50% UWB, 50% RIPA; (5) RIPA; (6) ABC (50mM pH=8.0 ammonium bicarbonate (Sigma, A6141)); (7) ABC; (8) ABC. For each time of the washes, the beads were completely resuspended by vortex and rotated end-over-end on a rotator for 5 min prior to magnetic separation. Before the last spin, 10% of the beads were saved for quality control.

### On-beads digestion

Roughly 2.5 ug of Sequencing Grade Modified Trypsin (Promega, V5113) was added to the bead in ABC together with 1mM DTT (Sigma, 43816) to increase the digesting efficiency^165^. The samples were then digested at 35℃ overnight on a thermomixer (Eppendorf) shaking at 1000 rpm. After digestion the supernatant was removed and the beads were washed once with enough ABC to cover. After 10 minutes of gentle shaking, the wash was removed and combined with the initial supernatant. The peptide extracts were lyophilized, then resuspended in 30 uL 0.1% trifluoroacetic acid. A small portion of the extract is used for fluorometric peptide quantification (Thermo Scientific Pierce).

### Label-free DIA LC-MS

For each sample, 500 ng total peptide was loaded onto a disposable Evotip C18 trap column (Evosep Biosystems, Denmark) and subjected to nanoLC on an Evosep One instrument (Evosep Biosystems). Tips were eluted directly onto a PepSep analytical column, dimensions: 150umx25cm C18 column (PepSep, Denmark) with 1.5 μm particle size (100 Å pores) (Bruker Daltronics). Mobile phases A and B were water with 0.1% formic acid (v/v) and 80/20/0.1% ACN/water/formic acid (v/v/vol), respectively. The standard pre-set method of 100 samples-per-day was used, which is a 14 min run. The mass Spectrometry was done on a hybrid trapped ion mobility spectrometry-quadrupole time of flight mass spectrometer (*timsTOF HT*, (Bruker Daltonics, Bremen, Germany), operated in PASEF mode. The acquisition mode was Data-independent analysis (DIA). The acquisition scheme used for DIA consisted of 18 precursor windows at a width of 50m/z per cycle. The TIMS scans layer the doubly and triply charged peptides over an ion mobility, 1/k0 range of 0.60-1.4 V*sec/cm^2^. Precursor windows began at 360 m/z and continued to 1200 m/z. The collision energy was ramped linearly as a function of the mobility from 63 eV at 1/K0=1.4 to 17 eV at 1/K0=0.6.

### MS data processing

Raw files were processed with Spectronaut version 18 (Biognosys, Zurich, Switzerland) using DirectDIA analysis mode. Mass tolerance/accuracy for precursor and fragment identification was set to default settings. The reviewed FASTA for Homo Sapiens, UP000005640 downloaded from Uniprot (on 11 Sep 2023) and a database of 112 common laboratory contaminants (https://www.thegpm.org/crap/) were used. A maximum of two missing cleavages were allowed, the required minimum peptide sequence length was 7 amino acids, and the peptide mass was limited to a maximum of 4600 Da. Carbamidomethylation of cysteine residues was set as a fixed modification, and methionine oxidation and acetylation of protein N termini as variable modifications. A decoy false discovery rate (FDR) at less than 1% for peptide spectrum matches and protein group identifications was used for spectra filtering (Spectronaut default). Decoy database hits, proteins identified as potential contaminants, and proteins identified exclusively by one site modification were excluded from further analysis.

### Post-MS data processing

7487 proteins were obtained from mass spectrometry and protein intensity values were log-transformed. Proteins with more than 50% missing values or mass spectrometry detection p value > 0.05 were removed, and a total of 6592 proteins were kept. We used impute.knn with default parameters from R package impute (DOI: 10.18129/B9.bioc.impute) for missing data imputation.

### Definition of TDP-43 interactome and differentially enriched TDP-43 interactors

To define the TDP-43 interactome under each experimental condition, we used nuclear export signal (NES) as a control for cytoplasm-localized TDP-43 and nuclear localization signal (NLS) as a control for nucleus-localized TDP-43. Differential interactors were identified by performing a t-test on log-transformed protein intensities between experimental groups and their respective controls. Proteins with log2-fold change (LFC) > 0.3 and p-value < 0.05 were considered part of the TDP-43 interactome. The resulting interactome lists were used as input for Gene Ontology (GO) term enrichment analysis using DAVID GO (Database for Annotation, Visualization, and Integrated Discovery), with default parameters and an FDR-corrected significance threshold of q < 0.05.

To define differential TDP-43 interactors between specific experimental conditions, we performed t-tests on log-transformed protein intensities between experimental groups and their matched controls. For comparisons within the same cellular compartment (both nuclear or both cytoplasmic), interactors were defined using LFC > 0.3 and p-value < 0.05. For comparisons across cellular compartments (nuclear vs. cytoplasmic), interactors were defined using LFC > 1 and q-value < 0.05 to account for larger expression differences. The lists of differential interactors were also analyzed for GO term enrichment using DAVID GO, with default parameters and an FDR-corrected significance threshold of q < 0.05.

To identify interactome alterations associated with RNA-binding impairment, TDP-43 mislocalization, and oxidative stress treatment respectively, we performed differential analysis using the limma package. Each experimental condition was modeled as an independent factor while adjusting for potential confounding effects from the other two conditions. Specifically, three separate linear models were fitted:

1. RNA-binding impairment analysis: Differential expression was assessed between RNA-binding deficient and RNA-binding competent groups while controlling for mislocalization and oxidative stress treatment.
2. TDP-43 mislocalization analysis: Cytoplasmic and nuclear TDP-43 conditions were compared, adjusting for RNA-binding impairment and oxidative stress.
3. Oxidative stress treatment analysis: Cells treated with sodium arsenite were compared to untreated controls, controlling for TDP-43 mislocalization and RNA-binding impairment.

For each contrast, a design matrix incorporating the focal variable and the additional covariates was constructed, and differential expression was computed using Limma’s empirical Bayes framework. Significantly altered genes were identified using moderated t-statistics and ranked based on adjusted p-values.

### Normalization of LFCs for cross-comparison analysis

To enable direct comparison across nine differential interactome analysis as well as differential expressed genes (DEG) from ALS/FLTD snRNA-seq, we adapted the normalization method from the snRNA-seq paper^55^. We standardized the magnitude of LFCs by converting them into Z-scores. For each differential comparison, LFC values were transformed using the R function scale(x, center = 0, scale = sd(x)), where x represents the vector of LFCs for that specific comparison. This transformation ensures that the LFC values are centered at 0 while normalizing their magnitude relative to the variability within each dataset. By applying this Z-score transformation, we accounted for differences in dynamic range across conditions and facilitated a direct comparison of relative effect sizes across all nine differential analyses.

### Identification and clustering of common significant genes

To identify common significant genes between our differential interactome analysis and patient-derived snRNA-seq data, we applied distinct selection criteria. RNA-seq differentially expressed genes (DEGs) were identified using thresholds from the original literature: |Z-score| > 1 and FDR-adjusted p-value < 0.05. For differential interactome analysis under mislocalized, oxidative stress (AS), and impaired RNA-binding conditions, a gene was considered significant if at least one of the three conditions showed p < 0.05 (t-test). The final common significant gene set was defined by the intersection of significant interactome genes across conditions with the RNA-seq DEGs.

To explore gene expression patterns, we performed hierarchical clustering using the pheatmap package in R. Both genes and conditions were clustered (cluster_rows = TRUE, cluster_cols = TRUE) with default settings. To refine functional classification, we applied cutree_rows = 15, partitioning the genes into 15 clusters. The resulting clusters were subsequently used as input for GO term enrichment and correlation analysis.

### RNA extraction

Total RNA was extracted with RNeasy Mini Kit (Qiagen, 74106) following the manufacturer’s protocol. The concentration of extracted RNA was measured by Nanodrop Spectrophotometer (Nanodrop, ND-1000). For mRNA sequencing, the RNA was accurately measured by Qubit RNA HS Assay Kit (Invitrogen, Q32852).

### RT-PCR

The extracted total RNA was reverse-transcripted into cDNA using Iscript Reverse Transcription Supermix (Biorad, 1708841) following the manufacturer’s instructions. Primers information is listed in Supplementary Table 5. PCR products were separated by agarose gel electrophoresis, and the bands were visualized with Amersham ImageQuant 800 GxP biomolecular imager system.

### mRNA sequencing and DEG analysis

mRNA-seq libraries were generated using the Illumina Stranded mRNA Prep Kit (Illumina, 20040534), following the protocol provided by the manufacturer. Library concentration and integrity were then evaluated with a Qubit fluorometer and an Agilent TapeStation. Subsequent sequencing was carried out on an Illumina NextSeq 2000 P3 platform, producing 100 bp paired-end reads. Each experiment was performed with three biological replicates, and each sample yielded approximately 50–60 million raw reads.

For a quick DEG analysis, the RNA was processed using the Salmon pipeline^166^ with the default setting and quantified using the GRCh38.p13 human genome build with an Ensembl version of the Gencode V45 transcripts. The processed data were in the form of TPM (transcript per million) for mRNAs. For comparisons against the CUTS-only (TKR1) group, replicates #2 was excluded due to being an outlier in PCA. One-way ANOVA or two-way ANOVA followed by a Tukey’s test was conducted when data underwent normality statistical comparisons between different groups.

### Sequencing data processing and alignment for splicing analysis

Trimmed reads were aligned to the GRCh38.p13 human genome build with Gencode V38 gene models using STAR (v2.7.5a). Samples were aligned using ENCODE options as described in the STAR manual^167^, as well as two-pass mapping (--twoPassMode Basic). Gene-level counts were quantified with featureCounts (v2.0.1), and sample clustering was manually checked with DESeq2 (v1.44.0) by using the vst() and plotPCA() functions.

### Splicing analysis with MAJIQ

Aligned reads were analyzed for group-level splicing alterations using MAJIQ/VOILA (v2.4.dev102+g2cae150)^168,169^. Briefly, all samples were used to build a database of splicing events with *majiq build,* using the flags --min-intronic-cov 1, --simplify, and default settings otherwise. Splicing changes between groups were analyzed with *majiq deltapsi* with default settings. For comparisons against the CUTS-only (TKR1) group, replicates #2 was excluded due to being an outlier in PCA. Results tables were created with *voila tsv* and parsed using in-house R code. Any splice junction with |ΔPSI|>0.1 and probability of changing >0.95 between conditions was considered a significant differential splicing event.

### Statistical Analysis

Unless specified, quantitative data was analyzed using GraphPad Prism 10.0 software. The specific statistical significance analysis methods, p-value indicators, and number of replicates in each experiment are detailed in the respective figure legends.

## Acknowledgements

We thank UC Davis Proteomics Core Facility for their DIA-LC-MS service. We thank the Health Sciences Sequencing Core at UPMC Children’s Hospital of Pittsburgh (HSSC) for their sequencing service. We thank Dr. Shawn M Ferguson, Ph.D. (Yale University School of Medicine) for kindly providing the TARDBP^-/-^ HeLa cell lines used in this study. We thank Olivia R. Shapiro (Donnelly Lab, University of Pittsburgh School of Medicine) for their assistance in the early portion of these studies. We thank Yuxi Ding and Xiaochen Li (School of Medicine, Tsinghua University) for their helpful discussions. Some figure subpanels were created with BioRender.com.

## Author contributions

Conceptualization: LX, YZ, BTH, & CJD

Data curation: LX, YZ, BTH, JCM, MW, JLR, MN, GG, EBL, EK, MC, & CJD

Formal analysis: LX, YZ, BTH, JCM, MW, EEB, MN, CAB, JX, JM, GG, EBL, EK, MC, & CJD

Funding acquisition: LX, YZ, EBL, EK, MC, CJD Investigation: LX, YZ, BTH, JCM, MW, JLR, MN, GG, & FL

Methodology: LX, YZ, BTH, JCM, MW, JLR, MN, GG, AMG, EBL, FL, WOR, EK, MC, & CJD

Project administration: LX, YZ, & CJD

Resources: LX, YZ, BTH, JCM, MW, JLR, MN, GG, EBL, EK, MC, & CJD

Supervision: LX, YZ, WOR, WK, MC, & CJD

Validation: LX, YZ, JCM, MW, JLR, EEB, MN, GG, EBL, FL, WOR, EK, MC, & CJD Visualization: LX, YZ, BTH, JCM, MW, EEB, CAB, GG, MC, & CJD

Writing – original draft: LX, YZ, & CJD

Writing – review & editing: LX, YZ, BTH, JCM, MW, JLR, EEB, MN, CAB, JX, JM, GG, AMG, FL, WOR, EBL, EK, MC, & CJD

## Data and materials availability

The plasmids developed in this study will be freely accessible to the scientific community. All distribution requests will be fulfilled promptly. Material transfers will be conducted under terms no more restrictive than those outlined in the Simple Letter Agreement or the Uniform Biological Materials Transfer Agreement without any reach-through conditions. The lead contact will provide the data and code presented in this paper upon request. Any supplementary details needed for data reanalysis can be obtained from the lead contact upon request.

---

**Extended Figure 1:**
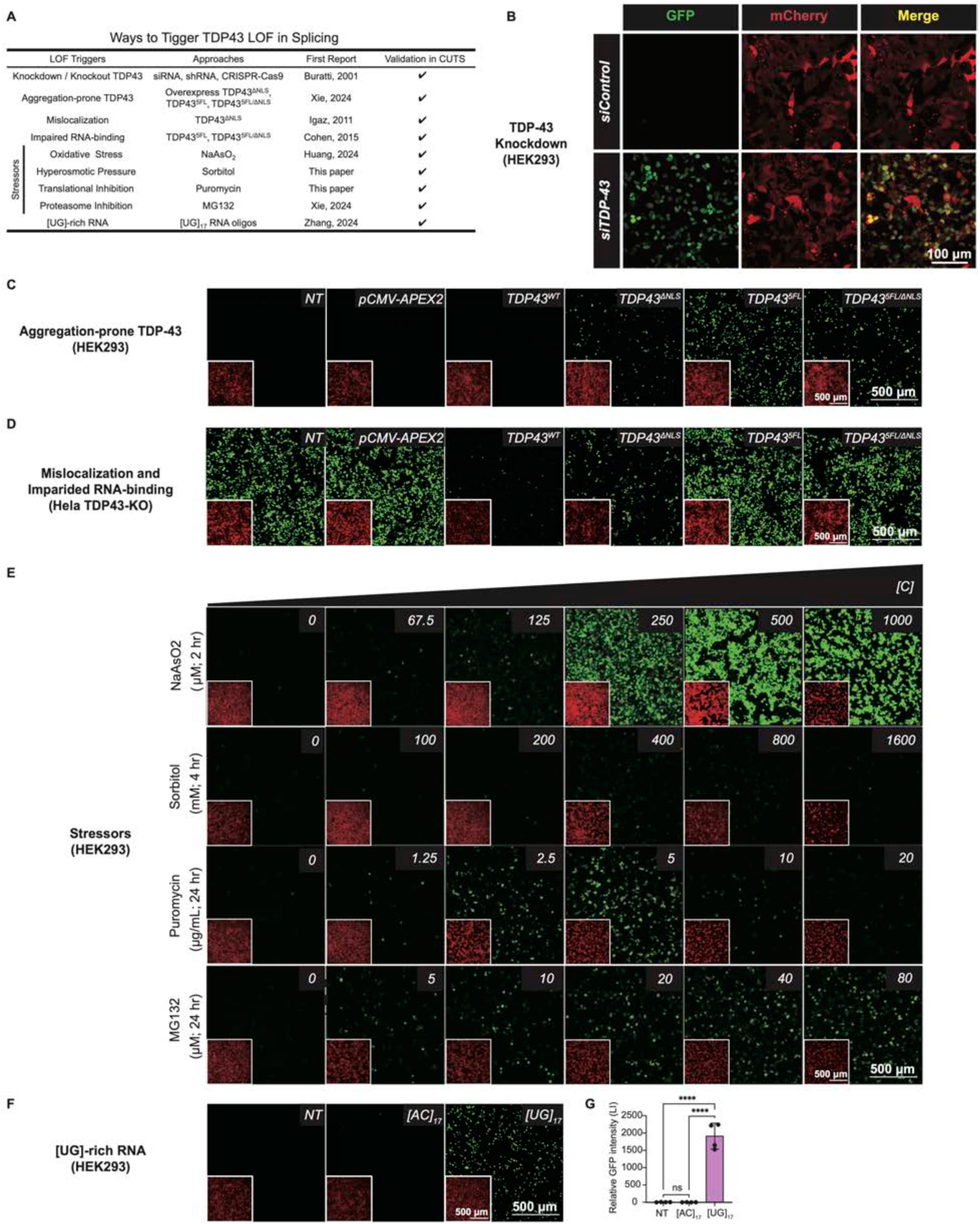
TDP-43’s loss of function in splicing can be triggered by functional mutations, extracellular stressors, and RNA using the CUTS biosensor. **(A)** Systematic summary of TDP-43 loss of function triggers and validations. **(B)** Representative live confocal images of the stable HEK293 cells expressing doxycycline-inducible (72 h of 1 mg/mL doxycycline) CUTS with reverse transfected siRNA control (siControl) (20 nM, 72 h) or TDP-43 (siTDP-43) (20 nM, 72 h). **(C)** Representative live confocal images in stable HEK293 cells expressing doxycycline-inducible (72 h of 1 mg/mL doxycycline) CUTS with the transfection of pCMV-APEX2 backbone, TDP- 43^WT^, TDP-43^ΔNLS^, TDP-43^5FL^, TDP-43^5FL/ΔNLS^, or non-transfected (72 h, N=3 biological replicates). **(D)** Representative live confocal images in Hela TDP-43-KO cells with the co-transfection of doxycycline-inducible (72 h of 1 mg/mL doxycycline) CUTS plasmid and pCMV-APEX2 backbone, TDP-43^WT^, TDP-43^ΔNLS^, TDP-43^5FL^, TDP-43^5FL/ΔNLS^, or no plasmid (72 h, N=3 biological replicates). **(E)** Representative live confocal images in stable HEK293 cells expressing doxycycline-inducible (72 h of 1 mg/mL doxycycline) CUTS with the treatment of NaAsO2 (2 h, gradient doses: 0, 67.5, 125, 250, 500, 1000 μM), Sorbitol (4 h, gradient doses: 0, 100, 200, 400, 800, 1600 mM), Puromycin (24 h, gradient doses: 0, 1.25, 2.5, 5, 10, 20 μg/mL), and MG132 (24 h, gradient doses: 0, 5, 10, 20, 40, 80 μM) (N=2 biological replicates). **(F)** Representative live confocal images in stable HEK293 cells expressing doxycycline-inducible (72 h of 1 mg/mL doxycycline) CUTS with forward transfected [AC]17 and [UG]17 RNA oligos (200 nM, 48 h) For (B) to (F), Green = GFP signal; Red = mCherry signal. Scale bar = 100 or 500 µm. **(G)** Relative GFP fluorescence intensity quantification from live confocal imaging in stable HEK293 cells expressing doxycycline-inducible (72 h of 1 mg/mL doxycycline) CUTS with forward transfected [AC]17 and [UG]17 RNA oligos (200 nM, 48 h). Statistical significance was determined by one-way ANOVA and Tukey’s multiple comparison test (* = P < 0.05; ** = P < 0.01; *** = P < 0.001; **** = P < 0.0001). Data are the mean ± s.d.

**Extended Figure 2:**
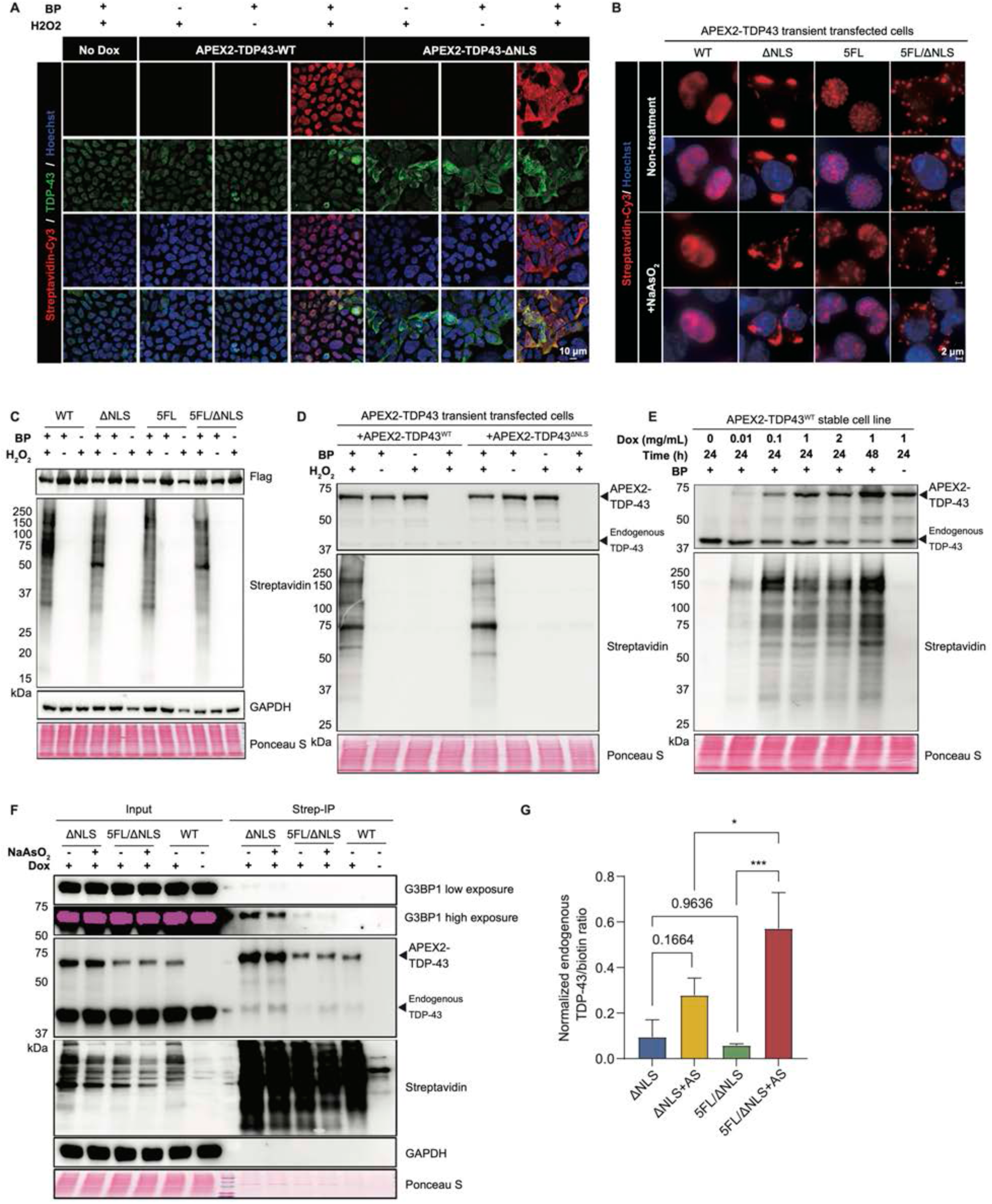
Intrinsic modifications and expression levels cause distinct interactome patterns labeled by APEX2-TDP-43. **(A)** Representative immunofluorescence staining images of TDP-43 and biotin signal labeled by APEX2-TDP-43^WT^ and TDP-43^ΔNLS^ in stable HEK293 cell lines (48 h of 1 mg/mL doxycycline) with or without BP and H2O2. Blue = DAPI; Green = TDP-43; Red = Streptavidin-Cy3. Scale bar = 5 µm. **(B)** Western blot analysis of stable HEK293 cell lines (48 h of 1 mg/mL doxycycline) expressing APEX2-TDP-43^WT^, TDP-43^ΔNLS^, TDP-43^5FL^, and TDP-43^5FL/ΔNLS^ with or without BP and H2O2. **(C)** Western blot analysis of HEK293 cells transfected with pCMV-APEX2-TDP-43^WT^ and TDP-43^ΔNLS^ (24 h) with or without BP and H2O2. **(D)** Western blot analysis of stable HEK293 cell line expressing APEX2-TDP-43^WT^ with different doxycycline doses (0, 0.01, 0.1, 1, 2 mg/mL for 24 h, 1 mg/mL for 48h). **(E)** Western blot analysis of Streptavidin-pull down in stable HEK293 cell lines (48 h of 1 mg/mL doxycycline) expressing APEX2-TDP-43^ΔNLS^, TDP-43^5FL/ΔNLS^, and TDP-43^WT^ with or without NaAsO2 (250 μM, 2 h). **(F)** Relative endogenous TDP-43 (∼43 kDa) protein intensity normalized to total biotin signal from the Streptavidin-pull down in (E) (N = 3 biological replicates). Statistical significance was determined by one-way ANOVA and Tukey’s multiple comparison test (* = P < 0.05; ** = P < 0.01; *** = P < 0.001; **** = P < 0.0001). Data are the mean ± s.d.

**Extended Figure 3:**
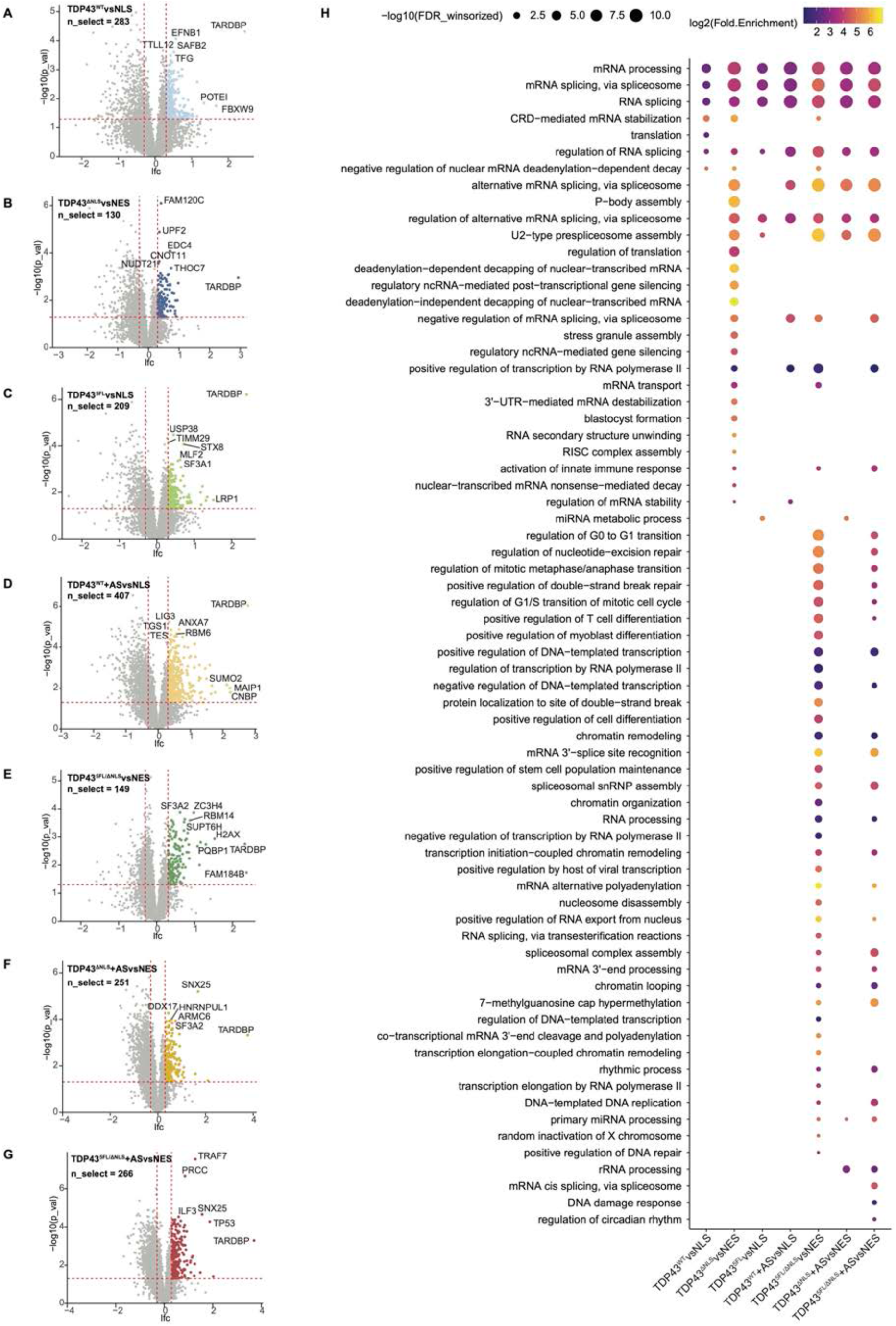
TDP-43 interactomes under different conditions as compared to the nuclear or cytoplasmic background. **(A)** Volcano plot of statistical significance against the log2 fold change (TDP-43^WT^ vs 3XNLS) of protein interactors labeled by the APEX2-TDP-43^WT^ and APEX2-3XNLS as the compartment control (nucleus). Proteins with p value <0.05 and log2FC < -0.3 are regarded as enriched in TDP- 43^WT^ (top-right quadrants). **(B)** Volcano plot of statistical significance against the log2 fold change (TDP-43^ΔNLS^ vs 3XNES) of protein interactors labeled by the APEX2-TDP-43^ΔNLS^ and APEX2-3XNES as the compartment control (cytoplasm). Proteins with p value <0.05 and log2FC < -0.3 are regarded as enriched in TDP-43^ΔNLS^ (top-right quadrants). **(C)** Volcano plot of statistical significance against the log2 fold change (TDP-43^5FL^ vs 3XNLS) of protein interactors labeled by the APEX2-TDP-43^5FL^ and APEX2-3XNLS as the compartment control (nucleus). Proteins with p value <0.05 and log2FC < -0.3 are regarded as enriched in TDP- 43^5FL^ (top-right quadrants). **(D)** Volcano plot of statistical significance against the log2 fold change (TDP-43^WT^+AS vs 3XNLS) of protein interactors labeled by the APEX2-TDP-43^WT^ with NaAsO2 treatment and APEX2- 3XNLS as the compartment control (nucleus). Proteins with p value <0.05 and log2FC < -0.3 are regarded as enriched in TDP-43^WT^+AS (top-right quadrants). **(E)** Volcano plot of statistical significance against the log2 fold change (TDP-43^5FL/ΔNLS^ vs 3XNES) of protein interactors labeled by the APEX2-TDP-43^5FL/ΔNLS^ and APEX2-3XNES as the compartment control (cytoplasm). Proteins with p value <0.05 and log2FC < -0.3 are regarded as enriched in TDP-43^5FL/ΔNLS^ (top-right quadrants). **(F)** Volcano plot of statistical significance against the log2 fold change (TDP-43^ΔNLS^+AS vs 3XNES) of protein interactors labeled by the APEX2-TDP-43^ΔNLS^ with NaAsO2 treatment and APEX2-3XNES as the compartment control (cytoplasm). Proteins with p value <0.05 and log2FC < -0.3 are regarded as enriched in TDP-43^ΔNLS^+AS (top-right quadrants). **(G)** Volcano plot of statistical significance against the log2 fold change (TDP-43^5FL/ΔNLS^ +AS vs 3XNES) of protein interactors labeled by the APEX2-TDP-43^5FL/ΔNLS^ with NaAsO2 treatment and APEX2-3XNES as the compartment control (cytoplasm). Proteins with p value <0.05 and log2FC < -0.3 are regarded as enriched in TDP-43^5FL/ΔNLS^ +AS (top-right quadrants). For (A) to (G), The p values were calculated with two-sided Student’s t-test. **(H)** Gene ontology (GO) term analysis of proteins enriched in TDP-43^WT^, , TDP-43^ΔNLS^, TDP- 43^5FL^, TDP-43^WT^+AS, TDP-43^5FL/ΔNLS^, TDP-43^ΔNLS^+AS, and TDP-43^5FL/ΔNLS^+AS groups. FDR was calculated based on adaptive linear step-up adjusted p-values.

**Extended Figure 4:**
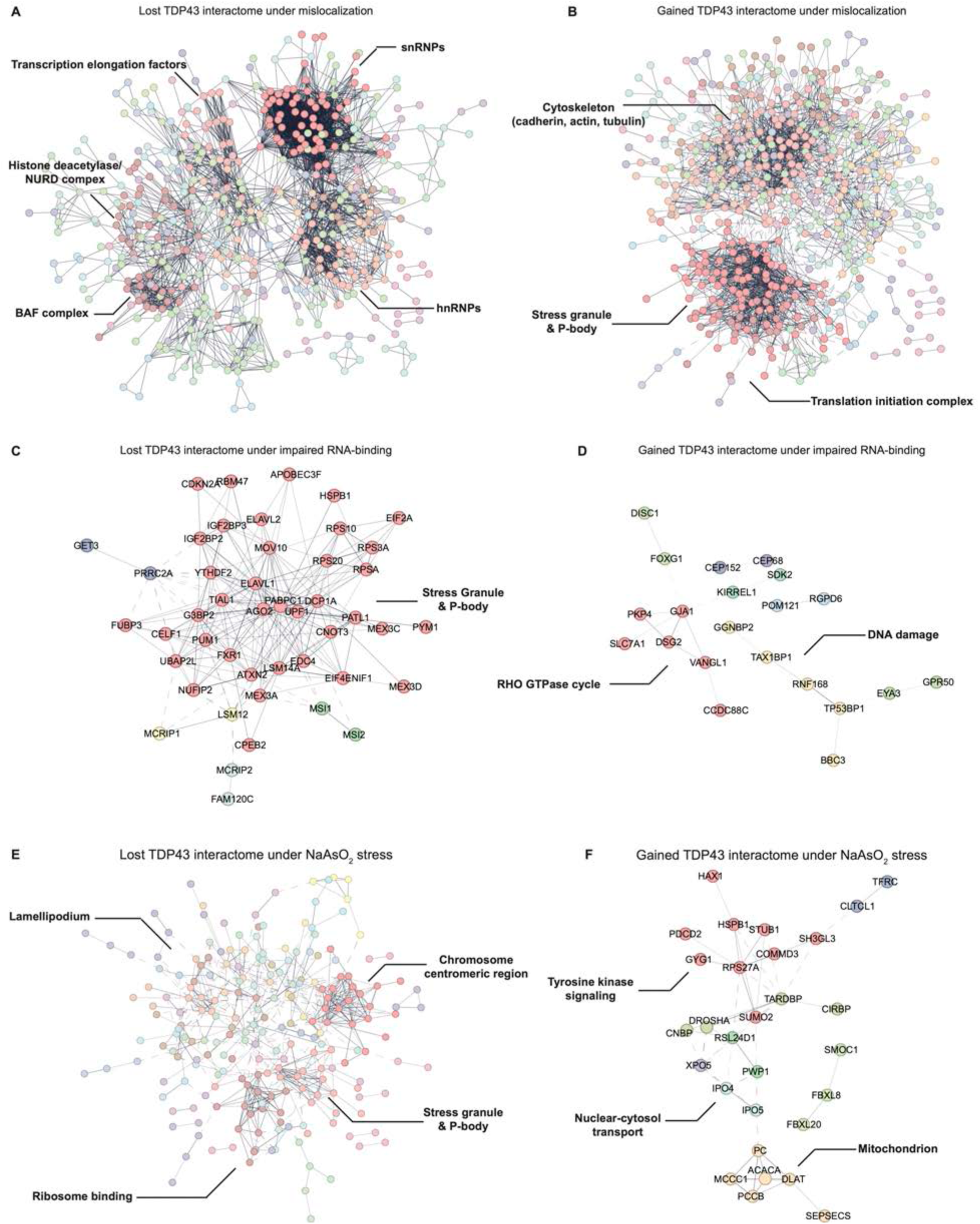
TDP-43’s interactome network is significantly changed under mislocalization, impaired RNA binding, and NaAsO2 stress, showing by lost or gained interaction with functional protein clusters. **(A)** Protein network and MCL clustering by STRING database of significantly lost TDP-43 interacting protein under mislocalization. Proteins with at least 2 times significantly decreased (q value < 0.05, log2FC < 1) in the direct comparisons of TDP-43^ΔNLS^ vs TDP-43^WT^, TDP-43^5FL/ΔNLS^ vs TDP-43^5FL^, and TDP-43^ΔNLS^+AS vs TDP-43^WT^+AS are included. **(B)** Protein network and MCL clustering by STRING database of significantly gained TDP-43 interacting protein under mislocalization. Proteins with at least 2 times significantly increased (q value < 0.05, log2FC > 1) in the direct comparisons of TDP-43^ΔNLS^ vs TDP-43^WT^, TDP-43^5FL/ΔNLS^ vs TDP-43^5FL^, and TDP-43^ΔNLS^+AS vs TDP-43^WT^+AS are included. **(C)** Protein network and MCL clustering by STRING database of significantly lost TDP-43 interacting protein under impaired RNA-binding. Proteins with at least 2 times significantly decreased (p value < 0.05, log2FC < 0.3) in the direct comparisons of TDP-43^5FL^ vs TDP-43^WT^, TDP-43^5FL/ΔNLS^ vs TDP-43^ΔNLS^, and TDP-43^5FL/ΔNLS^+AS vs TDP-43^ΔNLS^+AS are included. **(D)** Protein network and MCL clustering by STRING database of significantly gained TDP-43 interacting protein under impaired RNA-binding. Proteins with at least 2 times significantly increased (p value < 0.05, log2FC < 0.3) in the direct comparisons of TDP-43^5FL^ vs TDP-43^WT^, TDP-43^5FL/ΔNLS^ vs TDP-43^ΔNLS^, and TDP-43^5FL/ΔNLS^+AS vs TDP-43^ΔNLS^+AS are included. **(E)** Protein network and MCL clustering by STRING database of significantly lost TDP-43 interacting protein under NaAsO2 stress. Proteins with at least 2 times significantly decreased (p value < 0.05, log2FC < 0.3) in the direct comparisons of TDP-43^WT^+AS vs TDP-43^WT^, TDP- 43^ΔNLS^+AS vs TDP-43^ΔNLS^, and TDP-43^5FL/ΔNLS^+AS vs TDP-43^5FL/ΔNLS^ are included. **(F)** Protein network and MCL clustering by STRING database of significantly gained TDP-43 interacting protein under NaAsO2 stress. Proteins with at least 2 times significantly increased (p value < 0.05, log2FC < 0.3) in the direct comparisons of TDP-43^WT^+AS vs TDP-43^WT^, TDP- 43^ΔNLS^+AS vs TDP-43^ΔNLS^, and TDP-43^5FL/ΔNLS^+AS vs TDP-43^5FL/ΔNLS^ are included.

**Extended Figure 5:**
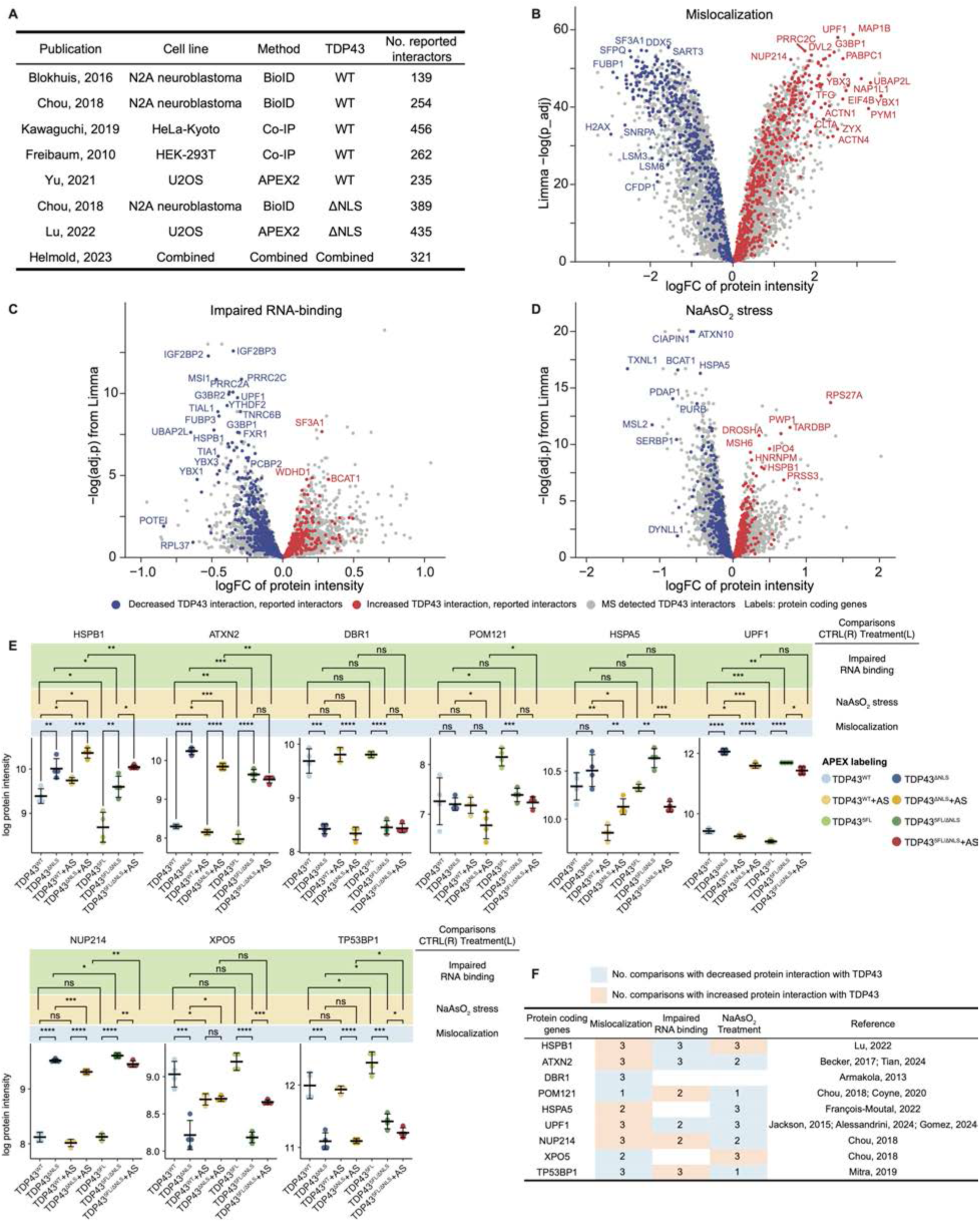
APEX2 proximity labeling identifies reported proteins and features associated with TDP-43 pathobiology. **(A)** Summary table of previously reported TDP-43 interactome datasets, including the references, model cell lines, methods for mapping interactome, TDP-43 variation (WT or ΔNLS), and reported interactor numbers. **(B)** Volcano plot of statistical significance against the log2 fold change of protein interactors under mislocalization labeled by APEX2-TDP-43 variants. This plot combines all the three different direct comparisons (TDP-43^ΔNLS^ vs TDP-43^WT^, TDP-43^5FL/ΔNLS^ vs TDP-43^5FL^, and TDP- 43^ΔNLS^+AS vs TDP-43^WT^+AS) related to mislocalization. The adjusted p values were calculated using two-sided Student’s t-test. Protein interactors reported by previous literature are labeled either increased (red) or decreased (blue). **(C)** Volcano plot of statistical significance against the log2 fold change of protein interactors under impaired RNA-binding labeled by APEX2-TDP-43 variants. This plot combines all the three different direct comparisons (TDP-43^5FL^ vs TDP-43^WT^, TDP-43^5FL/ΔNLS^ vs TDP-43^ΔNLS^, and TDP- 43^5FL/ΔNLS^+AS vs TDP-43^ΔNLS^+AS) related to impaired RNA-binding. The p values were calculated using two-sided Student’s t-test. Protein interactors reported by previous literature are labeled either increased (red) or decreased (blue). **(D)** Volcano plot of statistical significance against the log2 fold change of protein interactors under NaAsO2 stress labeled by APEX2-TDP-43 variants. This plot combines all the three different direct comparisons (TDP-43^WT^+AS vs TDP-43^WT^, TDP-43^ΔNLS^+AS vs TDP-43^ΔNLS^, and TDP- 43^5FL/ΔNLS^+AS vs TDP-43^5FL/ΔNLS^) related to NaAsO2 stress. The p values were calculated using two-sided Student’s t-test. Protein interactors reported by previous literature are labeled either increased (red) or decreased (blue). **(E)** Scatter plots of log protein intensity in the seven settings of representative previously reported protein modifiers of TDP-43 pathology. HSPB1, ATXN2, DBR1, POM121, HSPA5, UPF1, NUP214, XPO5, and TP53BP1 are included. **(F)** Summary table of representative previously reported protein modifiers of TDP-43 pathology and their changing patterns in our APEX2-TDP-43 proximity labeling.

**Extended Figure 6:**
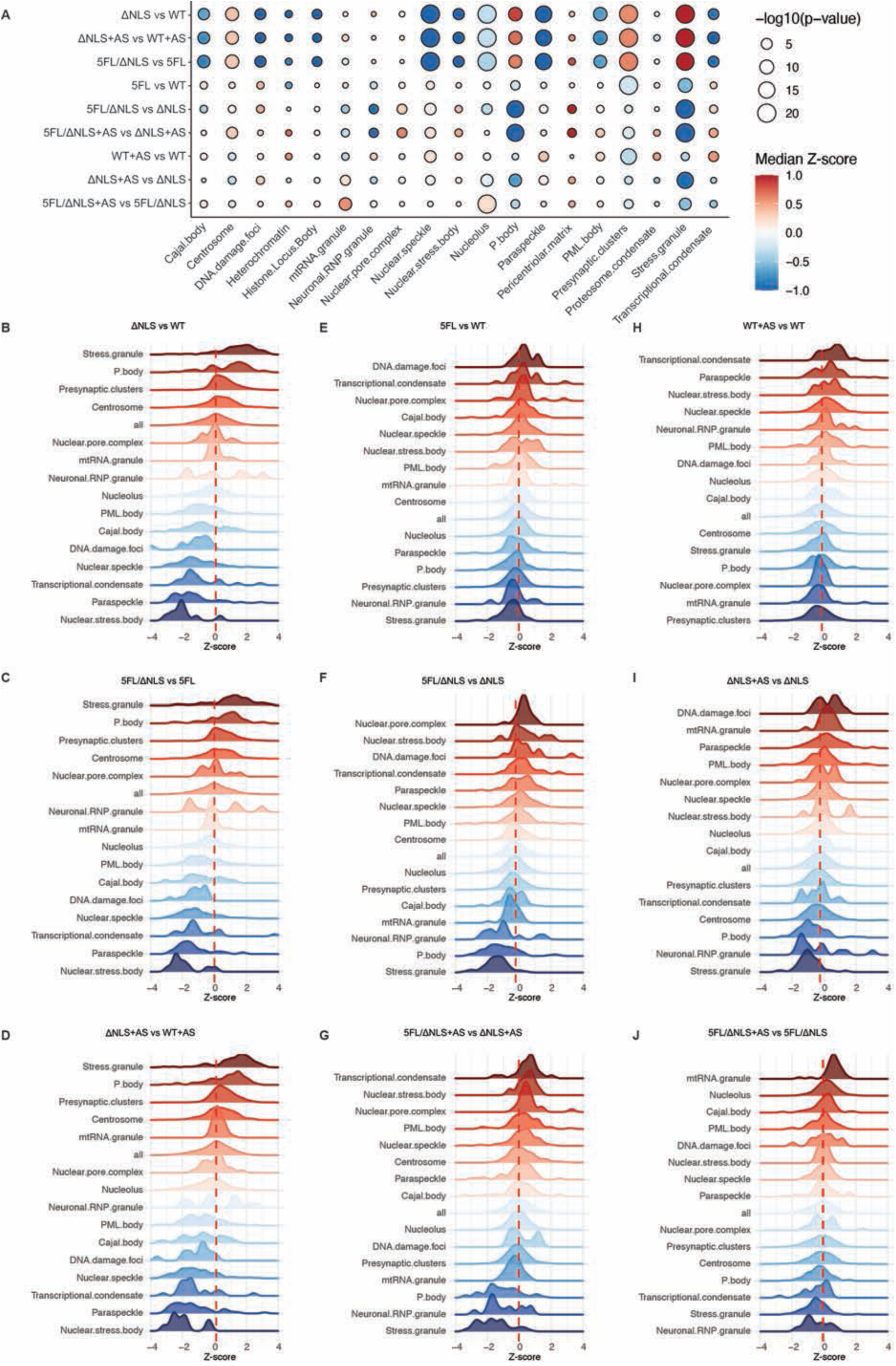
Pathological factors tune TDP-43 to shift among biomolecular condensates. **(A) Deviation** dot plot of all the evaluated BMCs from the nine pathological comparisons. Dot color represents normalized mean deviation value; Dot size represents p value. The p values were calculated using Wilcoxon signed-rank test. **(B)** Ranked distribution density plots of different BMCs’ proteins and all proteins in the comparison of TDP-43^ΔNLS^ vs TDP-43^WT^. **(C)** Ranked distribution density plots of different BMCs’ proteins and all proteins in the comparison of TDP-43^5FL^ vs TDP-43^WT^. **(D)** Ranked distribution density plots of different BMCs’ proteins and all proteins in the comparison of TDP-43^WT^+AS vs TDP-43^WT^. **(E)** Ranked distribution density plots of different BMCs’ proteins and all proteins in the comparison of TDP-43^5FL/ΔNLS^ vs TDP-43^5FL^. **(F)** Ranked distribution density plots of different BMCs’ proteins and all proteins in the comparison of TDP-43^5FL/ΔNLS^ vs TDP-43^ΔNLS^. **(G)** Ranked distribution density plots of different BMCs’ proteins and all proteins in the comparison of TDP-43^ΔNLS^+AS vs TDP-43^ΔNLS^. **(H)** Ranked distribution density plots of different BMCs’ proteins and all proteins in the comparison of TDP-43^ΔNLS^+AS vs TDP-43^WT^+AS. **(I)** Ranked distribution density plots of different BMCs’ proteins and all proteins in the comparison of TDP-43^5FL/ΔNLS^+AS vs TDP-43^ΔNLS^+AS. **(J)** Ranked distribution density plots of different BMCs’ proteins and all proteins in the comparison of TDP-43^5FL/ΔNLS^+AS vs TDP-43^5FL/ΔNLS^.

**Extended Figure 7:**
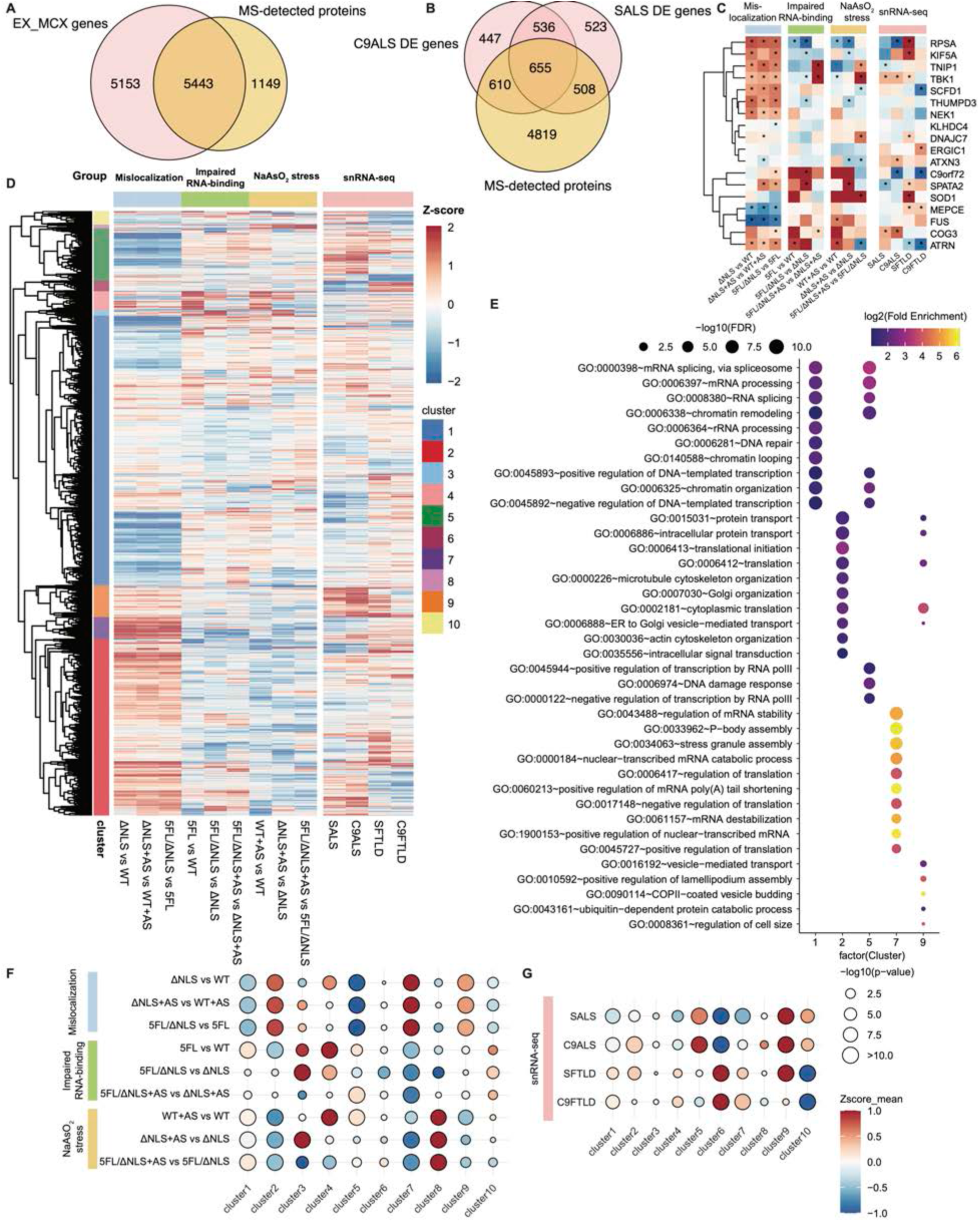
Context-dependent TDP-43 interactors are altered in snRNA-seq ALS/FTLD patient datasets. **(A)** Venn diagram between all the genes of the detected proteins by mass spectrum (MS) in this study (right) and all the genes detected by snRNA-seq from the motor cortex excitatory neurons of all patients (left). **(B)** Venn diagram between all the genes of the detected proteins by mass spectrum in this study (bottom) and differentially-expressed (DE) genes found by snRNA-seq from the motor cortex excitatory neurons of C9-ALS (top left) and sALS (top right) patients. **(C)** Heatmap of ALS/FTLD GWAS-associated genes in the nine pathological comparisons of proximity labeling and snRNA-seq of the motor cortex excitatory neurons of ALS or FTLD patients. Cell color represents normalized mean z-values with “*” labels p value < 0.05. The unadjusted p values were calculated using two-sided Student’s t-test. **(D)** Heatmap of the gene clusters acquired from the combination of APEX2 proximity labeling with the nine pathological comparisons and snRNA-seq comparisons of the excitatory neurons in the motor cortex between ALS or FTLD patients and controls. Clustering is performed by hierarchical clustering method. Cell color represents normalized mean z-values. **(E)** Gene ontology (GO) term analysis of genes enriched from each gene clusters in (D), showing GO terms with FDR < 0.05. FDR was calculated based on adaptive linear step-up adjusted p-values. **(F)** Deviation dot plot of all the gene clusters in (D) in the nine pathological comparisons. **(G)** Deviation dot plot of all the gene clusters in (D) in the snRNA-seq comparisons of the excitatory neurons in the motor cortex between ALS or FTLD patients and controls. For (F) and (G), dot color represents normalized mean deviation value; Dot size represents p value. The p values were calculated using Wilcoxon signed-rank test.

**Extended Figure 8:**
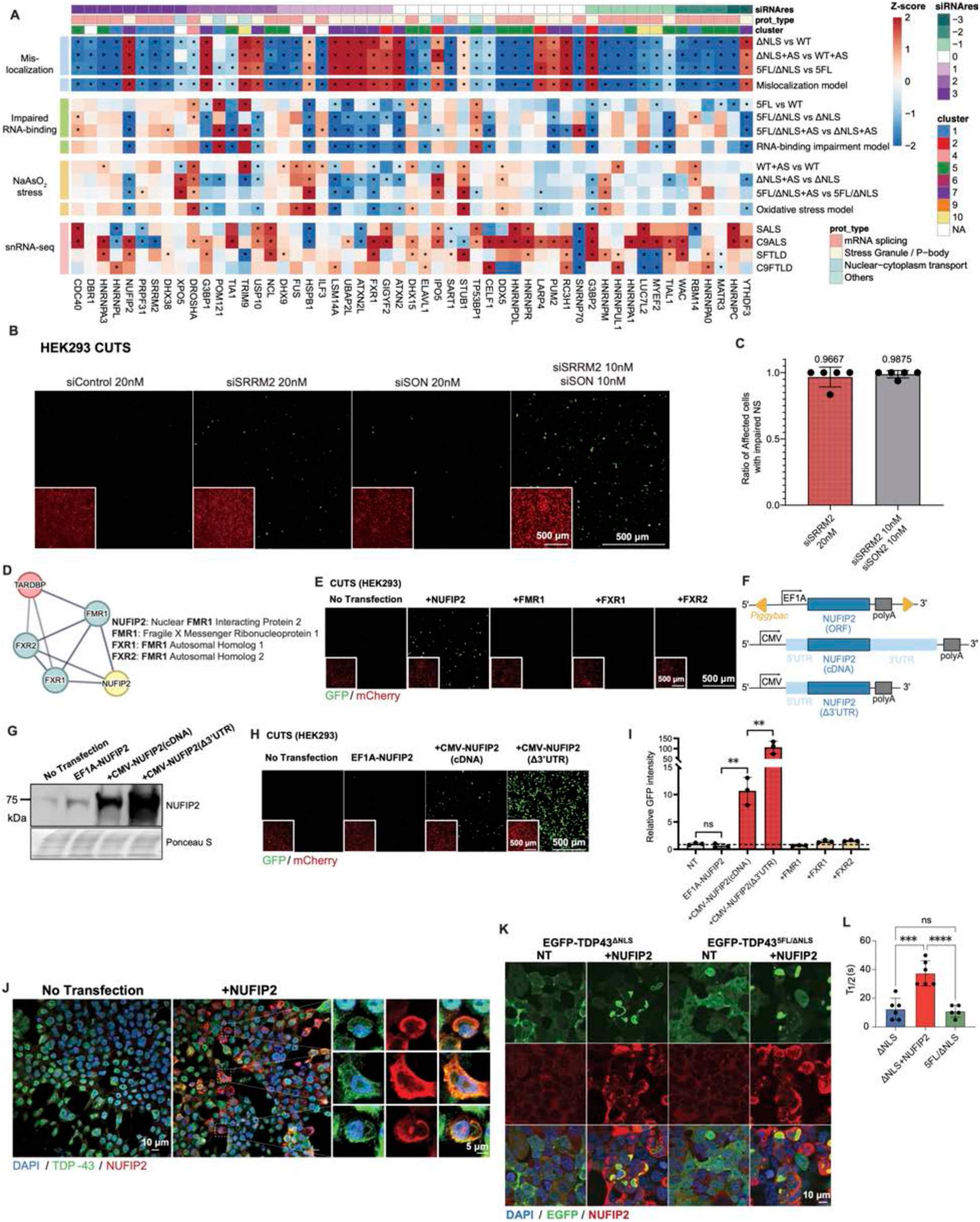
TDP-43 interactors trigger its loss of function with distinct mechanisms. (A) Heatmap of the ranked preliminary CUTS siRNA screening result (N = 3 biological replicates) of 53 selected proteins mostly from Cluster 1, 5, and 7 of three categories (mRNA splicing, stress granule or P-body, and nuclear-cytoplasm transport). Cell color shows the summary of the results from each replicate with whether increasing GFP signal (causing LOF, +1) or lowering GFP signal (rescuing LOF, -1) comparing to control siRNA (siControl). Heatmap also demonstrates the data of the 53 selected proteins in the nine pathological comparisons of proximity labeling and snRNA-seq comparisons of the excitatory neurons in the motor cortex between ALS or FTLD patients and controls. Cell color represents normalized mean z-values with “*” labels p value < 0.05. The unadjusted p values were calculated using a two-sided Student’s t-test. (B) Representative live confocal images of the stable HEK293 cells expressing doxycycline-inducible (72 h of 1 mg/mL doxycycline) CUTS with reverse transfected siRNA control (siControl) (20 nM, 96h), SRRM2 (siSRRM2) (20 nM, 96 h), SON (siSON) (20 nM, 96 h), or siSRRM2 + siSON (10 nM each, 96 h). (C) Ratio of the HEK293 cells with impaired nuclear speckles (no SC-35 condensates) in all of the abnormal cells (with nuclear-aggregated and cytoplasmic mislocalized EGFP-TDP-43^WT^) expressing EGFP-TDP-43^WT^ under siSRRM2 (20 nM, 96 h) or siSRRM2 + siSON (10 nM each, 96 h) (N=5 replicates with each >230 cells). (D) Protein network graph by STRING database of TDP-43 and NUFIP2-related proteins, including FMR1, FXR1, and FXR2. (E) Representative live confocal images of the stable HEK293 cells expressing doxycycline-inducible (72 h of 1 mg/mL doxycycline) CUTS with no transfection or transfection of NUFIP2 cDNA plasmid and FMR1, FXR1, and FXR2 open reading frame (ORF) plasmids (500 ng, 72 h). (F) Schematic diagram of the plasmids design to overexpress NUFIP2 at different levels. **EF1A- NUFIP2** (top) enables constitutive expression (stable cell line) of NUFIP2 ORF driven by EF1A promoter through *Piggybac* integration; **CMV-NUFIP2(cDNA)** (middle) produces transient expression of full length NUFIP2 cDNA upon transfection; **CMV-NUFIP2(Δ3’UTR)** (bottom) was derived from CMV-NUFIP2(cDNA) and has the 3’UTR removed. (G) Western blot analysis of NUFIP2 protein level in HEK293 cells under no transfection, EF1A- NUFIP2 (stable cell line), CMV-NUFIP2(cDNA), and CMV-NUFIP2(Δ3’UTR) (transfected with 500 ng, 72 h). (H) Representative live confocal images of the stable HEK293 cells expressing doxycycline-inducible (72 h of 1 mg/mL doxycycline) CUTS under no transfection, EF1A-NUFIP2 (stable cell line), CMV-NUFIP2(cDNA), and CMV-NUFIP2(Δ3’UTR) (transfected with 500 ng, 72 h). (I) Relative GFP fluorescence intensity quantification from live confocal imaging of (F) and (H). Statistical significance was determined by a two-sided Student’s t-test (* = P < 0.05; ** = P < 0.01; *** = P < 0.001; **** = P < 0.0001). Data are the mean ± s.d. (J) Representative immunofluorescence staining images of wildtype HEK293 cells with or without NUFIP2 overexpression. Magnified views (right) of the region with dashed white squares in the main image (left) represent examples of cytoplasmic “shell-like” structure by NUFIP2 and endogenous TDP-43. Blue = Hoechst; Green = TDP-43; Red = NUFIP2. Scale bar = 5 µm. (K) Representative immunofluorescence staining images of stable HEK293 cells expressing EGFP- TDP-43^ΔNLS^ and EGFP- TDP-43^5FL/ΔNLS^ with or without the transfection of NUFIP2 overexpression. Blue = Hoechst; Green = EGFP-TDP-43; Red = NUFIP2; Scale bar = 5 µm. (L) Half-recovery time (T1/2) analysis of the FRAP experiment of NUFIP2 overexpression on cytoplasmic TDP-43 with (EGFP- TDP-43^ΔNLS^) or without (EGFP- TDP-43^5FL/ΔNLS^) impaired RNA-binding. Statistical significance was determined by a two-sided Student’s t-test (* = P < 0.05; ** = P < 0.01; *** = P < 0.001; **** = P < 0.0001). Data are the mean ± s.d.

**Extended Figure 9:**
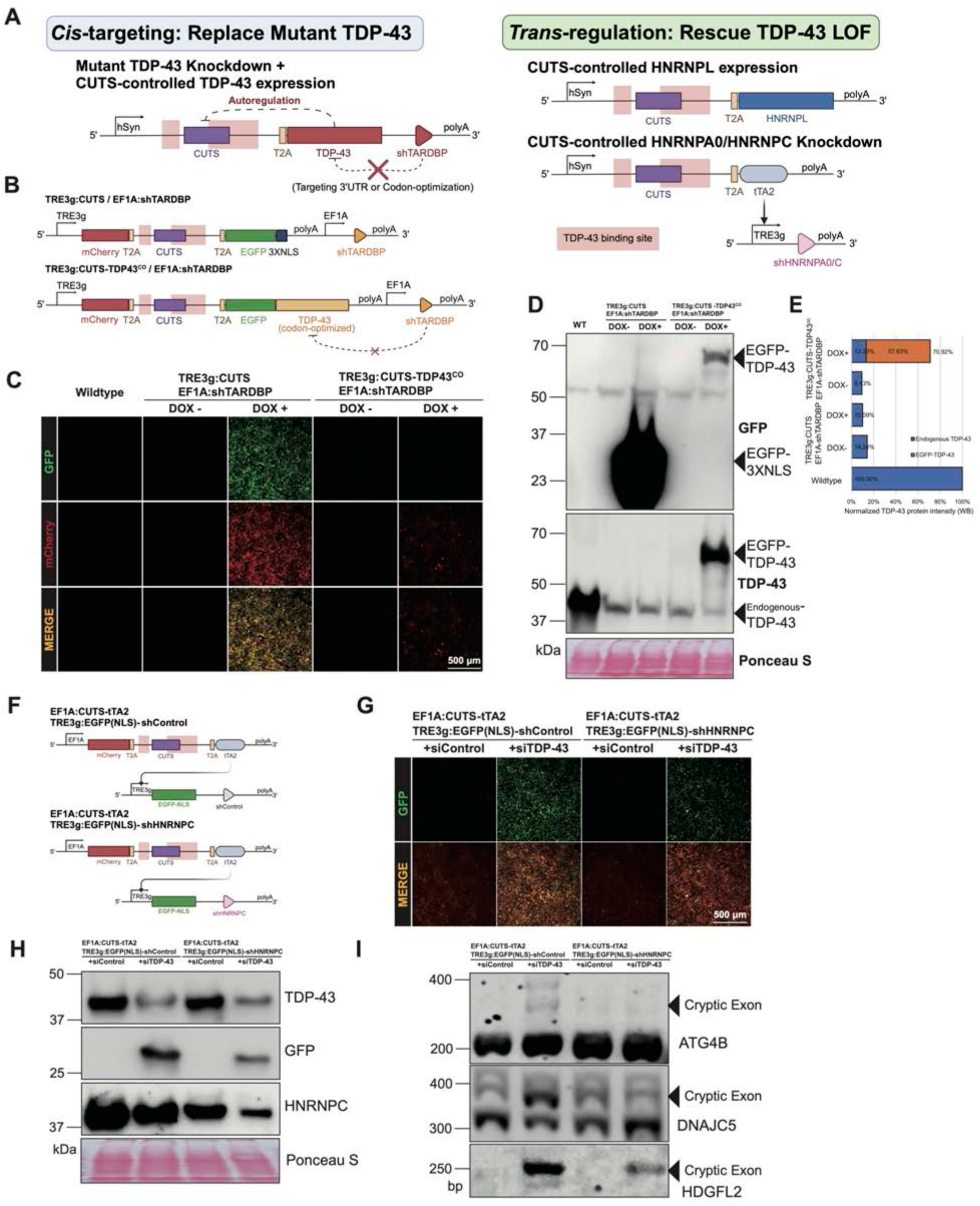
Modulating TDP-43 and interactors to rescue cryptic exon splicing. **(A)** Schematic diagram of two novel rescue approaches rescuing TDP-43 LOF by *cis*-targeting TDP-43 or *trans*-regulating TDP-43’s interactors. **(B)** Schematic diagram of the plasmids design to generate stable HEK293 cell lines with TDP-43 knockdown-replacement (TKR) strategy. Endogenous TDP-43 was knocked down by shRNA and replaced with doxycycline-inducible CUTS-controlled expression of codon-optimized EGFP- TDP-43. **(C)** Representative live confocal images of wildtype HEK293 or the stable cell lines expressing TKR or control systems in (B) with or without doxycycline (96 h of 1 mg/mL doxycycline). Green = GFP signal; Red = mCherry signal. Scale bar = 100 µm. **(D)** Western blot analysis of the cells in (C). **(E)** Percentage of endogenous or exogenous TDP-43 protein levels normalized to wildtype control from western blot analysis in (D). **(F)** Schematic diagram of the plasmid design to generate stable HEK293 cell lines with CUTS- controlled HNRNPC knockdown (CCK) rescue. EGFP-NLS-coupled shRNA targeting HNRNPC is controlled by CUTS through the Tet-On 3G system (TRE3g) to enable LOF-specific activation. **(G)** Representative live confocal images of wildtype HEK293 or the stable cell lines expressing CUTS-shHNRNPC or CUTS-shControl systems in (F) with reverse transfected siControl or siTDP-43 (20 nM, 7 days). Green = GFP signal; Red = mCherry signal. Scale bar = 100 µm. **(H)** Western blot analysis of the cells in (G). **(I)** Representative image of RT–PCR products from TDP-43-regulated endogenous cryptic exon targets (ATG4B, DNAJC5, and HDGFL2) in stable HEK293 cells from (H). Images represent similar results in two independent experiments.

**Extended Figure 10:**
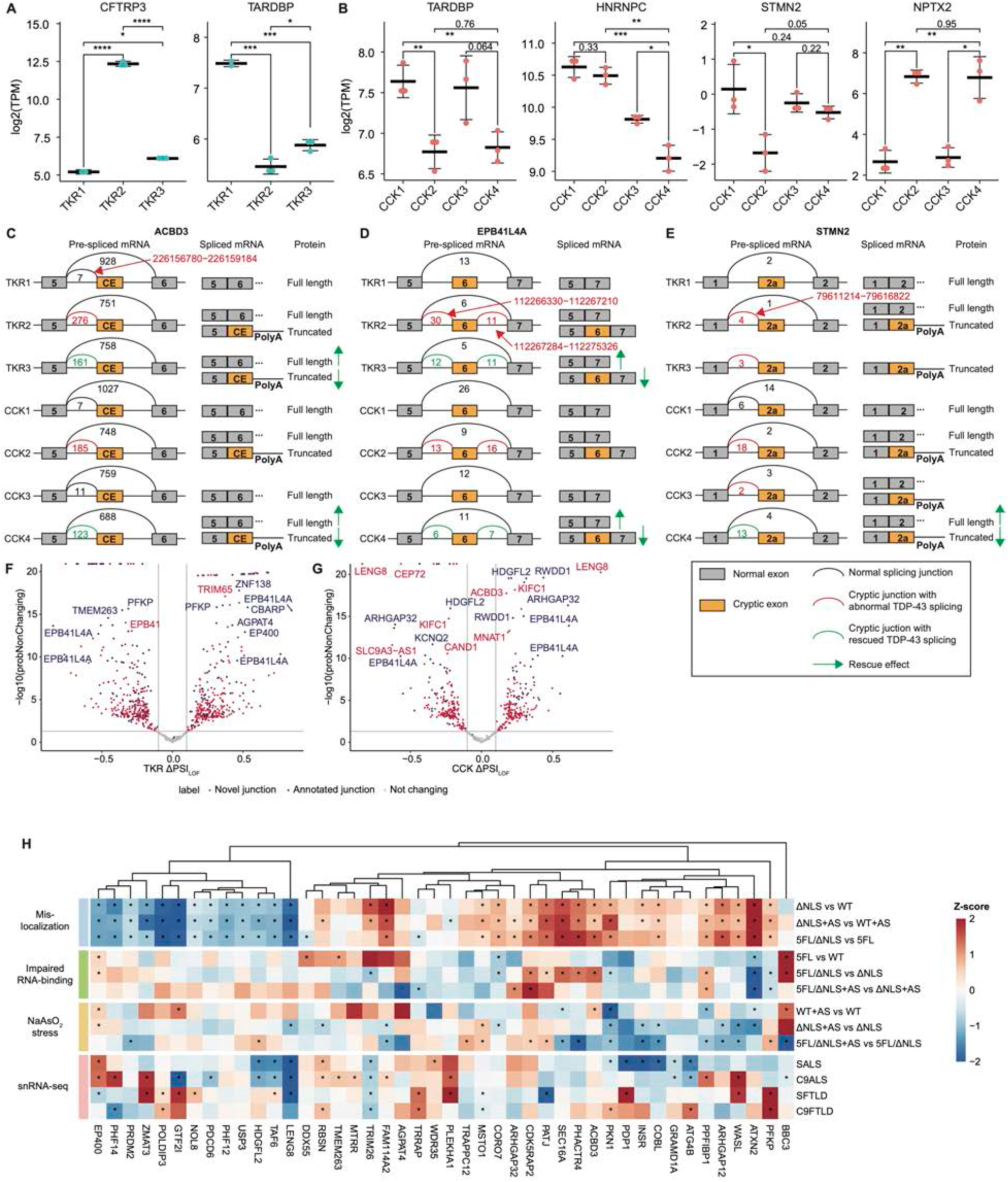
TDP-43 knockdown-replacement (TKR) and CUTS-controlled HNRNPC knockdown (CCK) exhibit distinct transcriptomic features. **(A)** Transcripts Per Million (TPM) analysis of CFTRP3 (CFTR Pseudogene 3) and TARDBP (TDP-43) mRNA levels in the TKR groups. CFTRP3 serves as a good evaluation of the CUTS activation level as it annotates the isoform of CFTR with Exon 9 inclusion, which is a cryptic exon regulated by TDP-43 and was used to generate CUTS. TARDBP’s TPM is slightly up-regulated in TKR3 than TKR2, because the codon-optimized TDP-43 still processed a ∼40bp identical sequence as wildtype TDP-43 on the 5’ end. **(B)** TPM analysis of TARDBP, HNRNPC, STMN2, and NPTX2 mRNA levels in CCK groups. For (A) and (B), statistical significance was determined by two-sided Student’s t-test with p values labeled (* = P < 0.05; ** = P < 0.01; *** = P < 0.001; **** = P < 0.0001). Data are the mean ± s.d. **(C)** Representative splicing plots of the cryptic exon (CE) in ACBD3 in TKR and CCK groups by Voila. The CE inclusion of ACBD3 is predicted to create an alternative polyadenylation event. **(D)** Representative splicing plots of the cryptic exon (Exon 6) in EPB41L4A in TKR and CCK groups by Voila. **(E)** Representative splicing plots of the cryptic exon (Exon 2a) in STMN2 in TKR and CCK groups by Voila. The CE inclusion of STMN was reported to create an alternative polyadenylation event. **(F)** Volcano plot of the probability of non-changing against the ΔPSI of the differential splicing analysis by MAJIQ between baseline control (TKR1, N=2) and TDP-43 knockdown positive control (TKR2, N=3) in the TKR systems. Each point shows a specific splicing junction with or without (novel) annotation from a previous study in an iPSC-derived model. **(G)** Volcano plot of the probability of non-changing against the ΔPSI of the differential splicing analysis by MAJIQ between baseline control (CCK1, N=3) and TDP-43 knockdown positive control (CCK2, N=3) in the CCK systems. Each point shows a specific splicing junction with or without (novel) annotation from a previous study in an iPSC-derived model. **(H)** Heatmap of TDP-43-regulated genes (from the 198 selected indexes in splicing analysis) in the nine pathological comparisons of proximity labeling and snRNA-seq comparisons of the excitatory neurons in the motor cortex between ALS or FTLD patients and controls. Cell color represents normalized mean z-values with “*” labels p value < 0.05. The unadjusted p values were calculated using a two-sided Student’s t-test.

